# Mapping the evolution of T cell states during response and resistance to adoptive cellular therapy

**DOI:** 10.1101/2020.07.08.194332

**Authors:** Pavan Bachireddy, Elham Azizi, Cassandra Burdziak, Vinhkhang N Nguyen, Christina Ennis, Zi-Ning Choo, Shuqiang Li, Kenneth J. Livak, Donna S Neuberg, Robert J Soiffer, Jerome Ritz, Edwin P Alyea, Dana Pe’er, Catherine J Wu

**Affiliations:** Broad Institute of MIT and Harvard, Cambridge, MA, USA; Department of Medical Oncology, Dana-Farber Cancer Institute, Boston, MA, USA; Harvard Medical School, Boston, MA, USA; Computational and Systems Biology Program, Sloan Kettering Institute, Memorial Sloan Kettering Cancer Center, New York, NY, USA; Department of Biomedical Engineering and Irving Institute for Cancer Dynamics, Columbia University, New York, NY, USA; Translational Immunogenomics Laboratory, Dana-Farber Cancer Institute, Boston, MA, USA; Department of Data Sciences, Dana-Farber Cancer Institute, Boston, MA, USA; Parker Institute of Immunotherapy, Memorial Sloan Kettering Cancer Center, New York, NY, USA

**Keywords:** immunotherapy, donor lymphocyte infusion, leukemia, scRNA-seq, response, resistance, probabilistic models, statistical machine learning, T cell exhaustion, scTCR-seq, ATAC-seq, gene regulatory networks, allogeneic hematopoietic stem cell transplant

## Abstract

Immune therapies have transformed the cancer therapeutic landscape but fail to benefit most patients. To elucidate the underlying mechanisms by which T cells mediate elimination of leukemia, we generated a high-resolution map of longitudinal T cell dynamics within the same tumor microenvironment (TME) during response or resistance to donor lymphocyte infusion (DLI), a widely used immunotherapy for relapsed leukemia. We analyzed 87,939 bone marrow-derived single T cell transcriptomes, along with chromatin accessibility and single T cell receptor clonality profiles, by developing novel machine learning tools for integrating longitudinal and multimodal data. We found that pre-treatment enrichment and post-treatment rapid, durable expansion of ‘terminal’ (T_EX_) and ‘precursor’ (T_PEX_) exhausted subsets, respectively, defined DLI response. A contrasting, heterogeneous pattern of T cell dysfunction marked DLI resistance. Unexpectedly, T_PEX_ cells that expanded in responders did not arise from the infusion product but instead from both pre-existing and novel clonotypes recruited to the TME. Our unbiased dissection of the TME using a Bayesian method, Symphony, defined the T cell circuitry underlying effective human anti-leukemic immune responses that may be broadly relevant to other exhaustion antagonists across cancers. Finally, we provide a general analysis paradigm for exploiting temporal single-cell genomic profiling for deep understanding of therapeutic scenarios beyond oncology.

---

Despite the potency of cancer immunotherapy for a subset of cancer patients, the variability in responses and efficacy suggests that the fundamental mechanisms, cell types and pathways driving clinical outcomes remain elusive^1^. Single-cell transcriptomic profiling is a powerful technology that can characterize the full range of immune cell states and gene programs in the tumor microenvironment (TME) in a comprehensive and unbiased manner. Studying the evolution of the TME at single-cell resolution before and after therapy can thus reveal how heterogeneous cell states evolve in relation to distinct clinical outcomes and illuminate the molecular and cellular determinants of immunotherapeutic response or resistance^1,2^. However, high-resolution studies of such temporal dynamics are typically performed in animal model systems^3^ due to confounding factors and logistical challenges, and they may not fully capture the response of tumors in patients.

To overcome challenges in human clinical studies, we leveraged a well-annotated longitudinal cohort of patients treated with DLI, an established adoptive cellular therapy for relapsed leukemia after allogeneic hematopoietic stem cell transplant (allo-SCT). The clear, binary outcomes of response or resistance; the clinical samples collected over a multi-year time-span; and the lack of confounding chemotherapy or immunomodulators has made DLI therapy an attractive immunotherapeutic setting to study the essential ‘search and destroy’ functions of donor-derived T cell responses that underlie the therapeutic graft-versus-leukemia (GvL) effect of allo-SCT^4,5^. Over the last 30 years, DLI has directly demonstrated the potency of GvL by inducing durable molecular remissions in ~75% of patients with relapsed chronic myelogenous leukemia (CML) following allo-SCT, in the absence of further chemo- or radiotherapy^6,7^.

Response to DLI modified by CD8-depletion has been associated with decreased toxicity^8–11^, increased T cell receptor (TCR) repertoire diversity^12^, expansion of endogenous, tumor-specific, marrow resident CD8+ T cells^13^, and reversal of T cell exhaustion^14^. Similar observations in acute myelogenous leukemia^15^ suggest that the study of DLI in CML can reveal insights that are broadly relevant across hematologic malignancies. Yet despite the widespread use of DLI for the treatment of relapsed disease following allo-SCT^6,16^, the mechanistic basis for its effectiveness remains incompletely understood. Such insight would elucidate the pathways driving GvL clinical outcomes and inform therapeutic strategies to prevent or treat relapse following allo-SCT.

To elucidate the T cell subsets mediating DLI resistance, response and exhaustion after DLI therapy, we analyze single-cell T cell transcriptomes, bulk chromatin accessibility profiles, cluster-specific gene regulatory networks and single T cell clonality data from bone marrow biopsies of a longitudinal cohort of patients with relapsed CML after allo-SCT treated with DLI^10^. We introduce new computational models to integrate data across multiple timepoints and modalities and use this unbiased framework to reveal the subsets of exhausted T cells whose enrichment and divergent dynamics define immunotherapeutic responses in human leukemia. Our findings parallel the role of similar exhausted subsets of T cells during response to checkpoint blockade in murine models of chronic viral infection and human melanoma, now implicating them in adoptive cellular therapy and the GvL effect as well as defining their underlying regulatory circuitries. We also present a general computational framework that can be applied to high-dimensional temporal analyses of other cancer types and therapeutic scenarios beyond oncology.

## A global map of T cell states in the leukemic microenvironment

To delineate the evolving landscape of cellular phenotypic states for marrow-infiltrating T cells in relation to DLI therapy, we assembled a cohort of 12 patients treated with CD8-depleted DLI for relapsed CML^10^. Six patients were long-term DLI responders (“Rs”), defined as having achieved molecular remission (i.e. RT-PCR negative for the *BCR-ABL* transcript) after DLI, and 6 were nonresponders (“NRs”), who did not achieve measurable tumor reduction following DLI. None of the patients developed acute graft-versus-host disease (GvHD) after DLI, and the development of GvHD was unnecessary for DLI response (**Extended Data Table 1**). Serial bone marrow (BM) biopsies were collected before and after DLI treatment at a median of 3 timepoints per patient (**Suppl. Note 1**). The cohorts had comparable timing between allo-SCT and DLI therapy (median 702 (R) and 1064 (NR) days), and between pre- and post-DLI sampling (**Extended Data Fig. 1a**; **Suppl. Note 1; Extended Data Table 1**). As reference, we also analyzed post-transplant BM biopsies from 2 CML patients who never relapsed after allo-SCT. From each of the 41 total BM samples, we obtained scRNA-seq on viable mononuclear cells and chromatin accessibility profiles (using ATAC-seq) on isolated CD45RA^+^ and CD45RA^-^, CD4^+^ and CD8^+^ T cells (**Fig. 1a**, **Suppl. Note 1**).

**Fig. 1.**
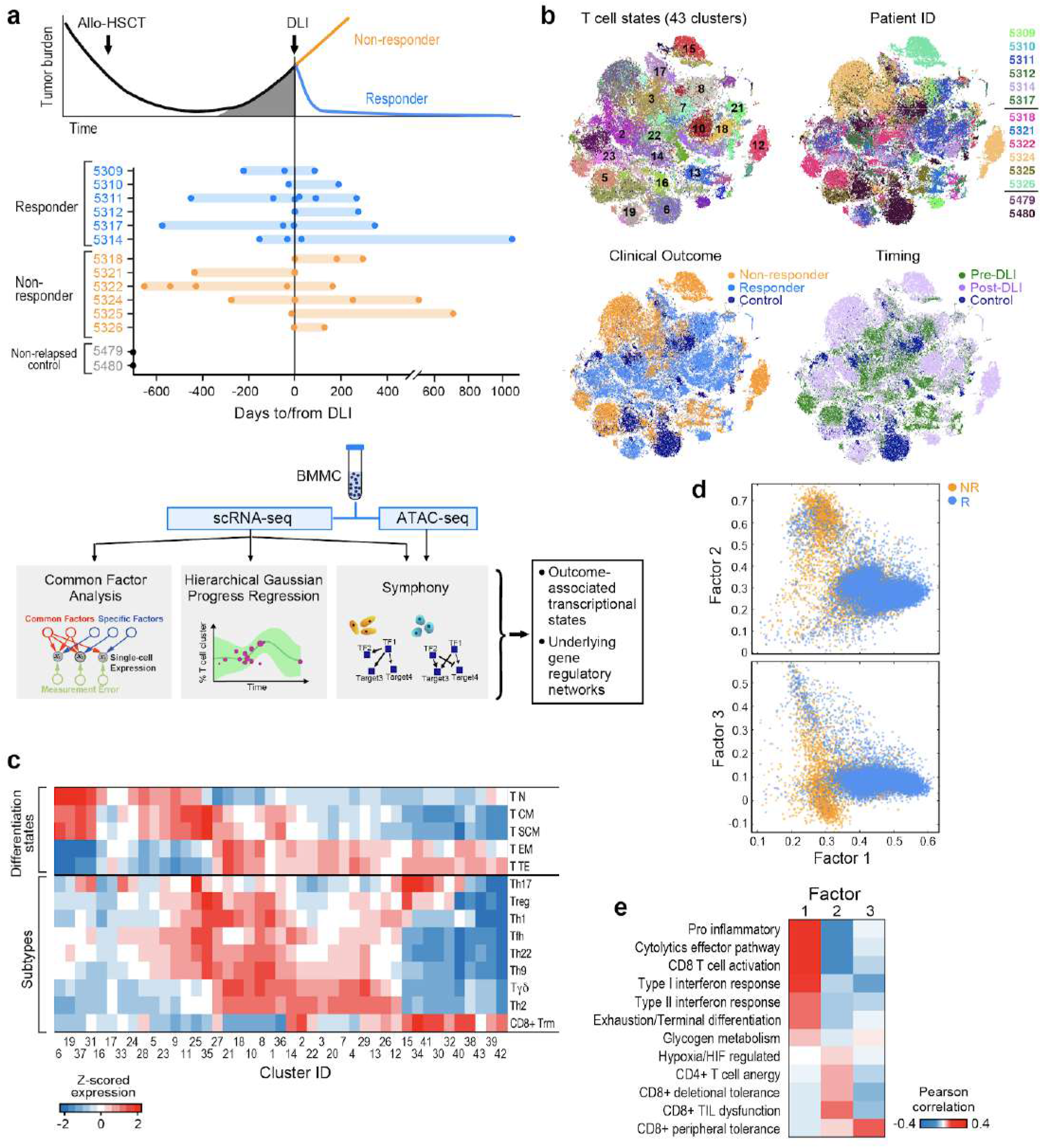
Experimental design and global map of T cell states. **a**, Clinical cohort, and flow chart of experimental and analysis schema. **b**, t-SNE projection of normalized scRNA-seq data for all T cells from 41 samples. Each dot represents a cell colored by cluster, patient ID, clinical outcome and timing respectively (expanded in Fig. S3). **c**, Mean expression for a curated set of transcriptomic signatures representing T cell subtypes and differentiation states for each T cell cluster; expression values are z-scored relative to all T cell clusters. **d**, Common Factor Analysis of T cells identifying 3 common latent factors distinguishing T cells between responders (R) and non-responders (NR). Each dot represents a cell colored by patient outcome and axes show factor loadings. **e**, Pearson correlation between common latent factors and mean expression of curated signatures.

In total, we identified 381,462 cells that passed our quality metrics, with a median of 8735 cells/sample (**Extended Data Table 2**). We used Phenograph^17^ to cluster the data into 62 distinct cell states, including subtypes of T, B, NK, monocytes, progenitor cells and CD34+ stem cells (**Suppl. Note 2**). Given the established critical role of T cells in the anti-leukemic potency of DLI^5^, we normalized and clustered the 87,939 T cells in our data, using Biscuit^18,19^ which robustly accounts for artifacts such as batch effects and library size variation (**Suppl. Note 2**). This analysis yielded 43 distinct T cell subsets spanning combinations of subtypes and differentiation states with variably expressed gene programs related to environmental stimuli (**Fig. 1b,c; Extended Data Fig. 1b-d**). For example, clusters 6, 19, 37 and 31 exhibited similar differentiation states and subtypes, for which we observed differential enrichment of pathways involving adenosine suppression, glucose deprivation, and anergy. Thus our global T cell map reveals substantial diversity corresponding to established T cell subtypes and states, marked by known and novel markers, that are shared across groups of patients.

## DLI resistance comprises multiple states of T cell dysfunction

While most T cell clusters were shared across patients, they were variably distributed across clinical features such as timing relative to DLI and clinical outcome (R vs NR) (**Extended Data Fig. 1e, Fig. 1b**), motivating us to identify the gene expression programs that might underlie these clinical variables. We tested standard techniques used to decompose single-cell data to identify trends underlying its variance (**Suppl. Note 2, Extended Data Fig. 2a**), but no principal or diffusion component was associated with R or NR status. Instead, we chose to use common factor analysis^20^, an unsupervised approach to uncover latent factors that explain shared variance across T cells, ignoring the portion of variance unique to cells (**Extended Data Fig. 2b**, **Suppl. Note 2**). Our rationale was that covariation across T cells can potentially capture factors underlying clinical response while de-emphasizing patient-specific variation. We identified 3 factors that explained 67% of the variation in our data which segregated R and NR T cells; co-variation in R T cells was found to be defined by Factor 1, while that in NR T cells was defined by Factors 2 and 3 (**Fig. 1d**). We associated each of these factors with manually curated gene sets relating to T cell biology and found Factor 1 to correlate with profiles associated with T cell activation (i.e. cytolytic effectors, interferon response, glycogen metabolism, CD8+ T cell activation, T cell exhaustion; **Fig. 1e**). We further confirmed enrichment of T cell exhaustion pre-DLI in R compared to NR, as previously observed^14^ (*P*<10^−6;^ **Extended Data Fig. 2c**). In contrast, Factors 2 and 3 correlated with non-overlapping signatures related to multiple, distinct T cell dysfunctional states (i.e. hypoxia, anergy, peripheral and deletional tolerance, tumor-infiltrating lymphocyte dysfunction; **Fig. 1e**, **Extended Data Fig. 2d**), suggesting that DLI resistance may be driven by not one, but multiple types of T cell dysfunction.

## DLI response is heralded by enrichment of activated and cytotoxic T cells prior to DLI

Given the substantial diversity of T cell subsets and gene programs in the leukemic microenvironment, we aimed to quantify this heterogeneity and study its change with outcome. T cell states are known to reside on continuous trajectories, which explain the majority of their variation^19,21,22^. We thus quantified their diversity across all clusters using phenotypic volume^19^, defined as the pseudo-determinant of covariance between genes. Phenotypic volume serves as a measure of the diversity of co-expressed transcriptional programs, which increases with the number and degree of independence of gene programs (**Suppl. Note 2**). We found substantially higher phenotypic diversity in pre-DLI Rs compared to pre-DLI NRs (**Fig. 2a**, log fold change=104.6, *P*<10^−6^), suggesting that diverse T cell phenotypes pre-DLI could be essential for response.

**Fig. 2.**
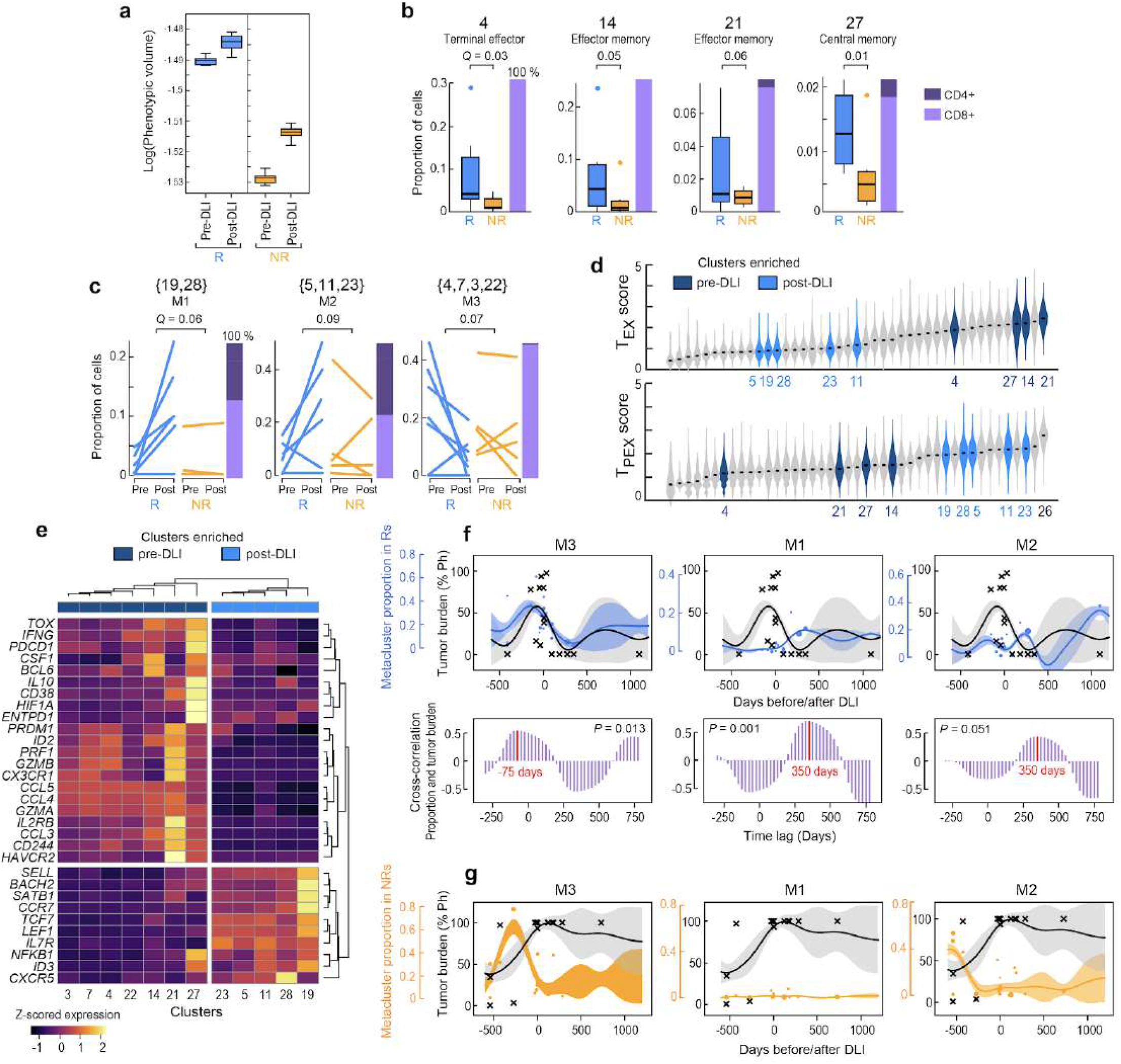
T cell states defining DLI response correspond to exhausted subsets. **a**, Phenotypic volume in log-scale (metric of transcriptional diversity^19^) of T cells before and after DLI in responders (R) and non-responders (NR). **b-c**, Proportion of T cells for pre-DLI only **(b)** or paired pre-/post-DLI **(c)** samples assigned to the indicated cluster or meta-cluster. *Q*-values determined from weighted t-test and empirical FDR estimation. Box plot elements display center line as median; box limits as first and third quartiles; whiskers extend to maximum/minimum data points (**a**) or 1.5x interquartile range with points as outliers (**b**). Each line in **(c)** indicates one patient. Stacked bars on the right indicate the proportion of CD4+ and CD8+ T cells. **d**, Violin plots showing density of T_EX_ (top) or T_PEX_ (bottom) viral signature scores^24^ across T cells grouped by cluster. Clusters are ordered by median score. Colored violins refer to clusters enriched in pre-DLI Rs (dark blue) or expanding in post-DLI Rs (light blue). Full labels provided in Fig. S5. **e**, Unsupervised hierarchical clustering based on tumor infiltrating T_PEX_ or T_EX_ genes^24^ segregates dark/light blue clusters. **f-g**, Hierarchical GP regression models (Suppl. Note 2; Extended Data Fig. 3f,g) for both the proportion of the indicated meta-cluster (in blue dots for Rs and in orange dots for NRs) and the percentage of tumor burden (in grey crosses) (indicated by percent positivity of the Philadelphia chromosome) per sample for Rs **(f)** or NRs (**g**). Each dot is one sample and dot size is proportional to sample size (total cells); inferred model mean is shown with lines and shaded area shows +/-1 standard deviation. Cross-correlation plots **(f**; purple) indicate the time shift between the models for meta-cluster proportion and tumor percentage, showing in-sync dynamics for M3 and tumor (left) and a lag between M1/M2 and tumor (middle, right).

In addition to finding increased overall phenotypic diversity in pre-DLI Rs, we next sought to identify distinct transcriptional states associated with clinical outcome. We tested each cluster for enrichment in baseline, pre-DLI samples from Rs compared to NRs (**Suppl. Table 1**). No cluster was consistently enriched in NRs, attesting to the notion of multiple pathways to DLI resistance rather than a common resistance mechanism shared across NRs. In contrast, within Rs, we identified four individual clusters (4, 14, 21, 27) that consistently enriched pre-DLI across responder patients (**Fig. 2b**, *FDR*<0.1); comprised predominantly CD8+ T cells; and shared the expression of genes involved in T cell activation (*CD160, HAVCR2, CD38*) and cytotoxicity (*CRTAM, GNLY, GZMK, GZMB*) (**Extended Data Fig. 3a**). Nevertheless, their distinct differentiation states (4, 14, 21: T_EM_/T_TE_; 27: T_CM_), subtypes (21: T_γδ_), and varied expression of chemokine receptors (14: *XCL2, CXCR4*; 21: *CXCR1*, *CXCR2*), tissue residency (14: *ITGA1*, *RGS1*; high score for “CD8+ T_RM_”) and cell cycle (27: *CDKN2A, TAF5, RRM2*) programs indicated the baseline diversity of these T cell states (**Fig. 1c**, **Extended Data Fig. 3a**).

We observed a marked increase in the number of T cell clusters in post-DLI samples compared to matched pre-DLI samples (mean 41 [range: 35-46] versus mean 38 [range: 34-41], *P*<0.001; **Suppl. Note 2**), suggesting that DLI expands the number of T cell transcriptional states. Indeed, both R and NR cases exhibited increases in phenotypic volume following DLI (*P*<10^−6^), (**Fig. 2a**). Rs displayed higher phenotypic volume than NRs at both pre- and post-DLI timepoints (*P*<10^−5^), whereas NRs displayed a far greater increase in phenotypic volume after DLI than Rs (*P*<10^−6^). Thus, despite an absent clinical response, NRs undergo marked T cell phenotypic remodeling. Of note, the phenotypic volumes of the non-relapsed reference samples were lower than samples from the study cohort, (*P*<10^−6^; **Extended Data Fig. 3b**). These results implicate more transcriptionally diverse local microenvironments within the leukemic bed that may persist even after leukemia remission following DLI.

## DLI response is marked by expansion of states consistent with precursor exhausted T cells

To identify T cell clusters that expand after DLI, we compared the cluster proportions in baseline pre-DLI samples to those from the remission timepoint following DLI. To increase our statistical power for detecting changes induced by DLI, we grouped transcriptionally similar clusters into meta-clusters (**Extended Data Fig. 3c**, **Suppl. Note 2**). In this fashion, we identified two meta-clusters which consistently expanded (M1:{19,28}, M2:{5,11,23}) and one that consistently contracted (M3:{4,7,3,22}) after DLI therapy, only in Rs (**Fig. 2c**). The T cell states that expanded in response to DLI comprised both CD4^+^ and CD8^+^ T cells; enriched for T_N_ (19, 28, and 5), T_CM_ (11), or both (23) states; and expressed corresponding gene programs for proliferation (*CDK20, CDK14, CDKL3*), lymph node homing (*SELL, CCR7*), and survival/self-renewal (*TCF7, IL7R, SATB1*) (**Extended Data Fig. 3a**). Analogous to the clusters enriched in pre-DLI R samples, the T cell states contracting in response to DLI comprised mostly CD8+ T cells, enriched similarly for T_EM_ and T_TE_ states, and expressed similar gene programs of cytotoxicity and activation. In contrast, no clusters or meta-clusters consistently changed in NRs.

Recent studies in murine models of chronic viral infection and cancer have delineated two major subsets of exhausted T cells that can be distinguished on the basis of gene expression signatures: terminal exhausted (T_EX_) cells, which possess superior cytotoxicity but shorter lifespan, and precursor exhausted (T_PEX_) cells which have greater polyfunctionality, expand following PD-1 blockade, and exert tumor control^23,24^. We hypothesized that the human CD8^+^ effector-like T cell clusters enriched pre-DLI and the rapidly expanding naive/memory-like T cell clusters enriched post-DLI might be phenotypically similar to these two subsets. Indeed, by scoring all clusters for T_EX_- or T_PEX_-defining signatures^24^, we found that clusters enriched in pre-DLI Rs (4, 14, 21, 27) scored highest for T_EX_ profiles whereas clusters consistently expanded post-DLI in Rs (M1, M2) scored highest for T_PEX_ profiles (**Fig. 2d**). Cluster 26 was the highest T_PEX_ scoring cluster and expanded only in R patient 5309 but did not meet the threshold for significance due to its small size and patient-dominant variation. Because patient 5309 was the only R without expansion in either of the two meta-clusters, M1 or M2 (**Extended Data Fig. 3d**), the expansion of cluster 26 suggests that all six Rs, in fact, demonstrated post-DLI expansion of T_PEX_ clusters. These T_EX_- or T_PEX_-defining signatures also segregated pre- and post-DLI enriched clusters in an unsupervised analysis (**Fig. 2e**). While pre-DLI enriched clusters expressed transcription factors (*TOX, ID2, PRDM1*), co-inhibitory receptors (*HAVCR2, PDCD1*, *ENTPD1, CD160, CD244*), chemokines and associated receptors (*CCL3, CCL4, CCL5, CX3CR1*), and effector molecules (*PRF1. GZMA. GZMB*) classically associated with T_EX_ cells, post-DLI enriched clusters expressed transcription factors (*TCF7, ID3, LEF1*), surface receptors (*CXCR5, IL7R*), and chromatin regulators (*SATB1*) consistent with T_PEX_ cells^23,25–31^ (**Fig. 2e**). Finally, unlike many studies using antigen-specific models of CD8^+^ T cell responses, we found a mixture of both CD4^+^ and CD8^+^ T cells to constitute these expanding T_PEX_-like clusters. Within the M1 and M2 meta-clusters, both subtypes exhibited global transcriptional similarity, with similar T_PEX_ scores and similar expression of key TFs such as *TCF7*, indicating the importance of both CD4^+^ and CD8^+^ subtypes to DLI response (**Extended Data Fig. 3e**).

Having identified response-associated T cell meta-clusters with diverging patterns after DLI (expanding M1 and M2, and contracting M3), we sought to characterize their evolution over time by merging samples across all time-points and clinical outcomes (**Suppl. Note 2**) and then modelling their temporal dynamics over the 4.5 year time period. To account for variability in timing, total cell number, and meta-cluster size on a per-sample basis, we constructed a hierarchical Gaussian Process (GP) regression model to capture dependencies between all pairs of time points per clinical group (R,NR) (**Extended Data Fig. 3f,g**; **Suppl. Note 2**). Our model revealed the M3 meta-cluster to gradually increase with leukemic growth in Rs and sharply contract during DLI response (time shift of 75 days; p=0.013, **Fig. 2f**, left) whereas both M1 and M2 meta-clusters robustly expanded as early as 3 weeks and endured as long as 3 years after DLI (**Fig. 2f**, middle, right). Notably, no association was detected between these meta-clusters and leukemic burden in NRs (**Fig. 2g**). Taken together, our data shows that reversal of T cell exhaustion is driven not by changes in gene expression, but rather by shifts in cell type composition – specifically, the expansion of T_PEX_ populations and contraction of T_EX_ subsets.

## Cell-state specific gene regulatory networks affirm exhausted subset identities

While recent work has described epigenetic T cell states that drive dedifferentiation^32^, effector “poising,”^33^ and exhaustion^34,35^, their relevance to clinical immunotherapeutic outcomes, especially following DLI, is unclear. To investigate the regulatory circuitry underlying the T cell transcriptional states associated with DLI outcome, we compared chromatin accessibility profiles between Rs and NRs (**Suppl. Note 3**). Consistent with our scRNA-seq analysis, we found increased chromatin accessibility in Rs in regions near T_PEX_- and T_EX_-associated genes (**Fig. 3a, Extended Data Fig 4a**), further supporting the association of these exhausted subsets with DLI response. Notably, we found similar accessibility for these genes among R samples, regardless of timing relative to DLI. In fact, we observed that the genome-wide accessibility landscape of T cells is more similar between pre- and post-DLI timepoints of Rs, than between Rs and NRs (**Fig 3b**), suggesting that DLI response involves selection of pre-existing epigenetic states as opposed to induction of global rewiring. This observation is consistent with our analysis of transcriptional states demonstrating that shifts in cell type composition underlie T cell phenotypic evolution during DLI response. Moreover, these results suggest the inflexibility of these epigenetic states of exhaustion in response to DLI, consistent with findings in murine models of chronic infection in response to PD-1 blockade^34,35^.

**Fig. 3.**
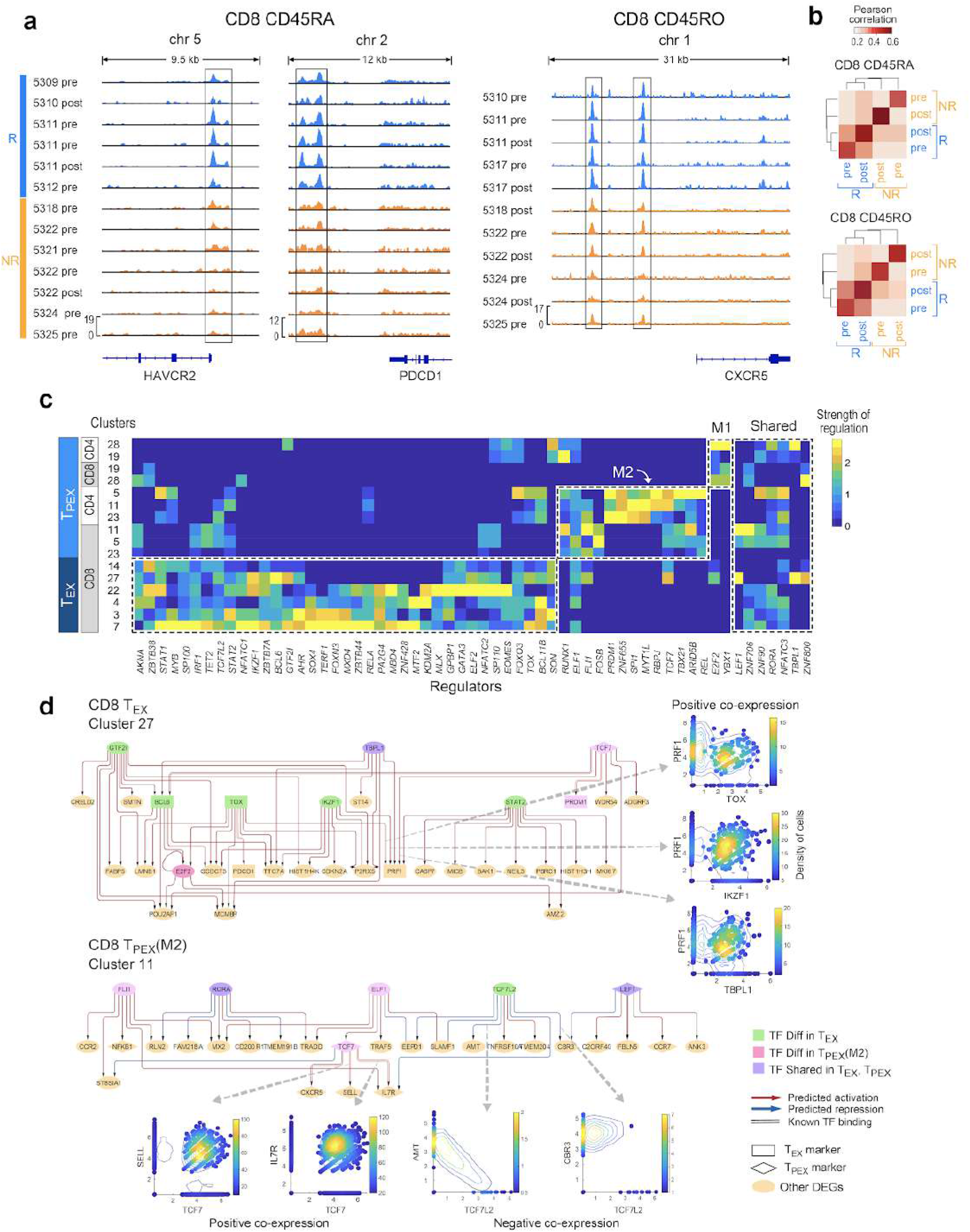
Epigenetic landscape and regulatory circuitry underlying T cell subsets. **a**, Chromatin accessibility signal from ATAC-seq data for CD8+ CD45RA+ (left) and CD8+ CD45RO+ (right) T cells indicating differential accessibility (*p*<0.05 indicated with boxes) between R and NR in regions near exhaustion marker genes. **b**, Average pairwise Pearson correlation between normalized ATAC-seq peak heights for CD8+ CD45RO+ (top) and CD8+ CD45RA+ (bottom) T cells from different clinical groups. **c**, Heatmap showing scaled values of predicted regulatory strength of TFs (i.e. magnitude of regulation independent of sign) from Symphony (Suppl. Note 3; Extended Data Fig. 4b), averaged across differentially expressed genes characterizing each cluster. Master regulators that are differential (t-test *p*<0.05) or shared between T_EX_ and T_PEX_ subsets are shown in dotted lines. **d**, Predicted regulatory circuitry for two example clusters; arrows between nodes indicate regulatory impact of a TF on a target gene. Master regulators that are differentially enriched in T_EX_ and T_PEX_ subsets are shown in green or pink nodes, respectively. Circuitry for other exhausted clusters are shown in Extended Data Fig 5.

To further study the circuitry underlying the distinct expanding T_PEX_ and contracting T_EX_ subsets, we developed Symphony^36^, a novel probabilistic multi-view model to infer gene regulation in each exhausted cluster (**Extended Data Fig 4b**). Symphony uses co-expression patterns between transcription factors (TF) and targets as evidence suggesting a potential regulatory impact. However, since co-expression between genes could be a by-product of indirect regulation or co-regulation, Symphony integrates scRNA-seq data with chromatin accessibility data from ATAC-seq, together with TF motif information to resolve direct links between genes. We first evaluated the performance of Symphony on data from well-characterized PBMCs^36^ and then confirmed the robustness of predicted links in our cohort with leave one (patient) out analysis (**Suppl. Note 3**).

To determine the strongest regulators underlying the differences in gene expression across the clusters, we summarized predicted regulatory networks in each cluster and defined master-regulators as TFs with strong average regulatory impact (either activation or repression) on the differentially expressed genes (DEGs) characterizing each cluster. Strikingly, the inferred master regulators organized into distinct groups associated with T_EX_ or T_PEX_ subsets (**Fig. 3c**). From our unsupervised analysis, we found many TFs previously known to associate with exhaustion in general (e.g. *EOMES*, *TBX21*)^37,38^ or regulate T_EX_ (e.g. *MYB*, *NFATC1, TOX*)^39^ and T_PEX_ subsets (e.g. *TCF7, PRDM1*, *LEF1*)^37^ in particular. Two of the identified TFs, *MTF2* and *GATA3*, were recently defined as mediators of intratumoral CD8^+^ T cell dysfunction in murine models^22^. While master regulators identified by T_EX_-associated DEGs were largely shared among disparate T_EX_ clusters, the two T_PEX_ meta-clusters were well-discriminated by two distinct sets of master regulators. We also observed a smaller group of master regulators including *LEF1* and *RORA* that were shared across T_PEX_ and T_EX_ subsets (**Fig. 3c**), suggesting a core shared regulatory program.

Despite shared master regulators even within highly related transcriptional T_EX_ or T_PEX_ states (dotted line boxes in **Fig. 3c**), Symphony revealed a distinct regulatory network architecture for each cluster (**Fig. 3d, Extended Data Fig. 5**) suggesting differences in wiring and target genes influenced by these regulators. Importantly, these cluster-specific regulatory networks imply that master regulators (shown in green, **Fig 3d** e.g. *TOX*) for pre-DLI enriched clusters appear to be directly linked to known T_EX_ markers; similarly, master regulators (shown in pink) for post-DLI enriched meta-clusters directly regulate known T_PEX_ markers. For example, in pre-DLI enriched cluster 27, *PDCD1* is inferred to be activated by *TOX*, while the effector molecule *PRF1* is predicted to be combinatorially activated by *TOX, IKZF1*, *TBPL1* and *STAT2* which are all up-regulated in this subset. Similarly, in post-DLI enriched cluster 11, *TCF7* acts as a hub, predicted to be regulated by *ELF1* and activating known T_PEX_ markers *IL7R*, *SELL* and *CXCR5* as expected. This connection between regulators found from our unbiased approach and known exhaustion markers, support the central role of these TFs in defining the identities of exhausted T cell clusters. Furthermore, their regulatory function, inferred with Symphony, is supported by evidence in TF and target gene co-expression (**Fig. 3d**) and/or chromatin accessibility (**Suppl. Note 3**). Thus, in addition to identifying known, exhaustion-related regulators driving these DLI response-associated T cell clusters, Symphony provides a roadmap for future investigation on the role of previously unexplored regulators.

## Confirmation of clonal properties and source of expanding T_PEX_ cells

In murine models, T_PEX_ and T_EX_ subsets have been reported to share a lineage relationship in which the former self-renews and gives rise to the latter^23^. For two Rs (5311, 5314), we used paired single-cell TCR- and RNA-seq to compare TCR clonotype sequences of T_PEX_ and T_EX_ clones (defined as >1 cell sharing the same TCR). We observed that 27% of T_PEX_ clones overlapped with T_EX_ clones (*p*<10^−14^ for both patients), confirming their lineage relationship (**Fig. 4a**; **Suppl. Note 4, Suppl. Table 7**). The expanded clones with T_PEX_ phenotype were predominantly CD4+ T cells (81%) and clones with T_EX_ phenotype were predominantly CD8+ T cells (99%) as were T_EX_/T_PEX_ overlapping clones (93%). T_EX_ clonotypes resided in larger clones than T_PEX_ clonotypes (**Extended Data Fig. 6a**). Clonotype diversity was higher in cells with a T_PEX_ phenotype than in those with a T_EX_ phenotype (*P*<0.05) for both patients (**Fig. 4b**), consistent with previous reports in murine and human studies^24^.

**Fig. 4.**
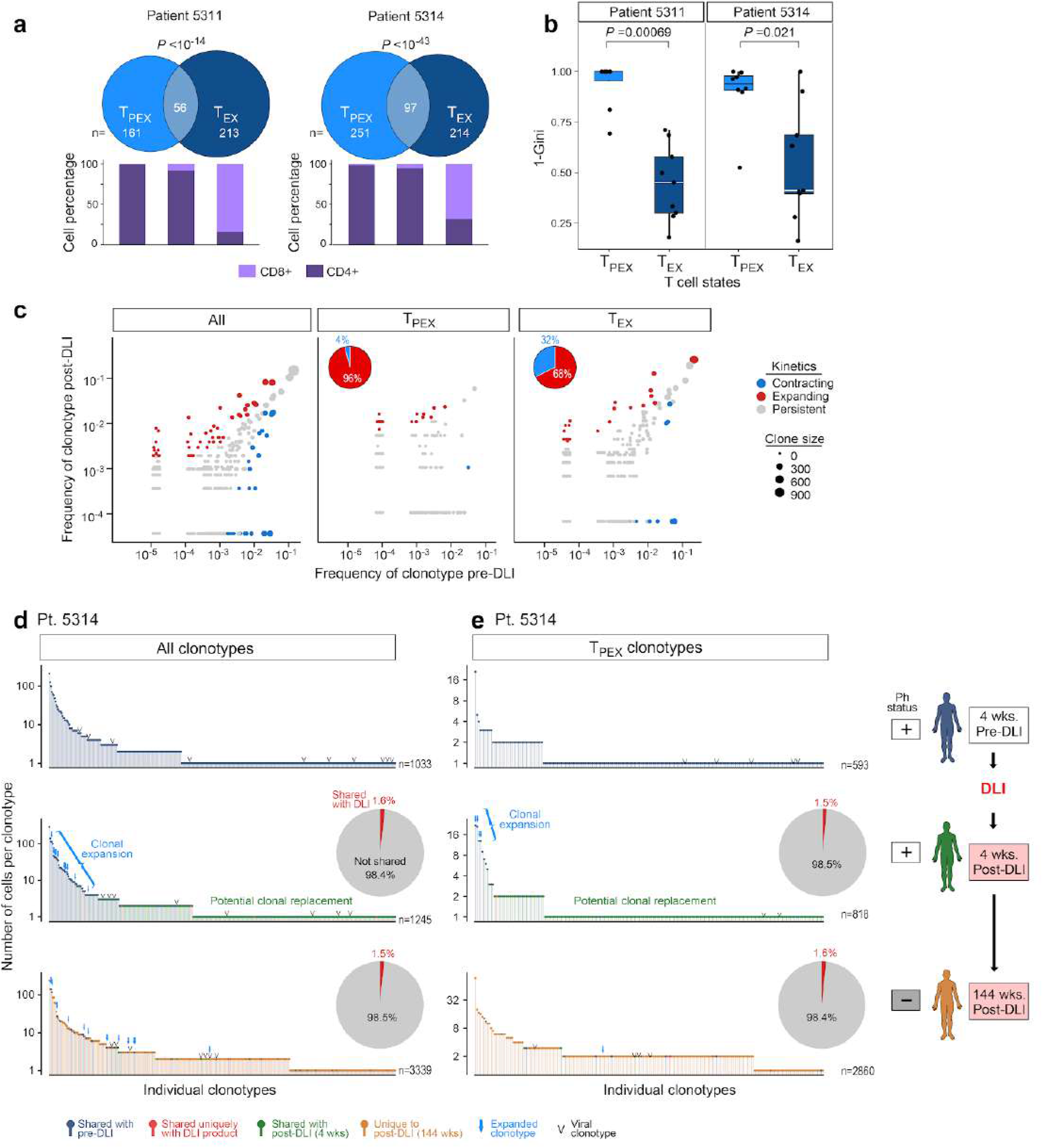
Confirmation of clonal properties and source of expanding T_PEX_ cells. **a**, Venn diagrams showing clonotype overlap between T_PEX_ and T_EX_ cells from two R patients (5311 and 5314), and stacked bars indicating percentage of CD8+ and CD4+ T cells in T_PEX_, T_EX_ and overlap categories. *P* value calculated from hypergeometric test. **b**, T_PEX_ clusters show increased TCR diversity (quantified with Gini coefficient; Suppl. Note 4) compared to T_EX_ clusters, Wilcoxon. Box plot elements display center line as median; box limits as first and third quartiles; whiskers extend to 1.5x interquartile range with points as outliers. **c**, Clonotype frequencies 1 month before and 1 month after DLI from both R patients. Each dot represents a clonotype with dot size proportional to size of clone for each cell subset. Expanding/contracting clonotypes determined with Fisher’s exact test (*P*<0.05). *Left*, clonotypes from all cells colored by one of three dynamic patterns: contracting, expanding, persistent. *Middle, right* dynamic clonotypes from T_PEX_ clusters are less likely to be contracting compared to clonotypes from T_EX_ clusters (pie charts). **d-e**, Frequency distribution of all (**d**) or T_PEX_ (**e**) clonotypes per time-point for patient 5314. Arrows indicate clonotype expansion from pre-DLI (*P*<0.05, Fisher’s exact test). Post-DLI clonotypes marked in red indicate unique match with DLI product and their proportions are displayed in pie charts for all (**d**) or T_PEX_ (**e**) post-DLI clonotypes (patient 5311 is shown in Extended Data Fig. 6b,c).

To study the dynamics of how clonal populations initially shifted in response to DLI in these two patients, we evaluated their TCR repertoire within one month before and after DLI and identified significantly expanding and contracting clonotypes (**Fig. 4c**, left). Consistent with our observation of expanding T_PEX_ states following DLI, dynamic clonotypes from T_PEX_ clusters were more likely to expand than contract compared to those from T_EX_ clusters (**Fig. 4c**, middle and right). Thus, the evolution of TCRs mirrors that of T_EX_/T_PEX_ transcriptional states after DLI.

We noted that clonally expanded TCRs following DLI were more likely to be shared with pre-DLI timepoints than were singletons, and many of these expanded clonotypes persisted even 3 years after DLI (**Fig. 4d,e**, left; **Extended Data Fig. 6b-d**; *P*<10^−15^, 4 wks and 144 wks post-DLI). Given that viral reactivity can be common in the post-transplant period^40^, we confirmed that viral antigen recognition only minimally accounted for the post-DLI clonotypes (<1.5% across the 2 patients) and did not explain the expansion or durability of T_PEX_ cells (**Extended Data Fig. 6e, Suppl. Note 4**). Upon examination of the source of expanding T_PEX_ states after DLI response, we found that only 1.4% of T_PEX_ cells from all post-DLI timepoints share clonotypes exclusively with the infusion product. These results demonstrate that DLI does not directly introduce the clonotypes that constitute T_PEX_ expansion (**Fig. 4d,e**, right; **Extended Data Fig. 6b,c**). Rather, post-DLI T_PEX_ cells consisted of expanding, pre-existing clonotypes as well as those that were not detected pre-DLI.

## DISCUSSION

In 1878, Leo Tolstoy published his masterpiece *<u>Anna Karenina</u>* and its eponymous principle that “all happy families are alike; each unhappy family is unhappy in its own way.” Likewise, our unbiased analysis of the evolution of T cell states following DLI unveiled common, shared pathways defining DLI response whereas multiple dysfunctional T cell states shaped DLI resistance, evoking a clinical outcome paradigm characteristic of other therapeutic scenarios where a limited set of targetable alterations predicts response in contrast to development of a diversified set of resistance mechanisms^41,42^.

To enable such clear insights from a limited patient cohort, we leveraged two critical features: samples collected from an informative clinical setting and innovative computational tools. Specifically, we exploited a scenario with unambiguous, binary clinical outcomes (response or resistance) in the absence of any toxicities; longitudinal sample collection; and uniform patient treatment with CD8-depleted DLI for relapsed CML in the absence of any confounding chemotherapy or immunomodulators. Furthermore, we consistently sampled the same bone marrow leukemic microenvironment for all patient-timepoints in contrast to studies in solid tumors where the sites of cancer involvement that are studied differ greatly even within the same patient^43^.

To overcome limitations of experimental design inherent to clinical studies such as variable timing of sample collection, patient heterogeneity, measurement uncertainty, and challenges in hypothesis testing on key populations, we adapted statistical techniques and developed novel longitudinal and integrative probabilistic models. Importantly, these computational approaches for dissecting global heterogeneity, identifying immune states related to dynamics of tumor burden, and integrative gene regulatory network inference are readily generalizable to other longitudinal, clinical settings. Indeed, with the increasing number of clinical correlative studies using longitudinal tumor biopsies^44,45^, we anticipate a growing need for such analytic frameworks.

Our findings, identified through direct interrogation of the human bone marrow microenvironment, dovetail with discoveries detected in model systems of chronic viral infections and solid tumors^21,46,47^. The pre- and post-DLI enriched T cell states we identified in Rs demonstrated dynamic, transcriptional, epigenetic and clonal hallmarks of T_EX_ and T_PEX_ exhaustion subsets, previously identified from murine models. Remarkably, the rapid expansion of T_PEX_-like states after DLI mirrored similar observations in these models during response to blockade of the PD-1 pathway in chronic viral infection^24,31,37,48,49^. In patients, recent studies have indicated a role for T_PEX_ cells during clinical outcomes to checkpoint blockade in advanced melanoma^24,43^. Our results now implicate the hierarchy of both T_EX_ and T_PEX_ subsets for human immunotherapeutic responses, extending the scope of their relevance beyond checkpoint blockade to adoptive cellular therapies for human leukemia and nominating this cellular program as a potent effector of GvL. Furthermore, these data confirm that reversal of T cell exhaustion is driven not by changes in gene expression, but rather by shifts in cell type composition - namely, expansion of T_PEX_ populations and contraction of T_EX_ subsets (**Fig. 5**). Because such distinctions cannot be delineated by bulk measurements, our findings highlight the advantages of single cell transcriptomics for discriminating between these possibilities.

**Fig. 5.**
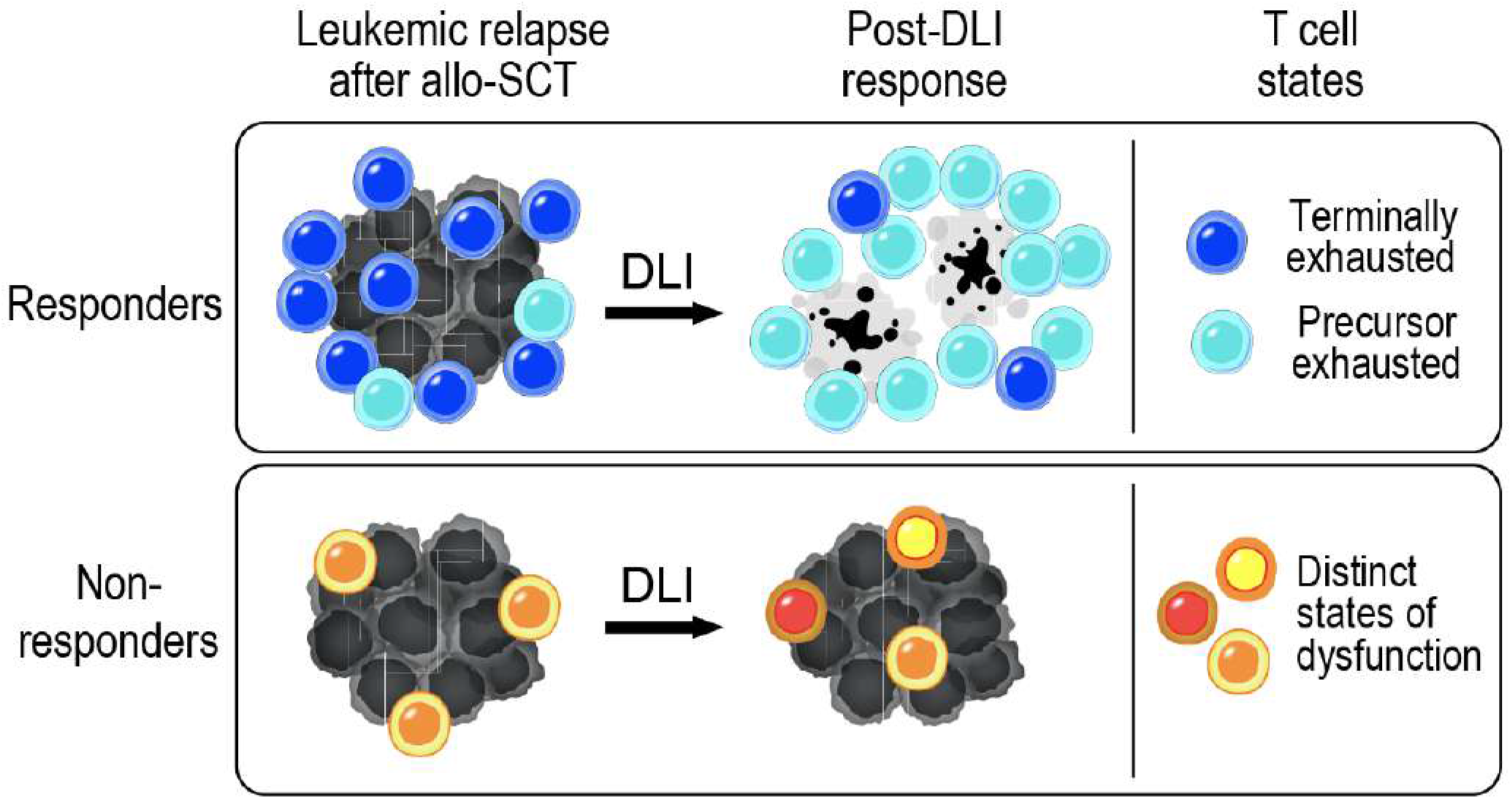
Summary model. Evolution of exhausted T cell states during DLI response and heterogeneity of distinct, dysfunctional T cell states during DLI resistance.

Our data moreover suggests novel mechanistic insight into DLI efficacy. Our scTCR analysis not only confirmed the lineage relationship between T_EX_ and T_PEX_-like states but now also explain that previous independent observations of increased TCR diversity detected in the setting of DLI response^12^ are a consequence of T_PEX_ subset expansion. Provocatively, this expansion of T_PEX_ cells during DLI response did not arise directly from the DLI product. Instead, we observed both recruitment of previously undetected clonotypes (potential clonal replacement^50^) and expansion of pre-existing ones (clonal expansion), suggesting that immunologic ‘help’ from DLI, rather than direct transfer of anti-leukemic T cells, drove leukemic remission. Similar results have been observed in murine models of exhaustion reversal after adoptive transfer of CD4+ T cells^51,52^. These data suggest that T_EX_/T_PEX_ subsets serve as both marker and mechanism for DLI response. Our findings motivate future clinical trial designs to test the status of T_EX_ cells as a biomarker for predicting DLI response and to evaluate therapeutic strategies that enhance T_PEX_ recruitment and expansion. Pursuing such approaches offers the possibility of enhancing the GvL effect during relapse after allo-SCT.

Functional interrogation of the novel regulatory networks proposed by our joint analysis of scRNA- and bulk ATAC-seq datasets through Symphony should accelerate these efforts. Future studies should also address the mechanism of DLI-induced T_PEX_ expansion and evaluate its relevance for newer adoptive cellular therapies such as chimeric antigen receptor T cells. In addition, while these T cell exhausted subsets have now been observed in multiple clinical settings, which aspects of their underlying molecular machinery and distinct regulatory circuits remain specific to the leukemic or GvL setting and which generally extend to other cancers and human diseases should be explored. Finally, our analytic approaches serve as a template for future studies that seek to harness such multidimensional data sets for clinical and therapeutic relevance.

## Acknowledgments

We thank Satyen Gohil, John Ray, Jaeyoung Chun, Manu Setty, Daniel Kim, and Tal Nawy for their valuable discussions on the generation and analysis of ATAC and scRNA-seq data, and Sandhya Prabhakaran for discussions on Symphony. We also thank all members of the Wu and Pe’er laboratories for helpful discussions, appreciate Haesook Kim for confirming clinical data from BMT repository, and thank Brian Miller and Eva Petschnigg for valuable discussion of the manuscript. Finally, we are grateful for the study nurses and clinical staff that obtained samples, the Pasquarello Tissue Bank in Hematologic Malignancies for processing and banking samples, and the patients who generously consented for the research use of these samples.

This work was supported in part by the NCI grants 1R01CA155010 (C.J.W.), P01CA206978 (C.J.W.), U10CA180861 (C.J.W.), P01CA229092 (R.J.S, J.R, C.J.W), and U54CA209975 (D.P.), and Ludwig Cancer Research (D.P.). P.B. was supported by a Physician-Scientist Training Award from the Damon Runyon Cancer Research Foundation, an Amy Strelzer Manasevit Scholar Award from the Be The Match Foundation, and an American Society of Hematology Fellow Scholar Award. E.A. was supported by NCI grant K99CA230195 and an American Cancer Society Postdoctoral Fellowship (PF-17-243-01-RMC). D.N. is supported by a grant from the NIH (5P30 CA006516). C.J.W. is a Scholar of the Leukemia and Lymphoma Society.

## Author Contributions

P.B. and E.A. conceived the study. P.B., E.A., D.P., and C.J.W. supervised the study. P.B., V.N.N., S.L., and K.J.L. designed and performed experiments. E.A., C.B., and D.P. designed and developed Symphony. E.A., Z-N.C., and D.P. developed statistical techniques and GP model. P.B., E.A., C.E., D.N., D.P. and C.J.W. analyzed and interpreted results. R.J.S., J.R., and E.P.A. designed and conducted DLI clinical trials. R.J.S., J.R., E.P.A., and C.J.W. oversaw patient care, contributed samples, and provided clinical data from the BMT repository. P.B., E.A., D.P., and C.J.W. wrote the manuscript with input from all authors.

## Data and materials availability

Single cell transcriptome and TCR as well as chromatin accessibility data will be submitted to NCBI’s Database of Genotypes and Phenotype (dbGaP; https://www.ncbi.nlm.nih.gov/gap) under study number: [will be available before publication].

## Code availability

The hierarchical Gaussian Process model is implemented using the probabilistic programming language pyro^53^ available at: https://github.com/dpeerlab/dli_gpr. The integrative model Symphony is implemented using the probabilistic language Edward^54^ with code available at: https://github.com/dpeerlab/Symphony with input data accessible for reviewers.

---

**Extended Data Fig. 1.**
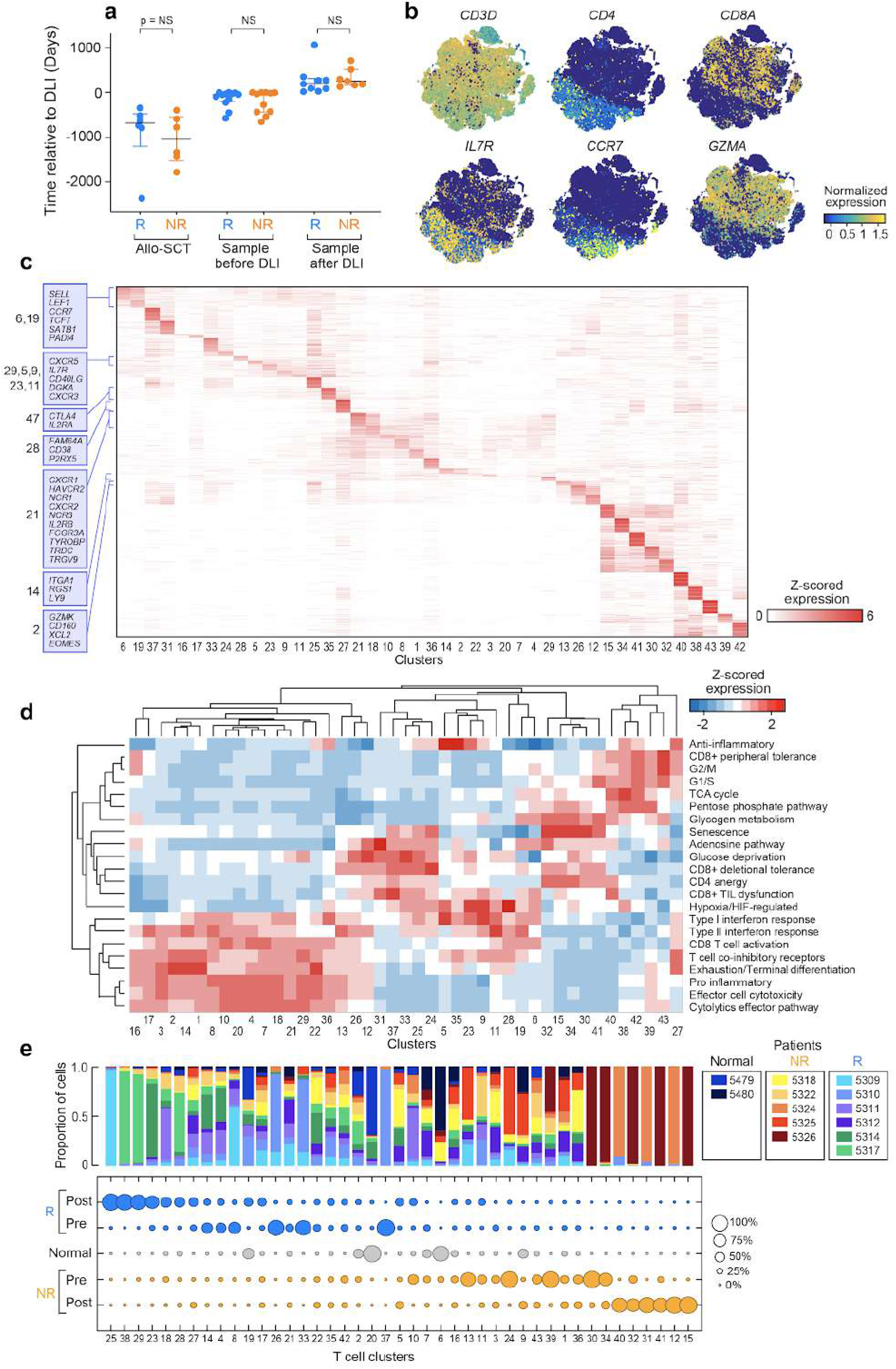
Clinical variables, biological features, and cluster distributions. **a**, Time from stem cell transplant (SCT) to DLI for each patient (left), from pre-DLI sample to DLI (middle) and from DLI to post-DLI sample (right). Center line as median with interquartile range and points as outliers. **b**, t-SNEs of all 43 T cell clusters, colored by indicated gene, recapitulating known immunobiology. **c**, Heatmap of differentially expressed genes (DEGs) per cluster for all clusters (full list provided in **Suppl. Table 3**). Indicated are informative genes for immune subtypes or differentiation states. **d**, Heatmap of mean expression for a curated set of signatures for each T cell cluster; expression values are z-scored relative to all T cell clusters. **e**, Distribution of clusters across patients (*top*, stacked bar plots) and clinical groups and timepoints (*bottom*, bubble plot).

**Extended Data Fig. 2.**
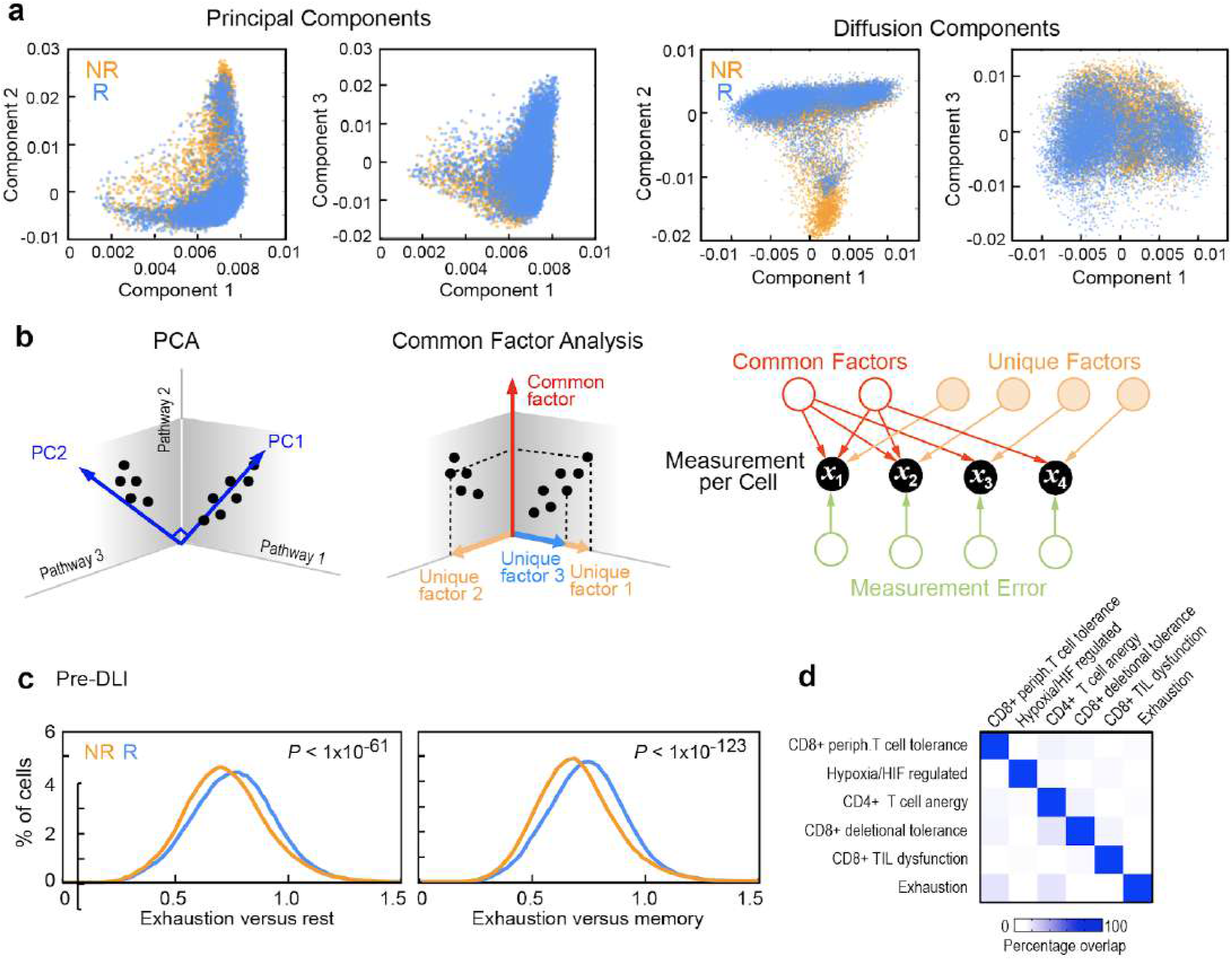
Identifying outcome-associated gene expression programs. **a**, Top three principal (**left**) or diffusion^55^ (**right**) components fail to discriminate clinical outcomes; each dot is a T cell colored by patient outcome **b**, Cartoon illustration for common factor analysis vs PCA. **c**, Distribution of scores for exhaustion signatures across cells, confirming increased exhaustion in pre-DLI T cells from Rs compared to NRs. **d**, Low percentage overlap of dysfunctional gene sets indicating discrete forms of T cell dysfunction associated with Factors 2,3 in Fig 1e.

**Extended Data Fig. 3.**
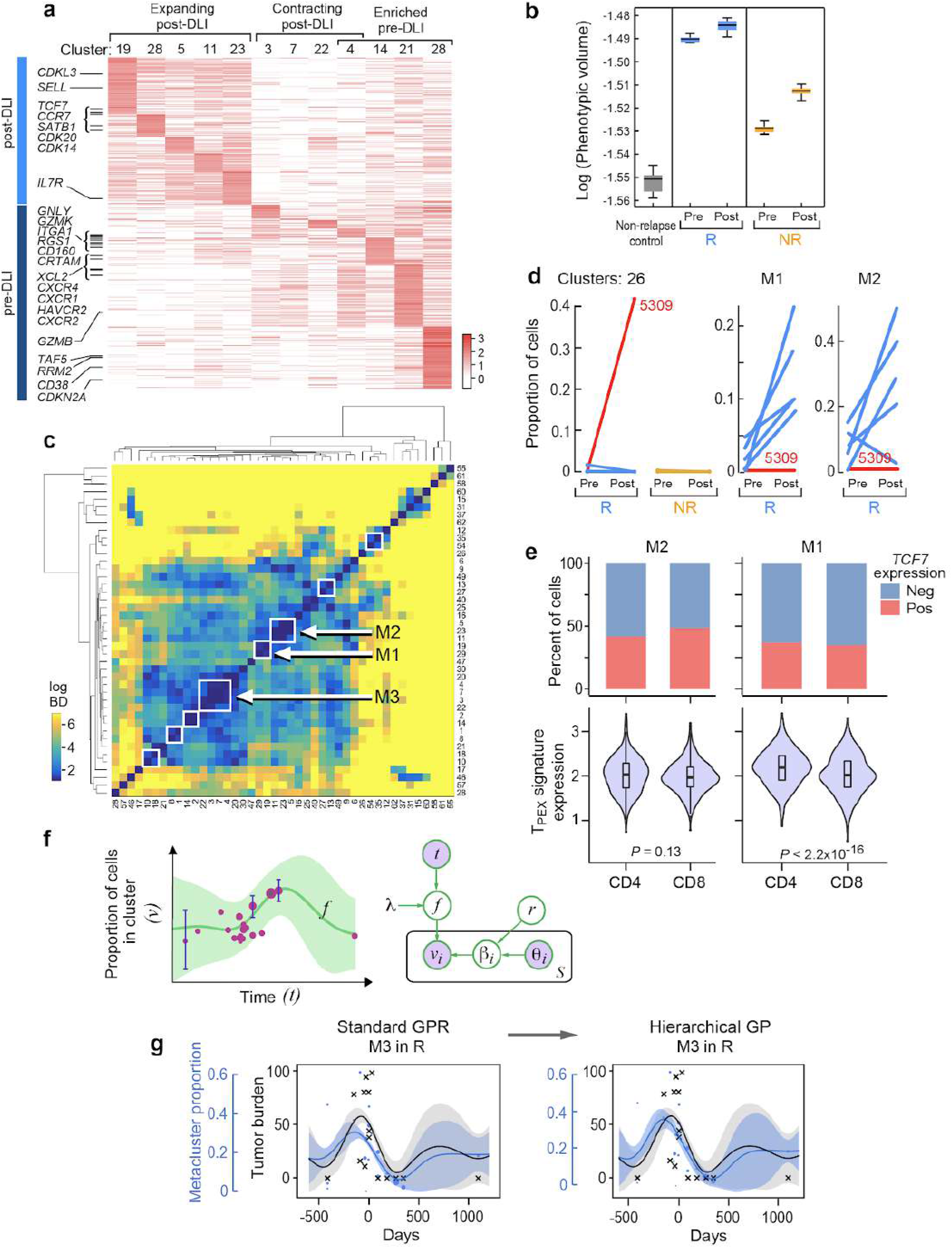
Expression profiles and longitudinal dynamics of pre- and post-DLI enriched clusters. **a**, Heatmap of normalized gene expression values for differentially expressed genes from Ext Data Fig 1c, subsetted for clusters enriched pre- or post-DLI. Center line in boxplots indicate the median and boxes indicate 25th and 75th percentiles. Whiskers extend to extreme data points. **b**, Phenotypic volume in log-scale (metric of transcriptional diversity) of T cells from non-relapse controls and before and after DLI in responders (R) and non-responders (NR). **c**, Heatmap of Bhattacharyya distances (BD) in log scale between pairs of clusters; closest clusters are grouped to form meta-clusters (white boxes). Meta-clusters significantly expanding (M1, M2) or contracting (M3) in Rs post-DLI are labeled. **d**, Proportion of T cells for paired pre-/post-DLI samples assigned to cluster 26 (left) or T_PEX_ meta-clusters (middle, right) with patient 5309 labeled in red. *Q* value determined from weighted t-test and empirical FDR estimation. **e**, Percentage of CD4+ or CD8+ T cells within M1 or M2 meta-clusters that expressed the *TCF7* gene (top). Violin plots of T_PEX_ gene set expression for CD4+ versus CD8+ T cells within M1 and M2 meta-clusters (bottom). **f**, Cartoon illustration (left) and plate model (right) for a hierarchical Gaussian Process (GP) model for inferring dynamics of clusters over time while accounting for measurement uncertainty. **g**, Inferred model for proportion of M3 meta-cluster (blue) using the hierarchical GP (right) compared to a standard GP (left); each blue dot represents a sample with dot size proportional to sample size (i.e. total number of T cells in the sample); blue lines show model mean and shaded area show +/-1 standard deviation (SD); grey crosses represent tumor burden data; grey line and shaded area show mean and SD for tumor burden model.

**Extended Data Fig. 4.**
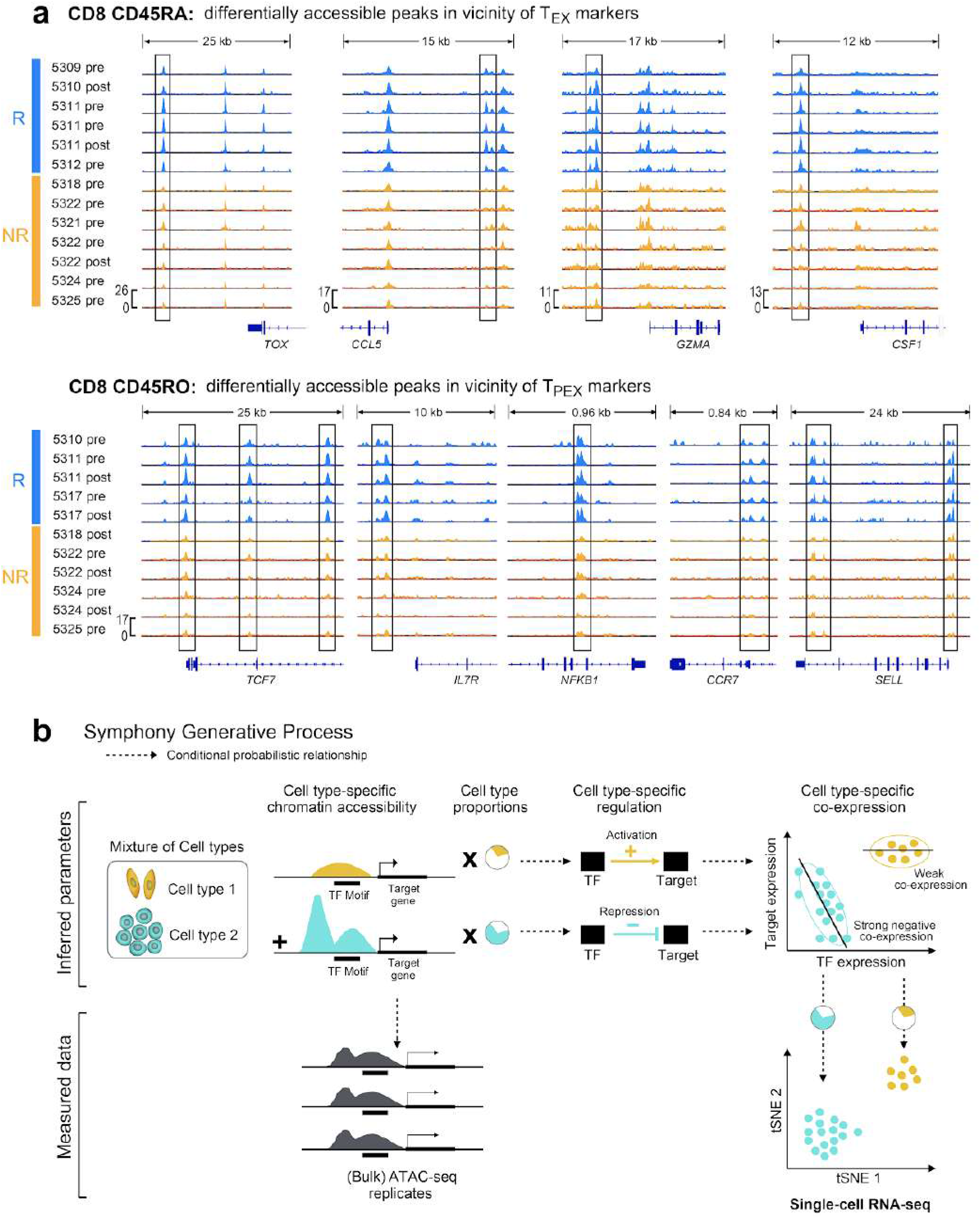
Epigenetic profiles of T cells and integrative modeling of regulation. **a**, Chromatin accessibility signal from ATAC-seq data for CD8+ CD45RA+ (top) and CD8+ CD45RO+ (bottom) T cells indicating differential accessibility (*p*<0.05 indicated with boxes) between R and NR in regions near exhaustion marker genes. **b**, Generative process for Symphony, a novel probabilistic model that infers regulation from integration of ATAC-seq and scRNA-seq data.

**Extended Data Fig. 5.**
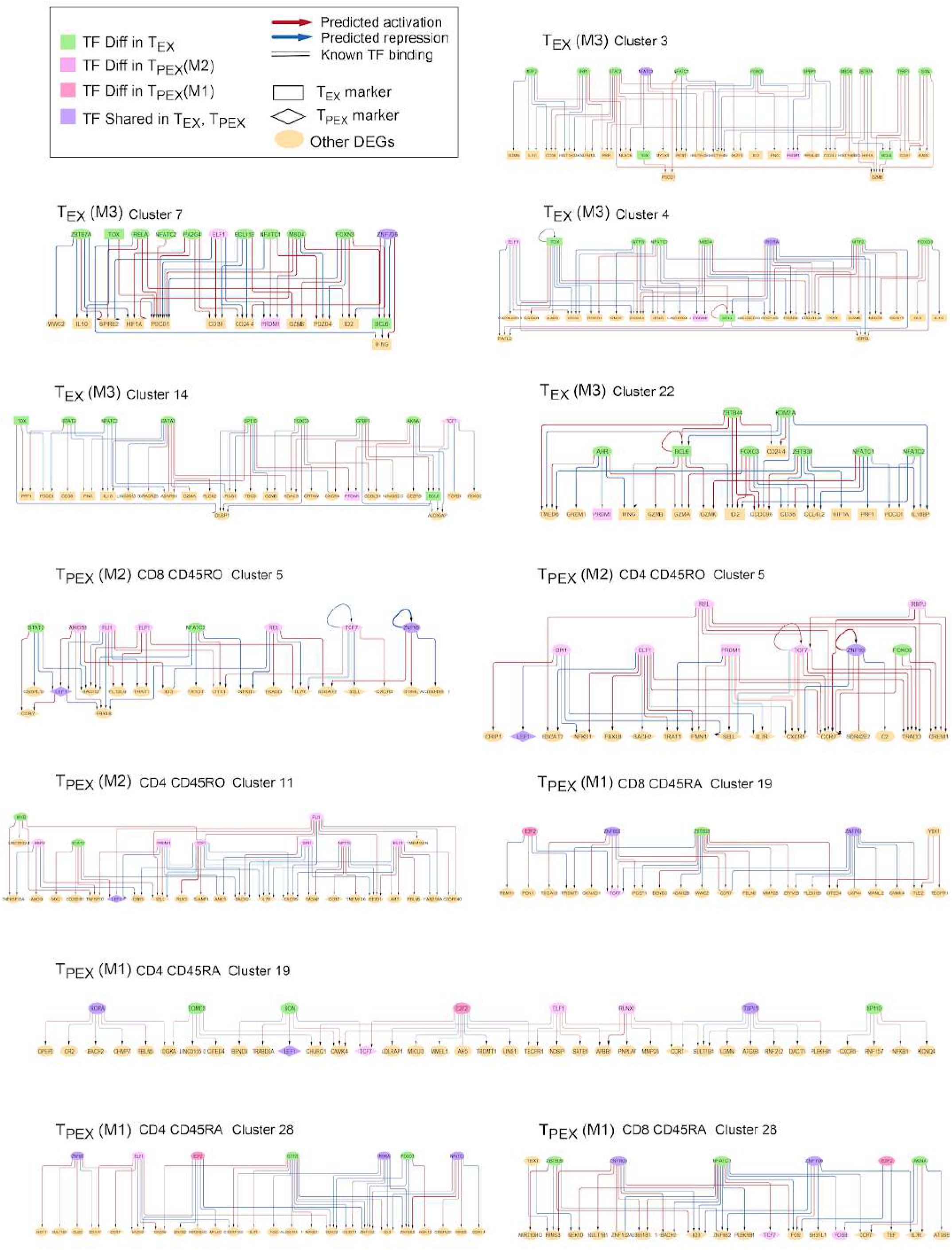
Variability in regulatory network architecture across clusters. Predicted regulatory circuitry for exhausted T cell clusters. Arrows between nodes indicate predicted regulatory impact of a TF on a target gene. Master regulators that are differentially enriched in T_EX_ or T_PEX_ subsets are shown in green or pink nodes respectively.

**Extended Data Fig. 6.**
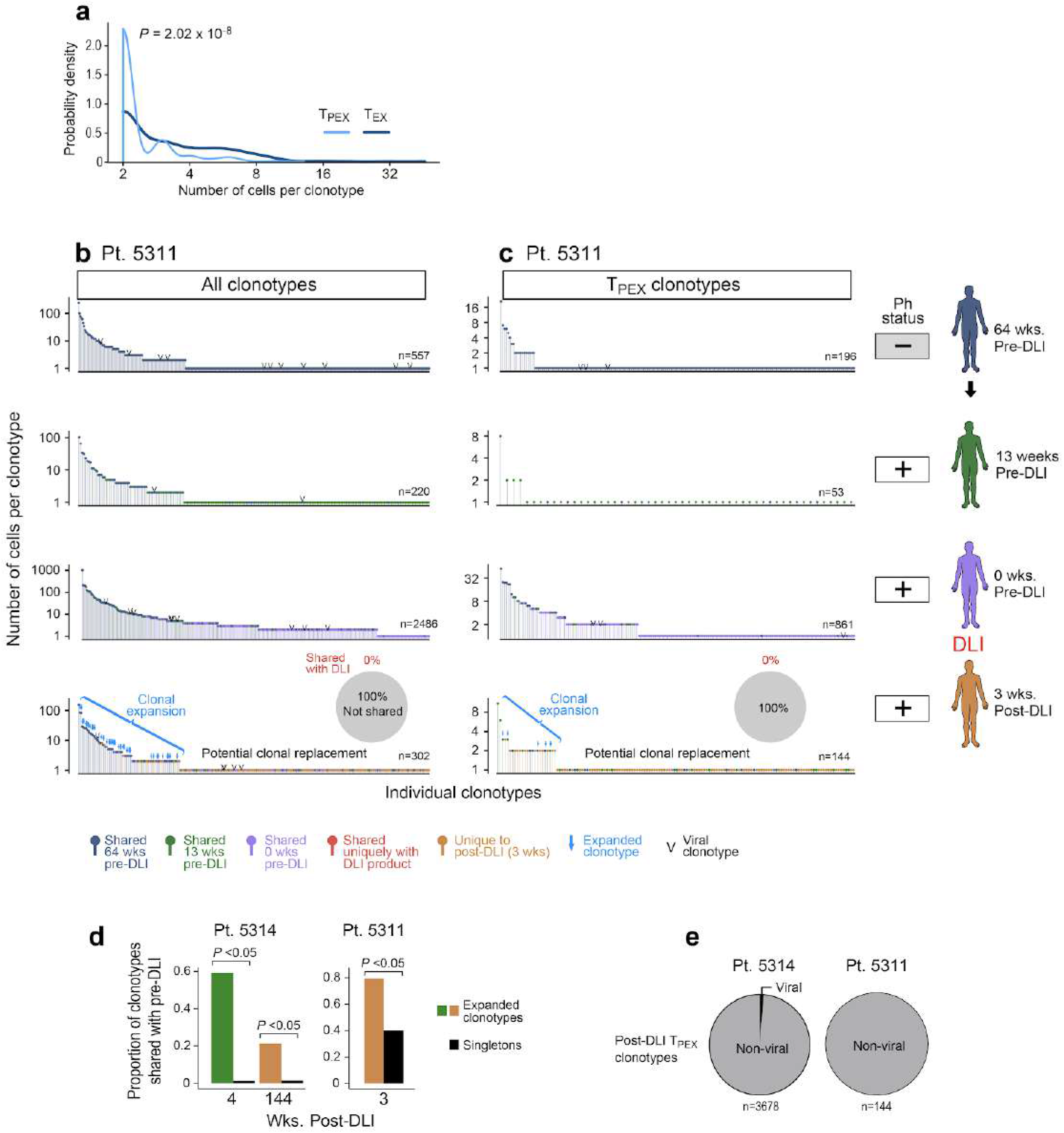
Clonotype properties, distributions and evolution during response to DLI. **a**, Probability densities of clone sizes for all T_EX_ and T_PEX_ cells from samples derived from R patients 5314 and 5311. **b-c**, Frequency distribution of all (**b**) or T_PEX_ (**c**) clonotypes per timepoint for patient 5311. Arrows indicate *P*<0.05, Exact Fisher’s Test, for clonotype expansion from pre-DLI. Post-DLI clonotypes marked in red indicate unique match with DLI product and their proportions are displayed in pie charts for all (**b**) or T_PEX_ (**c**) post-DLI clonotypes. **d**, Barplots of proportions of expanded versus singleton clonotypes shared with pre-DLI samples from each post-DLI timepoint from 5314 (n=2 post-DLI samples) and 5311 (n=1 post-DLI sample). **e**, Pie charts displaying the proportion of post-DLI T_PEX_ clonotypes matching publicly available viral-specific clonotypes.

**Extended Data Table 1.** Patient characteristics

| DFCI ID | Age at SCT | Donor source | Pt/Dnr Sex | Donor type | Type of transplant | GvHD ppx | Phase | Time from allo-SCT to DLI (days) | Best response | Max aGvHD grade | Max cGvHD grade |
| --- | --- | --- | --- | --- | --- | --- | --- | --- | --- | --- | --- |
| 5309 | 33 | BM | F/M | MRD | MA | TCD | Chronic | 362 | MR | 0 | Ext |
| 5310 | 34 | BM | M/F | MRD | MA | TCD | Chronic | 2371 | MR | 0 | N/A |
| 5311 | 49 | BM | M/M | MRD | MA | TCD | Chronic | 817 | MR | 0 | Lim |
| 5312 | 47 | BM | M/M | MRD | MA | TCD | Chronic | 649 | MR | 0 | Lim |
| 5314 | 57 | BM | F/F | MRD | MA | TCD | Chronic | 550 | MR | 0 | N/A |
| 5317 | 33 | BM | M/F | MRD | MA | TCD | Chronic | 755 | MR | 1 | Ext |
| 5318 | 31 | BM | F/M | MRD | MA | TCD | Chronic | 1330 | Progression | 0 | N/A |
| 5321 | 50 | BM | F/F | MRD | MA | TCD | Chronic | 797 | Progression | 1 | N/A |
| 5322 | 44 | BM | M/M | MRD | MA | TCD | Chronic | 1462 | Progression | 0 | N/A |
| 5324 | 39 | BM | M/M | MRD | MA | TCD | Chronic | 422 | Progression | 0 | N/A |
| 5325 | 49 | BM | F/M | MRD | MA | TCD | Chronic | 615 | Progression | 0 | N/A |
| 5326 | 48 | BM | F/M | MRD | MA | TCD | Blast crisis | 1787 | Progression | 1 | N/A |
| 5479 | 31 | BM | M/F | MRD | MA | TCD | Accelerated | No relapse | - | 0 | 0 |
| 5480 | 33 | BM | F/M | MRD | MA | TCD | Chronic | No relapse | - | 0 | N/A |

BM: bone marrow; F: female; M: male; MRD: matched related donor; MA: myeloablative; TCD: T cell depletion; MR: molecular response; Ext: extensive; Lim: limited; N/A: not available

**Extended Data Table 2.** Sample characteristics and QC metrics

| Patient ID | Outcome | Sample ID | Time relative to DLI (days) | Tumor burden (%Ph+) | # cells | # reads (mill) | median mols per cell |
| --- | --- | --- | --- | --- | --- | --- | --- |
| 5309 | Responder | B05 | -45 | 11 | 7228 | 294 | 2955 |
|  |  | B06 | 86 | 0 | 7462 | 295 | 2575 |
|  |  | B40 | -223 | 0 | 10346 | 686 | 3029 |
| 5310 | Responder | B01 | -27 | 94 | 4274 | 283 | 2840 |
|  |  | B02 | 190 | 0 | 5791 | 316 | 3199 |
| 5311 | Responder | B09 | 0 | 44 | 8780 | 321 | 2656 |
|  |  | B10 | 19 | ND | 4728 | 255 | 1540 |
|  |  | B11 | 91 | 0 | 8589 | 294 | 2692 |
|  |  | B12 | 268 | 0 | 14691 | 387 | 2783 |
|  |  | B29 | -451 | 0 | 6376 | 177 | 1273 |
|  |  | B30 | -94 | 16 | 6062 | 344 | 1666 |
| 5312 | Responder | B21 | 0 | 38 | 7266 | 212 | 2009 |
|  |  | B22 | 276 | 0 | 1279 | 175 | 1041 |
| 5314 | Responder | B25 | -33 | 94 | 9011 | 377 | 2090 |
|  |  | B26 | 1059 | 0 | 4465 | 313 | 2456 |
|  |  | B31 | -152 | 78 | 5994 | 374 | 2538 |
|  |  | B32 | 29 | 98 | 8058 | 379 | 2548 |
| 5317 | Responder | B23 | -5 | 80 | 12647 | 555 | 6694 |
|  |  | B24 | 347 | 0 | 14127 | 201 | 1893 |
|  |  | B38 | -574 | 0 | 10777 | 409 | 2756 |
|  |  | B39 | -50 | 80 | 8370 | 307 | 2389 |
| 5318 | Non-Responder | B27 | 0 | 100 | 9971 | 403 | 2445 |
|  |  | B28 | 182 | 100 | 4029 | 347 | 1452 |
|  |  | B41 | 295 | 100 | 4983 | 351 | 2313 |
| 5321 | Non-Responder | B42 | -435 | 0 | 10365 | 434 | 1867 |
|  |  | B43 | 0 | 100 | 8720 | 590 | 3117 |
| 5322 | Non-Responder | B03 | -34 | 100 | 9789 | 295 | 2015 |
|  |  | B04 | 164 | 100 | 10573 | 445 | 3439 |
|  |  | B33 | -652 | 0 | 9290 | 390 | 1723 |
|  |  | B34 | -540 | 34 | 8487 | 390 | 2373 |
|  |  | B35 | -428 | 97 | 11083 | 415 | 1630 |
| 5324 | Non-Responder | B07 | 0 | 93 | 7591 | 353 | 3042 |
|  |  | B08 | 252 | ND | 7247 | 306 | 3024 |
|  |  | B36 | -274 | 3 | 6234 | 318 | 1282 |
|  |  | B37 | 509 | 0 | 13874 | 305 | 2789 |
| 5325 | Non-Responder | B17 | -15 | 100 | 9027 | 321 | 2111 |
|  |  | B18 | 710 | 100 | 8735 | 288 | 2490 |
| 5326 | Non-Responder | B19 | -4 | 100 | 11720 | 270 | 3635 |
|  |  | B20 | 129 | 100 | 11880 | 350 | 7194 |
| 5479 | No relapse control | B44 | - | - | 9487 | 410 | 2602 |
| 5480 | No relapse control | B46 | - | - | 16247 | 413 | 2223 |

ND: not determined

**Extended Data Table 3.** Terminal and precursor exhaustion signatures^23^

| **Terminal exhaustion** | **Precursor exhaustion** |
| --- | --- |
| *CD3G* | *TCF7* |
| *FASLG* | *MS4A4A* |
| *ID2* | *TNFSF8* |
| *LAG3* | *CXCL10* |
| *RGS1* | *EEF1B2* |
| *CCL3 CCL3L1* | *ID3* |
| *KIAA1671* | *IL7R* |
| *SH2D2A* | *JUN* |
| *DUSP2* | *LTB* |
| *PDCD1* | *XCL1* |
| *CD7* | *SOCS3* |
| *NR4A2* | *TRAF1* |
| *CD160* | *EMB* |
| *PTPN22* | *CRTAM* |
| *ABI3* | *EEF1G* |
| *PTGER4* | *CD9* |
| *GZMK* | *ITGB1* |
| *GZMA* | *GPR183* |
| *MBNL1* | *ZFP36L1* |
| *VMP1* | *SLAMF6* |
| *PLAC8* | *LY6E* |
| *RGS3* |  |
| *EFHD2* |  |
| *GLRX* |  |
| *CXCR6* |  |
| *ARL6IP1* |  |
| *CCL4* |  |
| *ISG15* |  |
| *LAX1* |  |
| *CD8A* |  |
| *SERPINA3* |  |
| *GZMB* |  |

## Supplementary Information

is available for this paper.

Supplementary Note 1: <u>Sample details and library preparation</u>. Describes clinical cohort characteristics, bone marrow sample processing and isolation/preparation of cells for scRNA-/TCR-seq as well as bulk ATAC-seq.

Supplementary Note 2: <u>Single-cell RNA-seq data analysis</u>. Describes processing and analytic pipelines for scRNA-seq analysis; cluster visualization and annotation; common factor analysis to identify factors relating to clinical outcome; identification of meta-clusters and enriched clusters; and use of Gaussian Process Regression models to track cluster temporal dynamics in relation to tumor burden.

Supplementary Note 3: <u>Integration of single-cell RNA-seq and ATAC-seq</u>. Describes pre-processing of ATAC-seq data; correlations between accessibility profiles; and development and use of Symphony to infer cluster-specific gene regulatory networks.

Supplementary Note 4: <u>Analysis of paired single-cell TCR- and RNA-seq</u>. Describes preprocessing and identification of both exhausted clusters and TCR clonotypes.

## Supplementary Tables

Supplementary Table 1: Clusters, cell numbers, and distributions across patients and samples

Supplementary Table 2: Cluster mean expression

Supplementary Table 3: Differentially expressed genes (DEGs) per cluster

Supplementary Table 4: Signatures

Supplementary Table 5: ATAC-seq samples and QC metrics

Supplementary Table 6: Paired scTCR-seq and RNA-seq samples and QC metrics

## Supplementary Information

### Supplementary Note 1: Sample details and library preparation

#### Sample collection

Bone marrow (BM) biopsies were obtained pre- and post-DLI after relapse following allo-SCT (or during remission following allo-SCT) from patients enrolled in Dana-Farber Cancer Institute (DFCI) clinical trials (94-009, 95-011, 96-372, 96-022, and 96-277) between 1994-2001 that were approved by the DFCI Human Subjects Protection Committee. These studies were conducted in accordance with the Declaration of Helsinki. Bone marrow mononuclear cells (BMMCs) were isolated via Ficoll-Hypaque density gradient centrifugation, cryopreserved with 10% dimethyl sulfoxide, and stored in vapor-phase liquid nitrogen until the time of sample processing.

#### Cohort sample characteristics

All 14 patients had CML that was treated with CD6-T cell depleted allo-SCT. Of these, 12 patients had CML relapse after allo-SCT that was treated with CD8-depleted DLI, and 2 patients never had CML relapse and served as non-relapse controls (**Extended Data Table 1**). The median age of all samples was 23 years, ranging from 20-25 years. A median of 3 timepoints was available for each R and NR patient (range: 2-6), and there were no significant differences between R and NR cohorts regarding time from allo-SCT to DLI (R: median 702, range 362-2371 days; NR: median 1064, range 422-1787 days; *P*=0.6) (**Extended Data Fig. 1a**). Time from allo-SCT to sample for the non-relapsed controls was 1817 days for 5379 and 1113 days for 5380. Characteristics of samples are listed in **Extended Data Table 2**.

#### Cytogenetic and molecular information on CML tumor burden

The percent positivity of the Philadelphia (Ph) chromosome for each BM sample was extracted from the clinical record where available (as described previously^1^). Molecular remission was defined as achievement of molecular response (defined as the absence of BCR-ABL transcripts by RT-PCR). This data is shown in grey crosses in **Fig. 2f,g**.

#### Sample processing

Cryopreserved primary bone marrow mononuclear cells (BMMCs) were thawed on the day of sequencing at 37°C and dispensed drop-wise into a warmed solution of 10% FBS, 10% DNaseI (StemCell Technologies, cat. No. 07900) in PBS. The cell suspension was centrifuged at 200g for 10 minutes at room temperature. Viable cells were negatively selected using MACS Dead Cell Removal Kit (Miltenyi Biotec, cat. No. 130-090-101), running on MS columns to prevent sample loss. Collected live cells were resuspended in 0.04% BSA in PBS and diluted to a concentration of 1000 cells/uL. These cells were then divided into portions taken immediately for scRNA-seq (samples B1-B46) or for FACS isolation (described below) for subsequent ATAC-seq. For paired scTCR- and scRNA-seq on samples D1-D7 (**Suppl. Table 6**), BMMCs were processed as described here and then taken for FACS enrichment of T cells described below.

For cryopreserved PBMCs of DLI products (D8, D9; **Suppl. Table 6**), cells were thawed as described above, T cells were enriched using the human Pan T Cell Isolation Kit (Miltenyi Biotec), and then processed with the MACS Dead Cell Removal Kit (Miltenyi Biotec) before scRNA- and TCR-seq.

#### Fluorescence activated cell sorting (FACS)

For downstream ATAC-seq which was performed on samples B1-B46 (**Suppl. Table 5**), viable BMMC single-cell suspensions (prepared as above) were stained using antibody cocktails in the dark at 4°C, washed and run on a 5-laser FACSAria II (BD Biosciences) cell sorter. Cells then underwent FACS for the following CD14^-^CD19^-^CD3^+^ T cell populations: CD45RA^+^CD4^+^, CD45RA^-^CD4^+^, CD45RA^+^CD8^+^, and CD45RA^-^CD8^+^. The following fluorochrome-conjugated antibodies were used: CD14-FITC (M5E2, BD Biosciences); CD19-FITC (HIB19, BD Biosciences); CD3-PE (HIT3A, BD Biosciences); CD4-BUV395 (SK3, BD Biosciences); CD8-APC Vio770 (BW135/80, Miltenyi Biotec); CD45RA-BV510 (H100, BD Biosciences) (**Fig. S1**).

In order to perform paired scRNA- and scTCR-seq on samples D1-D7 (**Suppl. Table 6**), BMMCs were thawed as above without dead cell removal, stained with human Fc block (BD Pharmingen) for 10 minutes in the dark at 4°C, stained with antibody cocktail, washed and run on a 4-laser, FACSAria II (BD Biosciences) cell sorter. DAPI (BD Pharmingen) was used to exclude dead cells, and the following fluorochrome-conjugated antibodies were used to negatively select for T cells (to avoid stimulation of gene expression by anti-CD3 antibodies): *Lineage 1*: CD11c-FITC (B-ly6, BD Biosciences); CD14-FITC (M5E2, BD Biosciences); CD36-FITC (CB38, BD Biosciences); CD33-FITC (HIM3-4, BD Biosciences); CD16-FITC (3G8, BD Biosciences) *Lineage 2*: CD11b-PE (ICRF44, BD Biosciences); CD15-PE (HI98, BD Biosciences); CD34-PE (8G12, BD Biosciences); CD56-PE (B159, BD Biosciences); CD123-PE (7G3, BD Biosciences); CD235a-PE (GA-R2, BD Biosciences).

**Figure S1.**
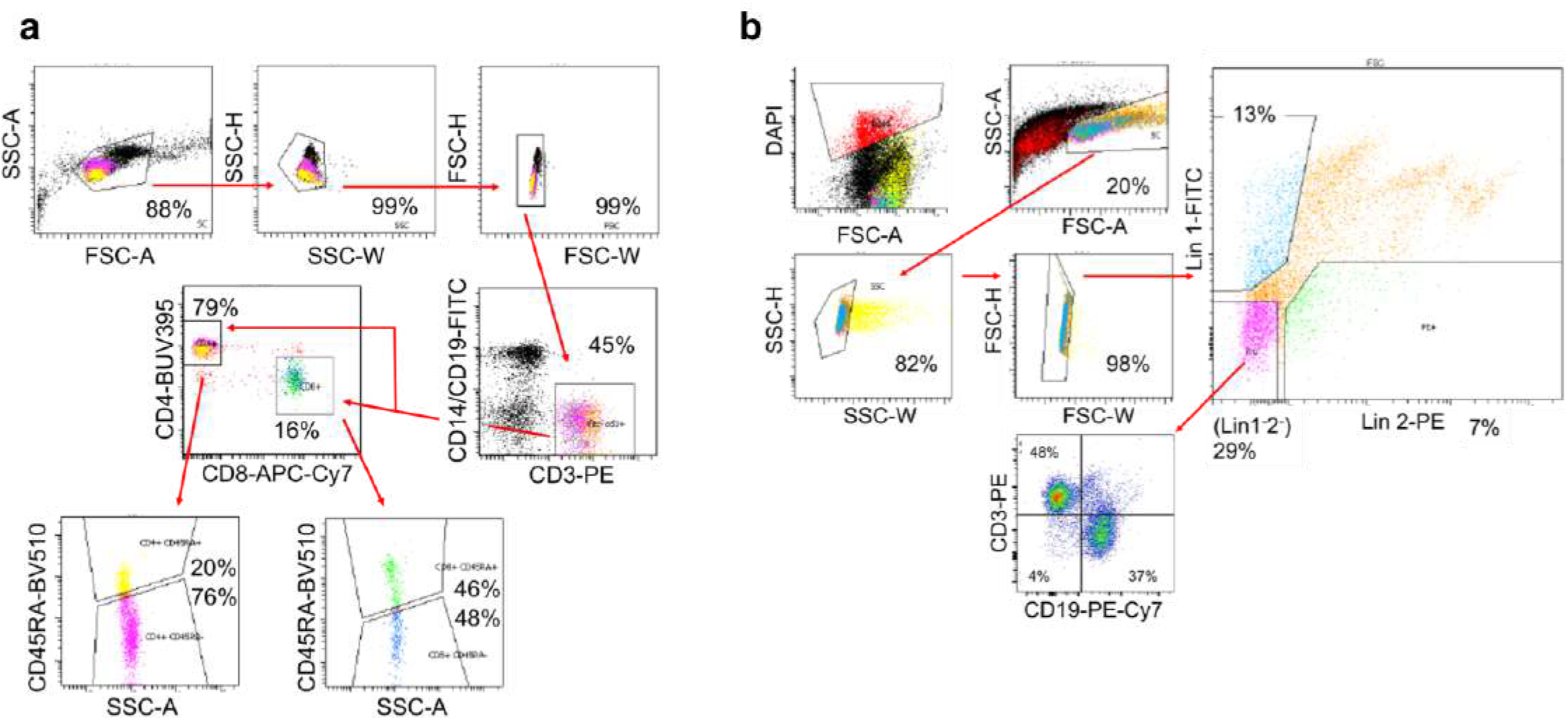
Gating strategy for sorting T cells from bone marrow mononuclear cells. **a**, Example BMMC sample shown of gating strategy used to isolate CD45RA^+^CD4^+^, CD45RA CD4^+^, CD45RA^+^CD8^+^, and CD45RA CD8’ T cell populations for ATAC-seq. **b**, Example BMMC sample shown of gating strategy for negative enrichment of T (and B) cell populations used for paired scRNA- and TCR-seq. Lineage 1 and 2 cocktails defined in text.

#### Library preparation for scRNA- and scTCR-seq

For BMMC samples B1-B46 (**Extended Data Table 2**), approximately 17,000 BMMCs (after dead cell removal) were loaded across 2 lanes onto a 10x Genomics Chromium™ instrument (10x Genomics) according to the manufacturer’s instructions. The scRNAseq libraries were processed using Chromium Single Cell 3’ Library & Gel Bead v2 Kit (10x Genomics). Quality control for amplified cDNA libraries and final sequencing libraries were performed using Bioanalyzer High Sensitivity DNA Kit (Agilent). scRNAseq libraries were normalized to 4nM concentration and pooled before loading onto Illumina sequencer. The pooled libraries were sequenced on the Illumina HiSeq X or NovaSeq S4 platform. The sequencing data were demultiplexed and processed as described below.

For BMMC samples processed for scRNA- and sc-TCRseq (D1-D7; **Suppl. Table 6**), 17,000 cells were loaded across two lanes onto a 10x Genomics Chromium™ instrument (10x Genomics) according to the manufacturer’s instructions. The scRNAseq libraries were processed using Chromium™ single cell 5’ library & gel bead kit and coupled scTCRseq libraries were obtained using Chromium™ single cell V(D)J enrichment kit (human T cell) (10x Genomics). Quality control for amplified cDNA libraries and final sequencing libraries were performed using Bioanalyzer High Sensitivity DNA Kit (Agilent). Both scRNAseq and scTCRseq libraries were normalized to 4nM concentration and pooled in a volume ratio of 4:1. The pooled libraries were sequenced on an Illumina NovaSeq S4 platform. The sequencing parameters were: Read 1 of 150bp, Read 2 of 150bp and Index 1 of 8bp. The scRNA- and TCR-seq data were processed as described in **Suppl. Note 4**.

#### Library preparation for ATAC-seq

After FACS isolation of CD45RA^+^CD4^+^, CD45RA^-^CD4^+^, CD45RA^+^CD8^+^, and CD45RA^-^CD8^+^ T cell populations, the Fast-ATAC protocol was then performed as previously described^2^. Briefly, fifty microliters of transposase mixture (25 μl of 2× TD buffer, 2.5 μl of TDE1, 0.5 μl of 1% digitonin, and 22 μl of nuclease-free water) (FC-121-1030, Illumina; G9441, Promega) was added to a cell pellet consisting of 10000-50000 cells and incubated at 37°C for 30 minutes. Transposed DNA was purified using a MinElute Reaction Cleanup kit (Qiagen), and purified DNA was eluted in 10 μl of elution buffer (10 mM Tris-HCl, pH 8). Libraries were barcoded (Nextera Index Kit, Illumina), amplified with NEBNext High Fidelity PCR Mix (New England Biolabs), and cleaned using a 1x volume of AMPure XP beads. Libraries were quantified using Agilent BioAnalyzer and sequenced on the HiSeq High Output and NovaSeq Illumina Sequencers (25 bp, paired-end).

### Supplementary Note 2: Single-cell RNA-seq data analysis

#### Preprocessing single-cell RNA-seq data

FASTQ files were preprocessed using the Sequence Quality Control (SEQC) bioinformatics pipeline^3^ with aligning reads to the hg38 genome and turning off the mitochondrial filter (using the option `--no-filter-mitochondriai-rna`). Characteristics of samples and quality control (QC) metrics are provided in **Extended Data Table 2**. In total, 381,462 total cells including 87,939 T cells (identified in the next section) from the combination of 41 bone marrow (BM) samples passed SEQC QC metrics, with a median of 2548 UMIs/cell and 8735 cells/sample.

#### Constructing global single cell map of T cells

##### Identifying T cells

To select T cells, we first normalized all *n* =381K BM cells to median library size and computed the log of normalized expression as 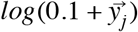 for each cell *j* = (1, …, *n*) where 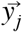 contains the normalized expression of genes in cell *j*. To identify major cell types, we filtered genes expressed in less than 2% of cells (resulting in 9767 genes) and performed PCA on the log-transformed normalized expression. The number of PCs was selected based on the knee-point (defined as minimum curvature radius) of eigenvalues. Then cells were clustered by applying Phenograph^4^ with the number of nearest neighbors set to 30, on the first 24 principal components (PCs), resulting in 94 clusters shown in **Fig. S2**.

**Figure S2.**
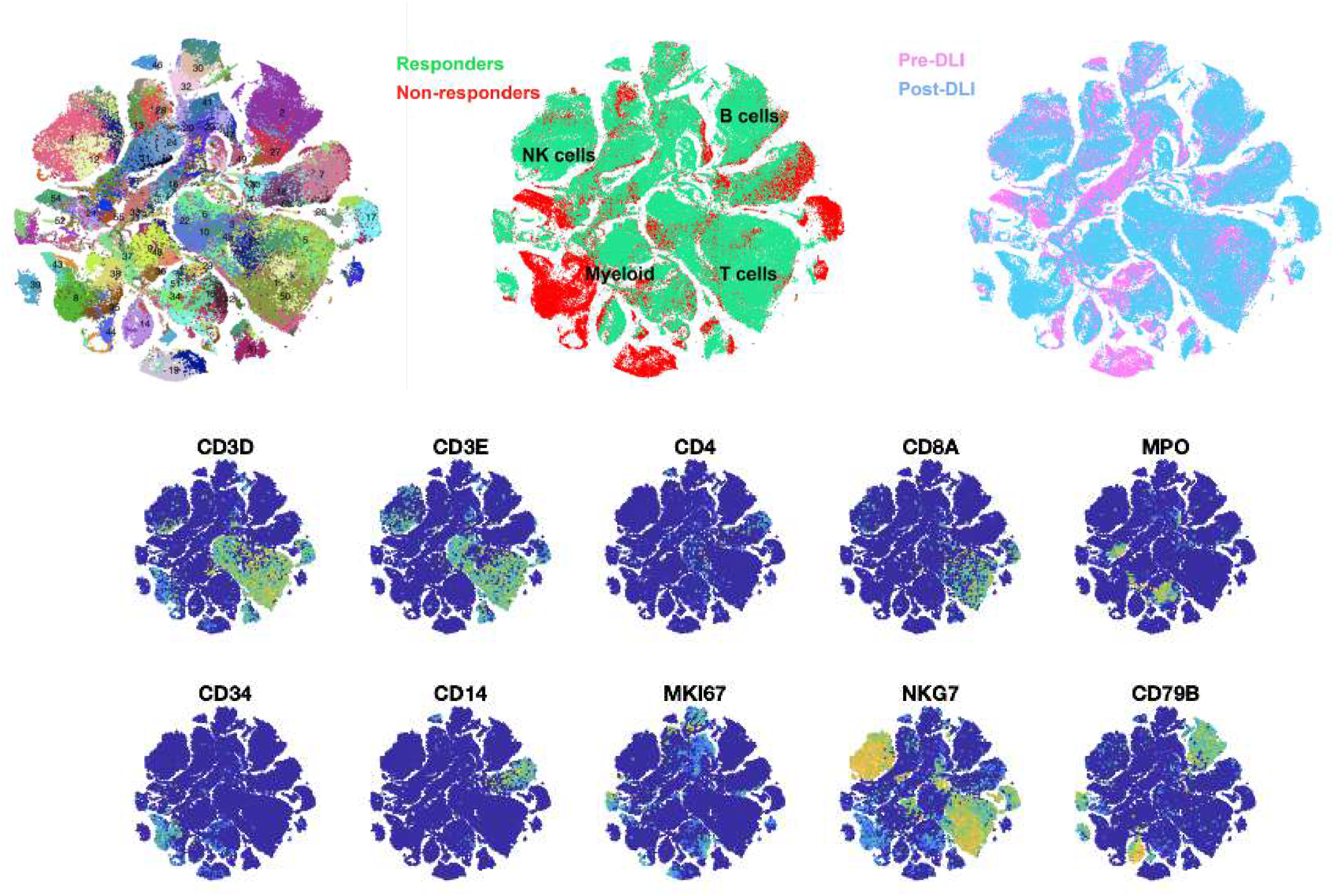
tSNE map of all transcriptomes from the leukemic microenvironment collected from 41 bone marrow samples colored by Phenograph cluster (top left), DLI outcome (top middle) and timing (top right) and markers of major cell types (bottom).

The normalized expression of {CD3D, CD3E} gene markers were averaged across cells in each Phenograph cluster and clusters with a high average expression of CD3 (right tail of distribution across all clusters) were selected as T cells, which consisted of 97,355 cells shown in **Fig. S3**.

**Figure S3.**
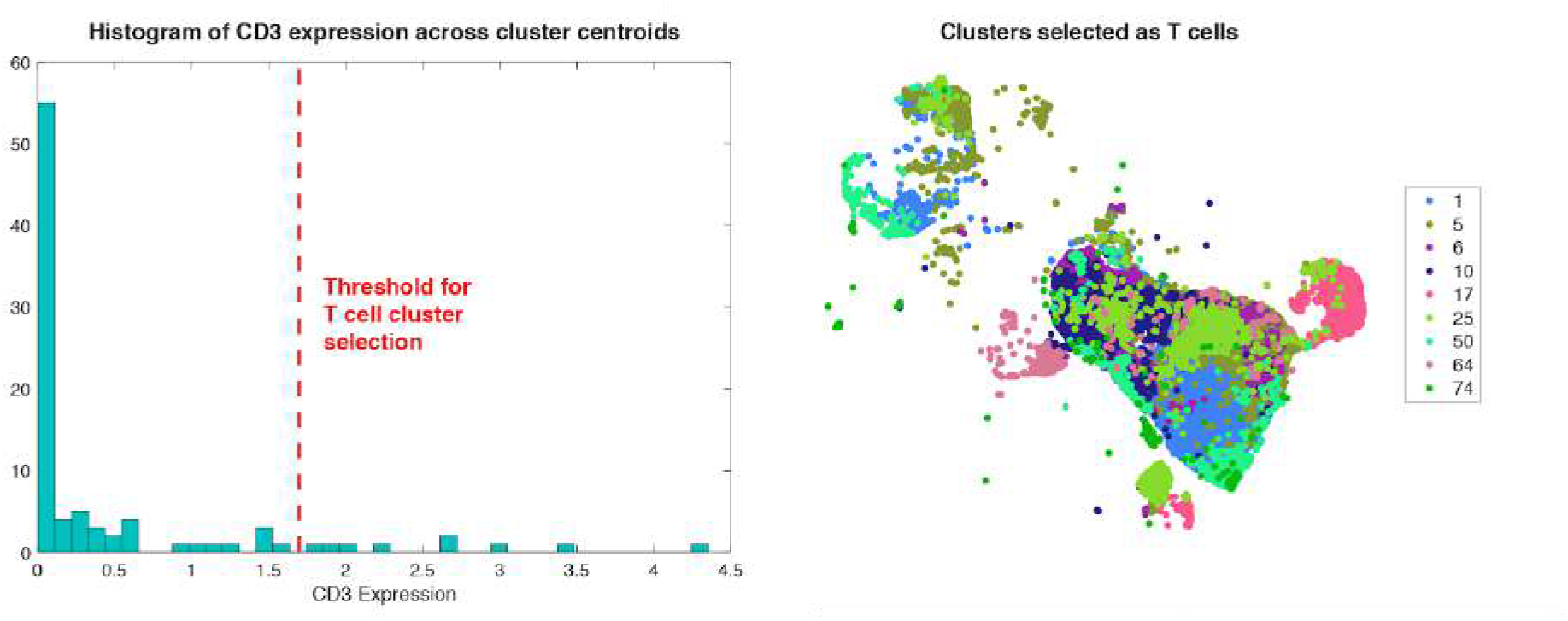
Histogram of expression of T cell markers across centroids of clusters (left) and subset of clusters selected as T cells marked (right) in the same tSNE map coordinates as in Fig. S2. These cells were then merged and further filtered for doublets and re-clustered as explained in the next sections for refined characterization of T cells.

##### Biscuit normalizing and clustering

To construct a more refined map of T cells, we performed simultaneous clustering and cluster-dependent normalization on raw counts for *n* = 97,355 T cells using Biscuit^3,5^. Using a hierarchical Dirichlet process mixture model, Biscuit performs a cell-type dependent normalization on the count matrix 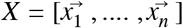 where each column 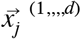 contains the expression (number of unique mRNA molecules) of *d* genes in cell *j*, while simultaneously inferring robust subsets of cells with *z_j_* denoting assignment of cell *j* to cluster *k*. Biscuit assumes that the log of counts 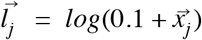 follow a multivariate Normal distribution: 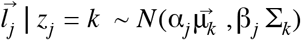 where 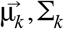 are the mean and covariance, respectively, of the *k*-th mixture component (cluster), and scalars α_*j*_, β_*j*_ are cell-dependent scaling factors used for normalization. We have previously shown that this cluster-dependent normalization removes batch effects while retaining biological signal^3^. In particular, Biscuit helps retain biological processes that are entwined with library size. For example in the case of immune cell activation, activated cells have a higher number of transcripts^6–9^ leading to higher total counts captured, hence variation due to real immune activation can be partially removed with methods that normalize cells by library size, whereas Biscuit performs a more careful normalization of cells conditioned on the cell state (captured by cluster assignment).

For faster inference, we used the implementation described in 3 (from https://github.com/sandhya212/BISCUIT_SingleCell_IMM_ICML_2016) which deploys a conjugate prior for the multivariate Gaussian, namely the Normal-inverse Wishart distribution for joint inference of cluster means and covariances.

After fitting the model, we transform the data from 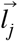 to 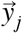 in which the expression is corrected for cell-specific factors α_*j*_, β_*j*_ using a linear transformation 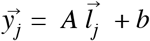 with 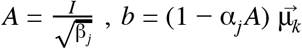 such that imputed expression for cell *j* follows *N*(μ_*k*_, Σ_*k*_) and hence all cells assigned to the same cluster follow the same distribution after correction.

Using Biscuit with 500 iterations; gene batch size set to 50, and alpha (dispersion parameter) set to 200, we identified 65 unique clusters. This choice of parameters led to both relatively good mixing of samples (**Fig. 1b** and **Extended Data Fig. 1e**), and distinct sets of differentially-expressed genes (**Extended Data Fig. 1c**). Only 3 clusters were found to be exclusive to one single patient (all 3 in NR 5326), who was the only patient with CML in blast crisis (**Extended Data Fig. 1e, Extended Data Table 1**).

**Extended data Fig 1e** shows the distribution of each cluster across clinical groups of R/NR and pre/post-DLI. Prior to computing the distribution, the number of cells in each cluster was first normalized by the total number of cells in each clinical group to account for imbalanced cell/sample numbers. The size of bubbles in each cluster is proportional to the distribution of normalized values and each cluster (column) sums to 100%.

Importantly, the interpretability of Biscuit enables the use of inferred parameters in downstream characterization of clusters: The inferred cluster mean μ_*k*_ and its conjugate prior 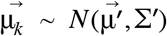 are used for estimating differentially expressed genes as detailed in the Cluster Annotation section below. To ensure each cluster is a legitimate cell population, we then scanned the clusters for doublets as explained below.

##### Removing doublets

Doublet cells were identified by applying DoubletDetection (https://github.com/dpeerlab/DoubletDetection), using the Biscuit derived clusters, with 50 iterations and `p_thresh=le-6, voter_thresh=0.8` followed by inspection of the co-occurrence of contradictory markers (including T cell and B cell markers; T cell and myeloid markers, T cell and erythroid markers etc). With this approach, 8.4% of cells were marked as doublets, which matches expectations given our cell loading (described in Suppl. Note 1). This resulted in 87,939 cells in 43 T cell clusters that were not flagged as doublets and retained for the remainder of the analysis.

##### Visualization

The Biscuit-normalized data for the 87,939 cells are projected to 2D in **Fig. 1b** and also expanded in **Fig. S4** using tSNE ^10,11^ on the first 18 PCs (identified based on knee-point of eigenvalues - defined as min curvature radius).

**Figure S4.**
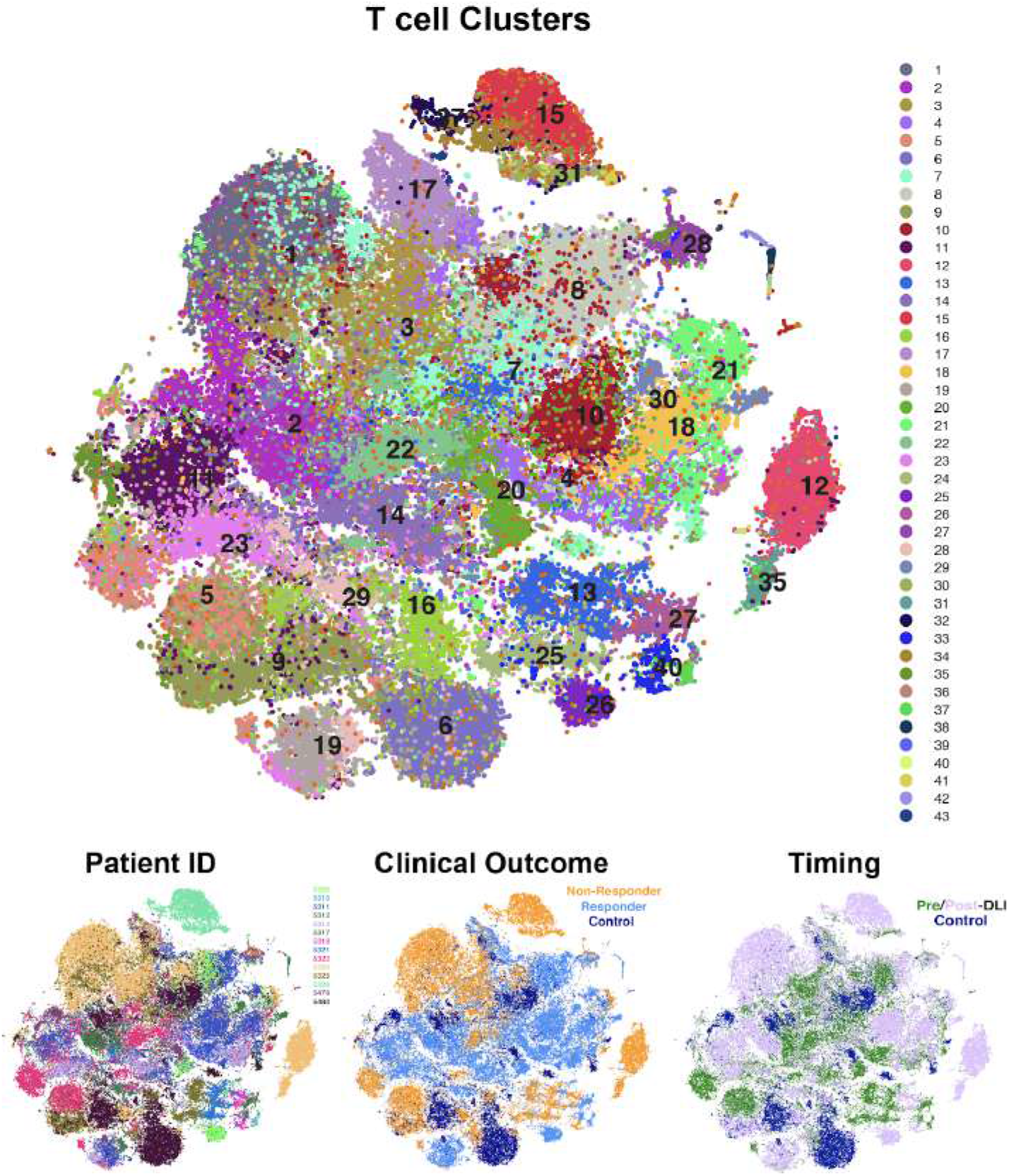
Expanded t-SNE projection of T cells (from Fig. 1b). Each dot represents a cell colored by cluster, patient ID, clinical outcome and timing respectively

##### Cluster annotation

T cell clusters were annotated through: (1) identifying cell type signatures enriched in each cluster (listed in **Suppl. Table 4**) by computing the expression of each signature (defined as average expression across all genes in a signature) per cluster and comparing to all other clusters using a t-test with p<0.1. The list of signatures compiled from literature are provided in **Supplementary Table 4**. The expression of enriched cell type signatures are shown in **Fig. 1c** and **Extended Data Fig. 1d**; (2) differentially expressed genes (DEGs) (**Extended Data Fig. 1c, Extended Data Fig. 3a**) were computed with t-test (p<0.01) comparing inferred mean expression of a gene in each cluster (listed in **Supplementary Table 2**) to its prior mean 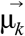 which represents expression across the entire population of cells. Since Biscuit fits a multivariate Gaussian mixture model to log-transformed data, the assumptions for a t-test are satisfied. **Extended Data Fig. 1c** shows the specificity of most DEGs to clusters as a block diagonal structure. The DEGs are listed in **Supplementary Table 3**.

The genesets derived from murine models of chronic viral infection^12^ were used for characterizing exhausted T cell subsets (**Fig. 2d)** listed in **Extended Data Table 3**. The T_EX_ and T_PEX_ score per cell was defined as normalized expression averaged across all genes in the geneset. Cell scores are aggregated by cluster in **Fig. 2d**.

For signatures related to T cell differentiation states (**Fig. 1c**, top), we used genesets from Gattinoni et al.^13^ To consider both up-regulated and down-regulated genes, we defined the expression of these signatures as a weighted sum of expression of genes in the geneset, with the weights being +1 or −1 for up-regulated and down-regulated genes respectively. We replaced *CD45RO* with the gene *HNRNPLL* gene which has been shown to regulate alternative splicing of CD45^14^.

#### Quantifying Diversity of T cell states

We evaluated if response to DLI was associated with a change in the number of distinct T cell transcriptional states. We found a marked increase in the number of T cell clusters in post-DLI samples compared to matched pre-DLI samples after controlling for cell number (t-test p-value <0.001). For this test, we corrected for differences in the number of cells. We downsampled each clinical group (R/NR, pre-/post-DLI, control) to 5000 cells by uniformly sampling with replacement from each group and clustering using Phenograph (using 10 PCs, K=30). This process was repeated 20 times and the number of clusters were compared with a t-test.

However, because T cell states are known to reside on continuous trajectories explaining the majority of variation^3,15,16^ we used the Phenotypic Volume metric devised in 3 to compare the global transcriptional diversity between clinical groups and before/after DLI.

Phenotypic volume (*V*) for a subpopulation of cells is defined as the determinant of the gene expression covariance matrix for that subpopulation, which considers covariance between all gene pairs in addition to their variance. The covariance matrix can be written as Σ^*d x d*^ and its pseudo-determinant *det* (Σ) is equal to the volume of a parallelepiped spanned by vectors of the covariance matrix ^17^ and can be computed as the product of nonzero eigenvalues of the covariance matrix. To improve sensitivity to noise and avoid multiplication of small nonzero eigenvalues, we compute the log of phenotypic volume which is the sum of log of non-zero eigenvalues:

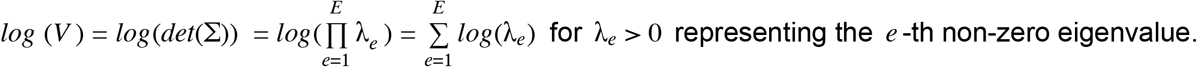

To correct for differences in number of cells, we downsampled each clinical group (R/NR, pre-/post-DLI, control) to 5000 cells by uniformly sampling with replacement from each group and computing the phenotypic volume. Only time points immediately pre-DLI and at remission post DLI (in Rs) were considered in this analysis. Patient 5321 was excluded in this analysis, as it did not have any post-DLI samples. **Table S1** shows the list of samples used in this analysis.

**Table S1.**
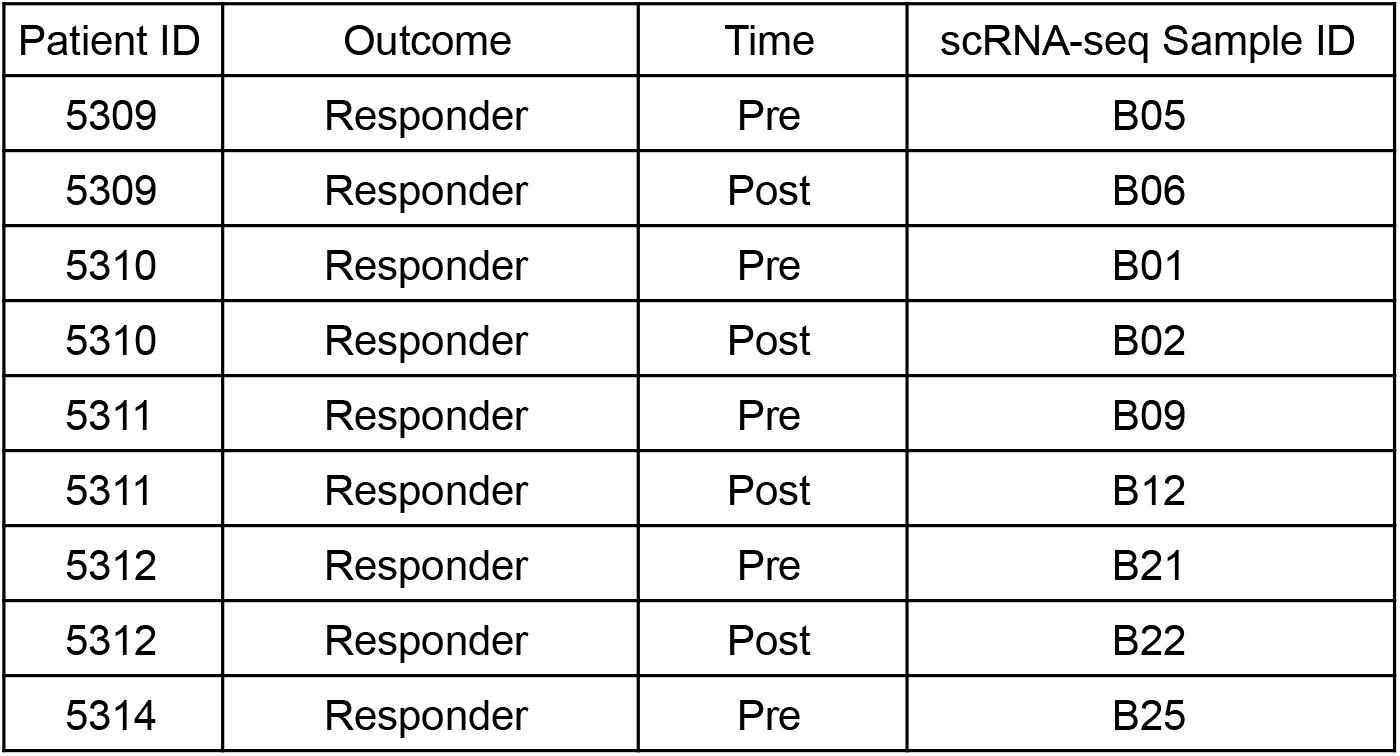
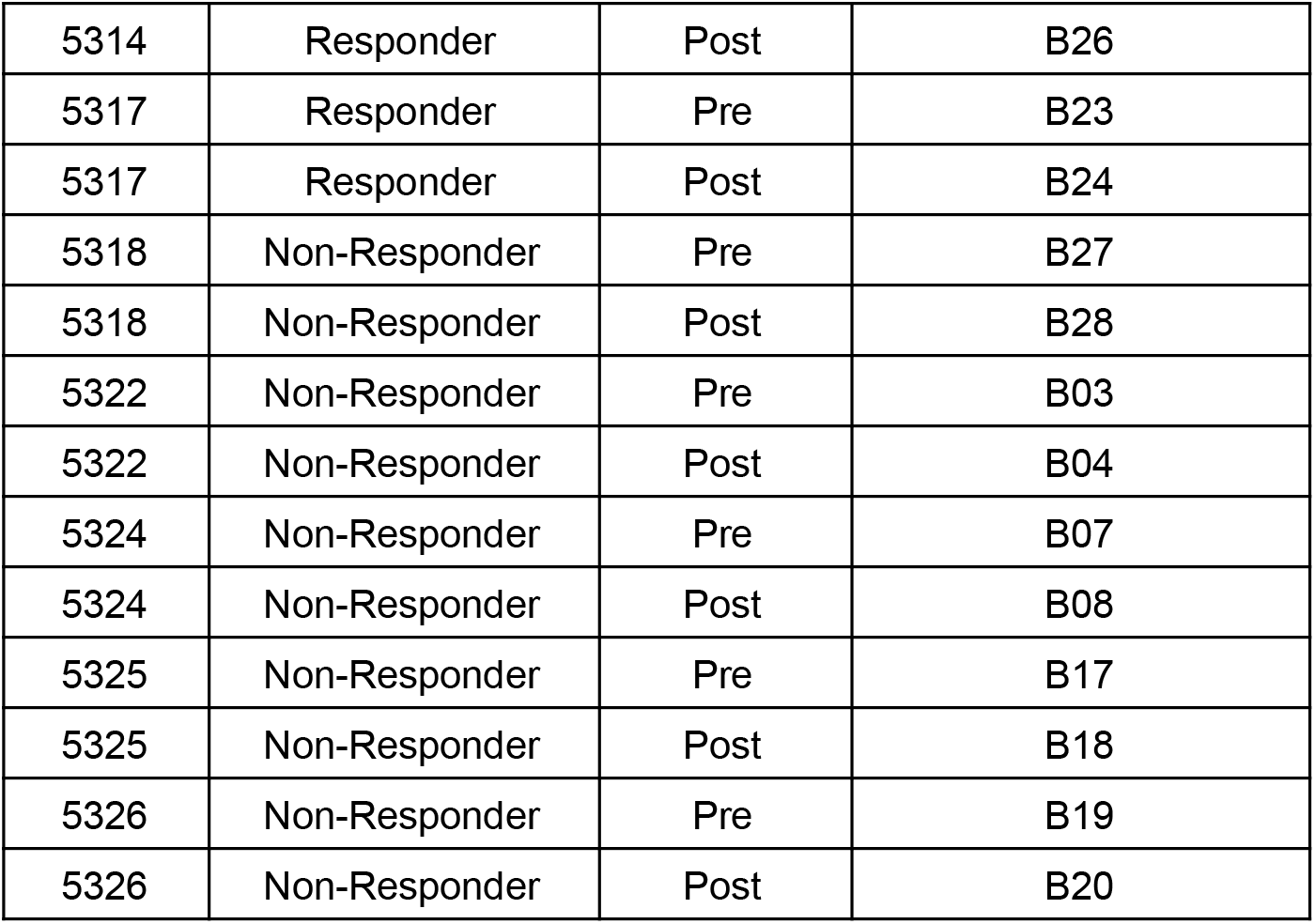
List of scRNA-seq sample IDs from baseline pre-DLI and the remission timepoint following DLI.

| Patient ID | Outcome | Time | scRNA-seq Sample ID |
| --- | --- | --- | --- |
| 5309 | Responder | Pre | B05 |
| 5309 | Responder | Post | B06 |
| 5310 | Responder | Pre | B01 |
| 5310 | Responder | Post | B02 |
| 5311 | Responder | Pre | B09 |
| 5311 | Responder | Post | B12 |
| 5312 | Responder | Pre | B21 |
| 5312 | Responder | Post | B22 |
| 5314 | Responder | Pre | B25 |
| 5314 | Responder | Post | B26 |
| 5317 | Responder | Pre | B23 |
| 5317 | Responder | Post | B24 |
| 5318 | Non-Responder | Pre | B27 |
| 5318 | Non-Responder | Post | B28 |
| 5322 | Non-Responder | Pre | B03 |
| 5322 | Non-Responder | Post | B04 |
| 5324 | Non-Responder | Pre | B07 |
| 5324 | Non-Responder | Post | B08 |
| 5325 | Non-Responder | Pre | B17 |
| 5325 | Non-Responder | Post | B18 |
| 5326 | Non-Responder | Pre | B19 |
| 5326 | Non-Responder | Post | B20 |

This process was repeated 50 times to achieve a range summarized in boxplots in **Fig. 2a, Extended Data Fig 3b** showing statistically significant expansion of volume after DLI in both Rs and NRs. Importantly, the phenotypic volume is higher in Rs compared to NRs in particular in baseline (pre-DLI). Both R and NR cases exhibited increases in phenotypic volume induced by DLI (log fold change=104.6, p<10^−6^). At both pre- and post-DLI timepoints, phenotypic volumes in R cases were higher than that of NR cases, (mean R-pre vs mean NR-pre, log-fold change = 199.1, p<10^−6^; mean R-post vs mean NR-post, log-fold change = 49.3, p=1.5×10^−6^), but a far greater increase in phenotypic volume was observed within NRs than within R’s (log-fold change [NR-post vs pre] = 203.8 vs log fold change [R-post vs pre) = 54.1; p<10^−6^].

Comparing the pre-DLI volume to that in non-relapse control samples in **Extended Data Fig 3b** reveals greater diversity of T cells in the leukemic microenvironment (in R/NR pre-DLI samples) than in non-relapse control samples which are leukemia-free. This increase in transcriptional diversity is similar to the expansion of phenotypic volume of T cells reported in the breast tumor microenvironment compared to normal tumor-free matched tissue^3^.

#### Common Factor Analysis

We aimed to decompose the T cells to uncover components potentially corresponding to response/resistance. The samples in Table S1 were used in this analysis. To correct for differences in numbers of cells across samples, we first downsampled T cells from each sample to 1000 cells, resulting in a total of 20,682 cells.

Applying PCA or diffusion component analysis^18,19^ showed that the top linear/nonlinear components explaining most of the variance across T cells are not highly correlated with response (**Extended data Fig. 2a**). Instead, we used Common Factor Analysis (CFA), a method that assumes there are underlying latent (unknown) factors that explain *shared* variance between cells, and thus explains *co-variation* of cells **Extended Data Fig. 2b** illustrates an example where cells are varying along two trajectories that could be related to different gene programs, e.g. T cell activation and exhaustion. If these trajectories are correlated but not colinear, dimensionality reductions methods that maximize explained variance will capture the two trajectories. CFA however will seek underlying (latent) factors that explain the shared variance between the two trajectories, ignoring the portion of variance unique to cells. Our assumption is that response or resistance might involve underlying latent factors associated with multiple distinct processes that might co-vary across the cells. Thus, common factors identified through CFA could potentially be related to response or resistance mechanisms affecting the majority of cells through multiple pathways (**Extended data Fig. 2b**). A brief description of CFA follows:

Shared factors are denoted as *f*_1_, *f*_2_, …, *f_m_* for expression of *n* cells denoted with *x*_1_,…, *x_n_*:

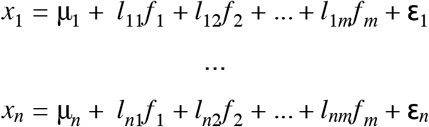

CFA assumes that *cov*(*f_i_,f_j_*) = 0 and *cov*(**ε**_*i*_, **ε**_*j*_) = 0 for *i ≠ j* and *cov*(**ε**_*i*_. *f_j_*) = 0.

Common factors were extracted using `factanal` function in R (https://www.rdocumentation.org/packages/FAiR/versions/0.2-0/topics/Factanal) with the method of maximum likelihood and `“varimax”` rotation. Setting the number of factors to two, a chi-square test rejected the hypothesis of model fit (p<0.05). Hence, we increased the number of factors to three which indicated that the hypothesis of perfect fit cannot be rejected. The first three common factors (**Fig. 1d.)** explain 67% of variance and separate groups of T cells enriched in Rs or NRs. To annotate the factors, we correlated the loadings of cells on each factor with expression of gene signatures. **Fig. 2e** shows gene signatures with the highest correlations with factors 1-3. **Extended Data Fig. 2d** shows that the signatures enriched for factors 2 and 3 are mostly non-overlapping, thus suggesting the involvement of different T cell dysfunction mechanisms in DLI resistance. Increasing the number of common factors to 4 and 5, we did not find any gene signatures highly correlated with the additional factors and factor 4 showed weak correlation with Hypoxia. We repeated this analysis on multiple downsampled sets and achieved the same conclusions with regard to signatures most correlated with factors.

#### Identifying T cell clusters enriched pre-therapy

We aimed to find any pre-DLI T cell states that are differentially enriched between Rs and NRs, that could potentially be predictive of response or resistance. Since different samples had differences in the total number of cells collected, this impacted our resolution of detecting a T cell state (cluster) in a patient. We therefore accounted for this uncertainty using a weighted one-sided t-test (using `statsmodels.stats.weightstats.ttest_ind` in Python). Within each clinical group (Rs or NRs), the weight of the *i*-th patient was given by: 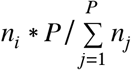 with *n_i_* denoting total number of T cells in patient *i* pre-DLI and *P* being the total number of patients in that group (R or NR).

We also corrected the p-values for the size of clusters using a bootstrapping technique: For each cluster *k* with size *u_k_*, we randomly select *u_k_* number of cells from the pool of all (R or NR) samples, and compute the p-value using the above test. Repeating this for 2000 iterations, we achieve a null hypothesis for p-values. The actual p-value for the cluster is then compared to the null, resulting in an empirical FDR (q-value) calculation. Applying this to pre-DLI samples shown in **Table S1**, we found clusters 4, 14, 21, and 27 were differentially enriched consistently across R patients compared to NRs (FDR<0.1) as shown in **Fig. 2b**. These clusters are enriched for T_EX_ gene signatures shown in **Fig. 2d,e**.

Aligned with our global observation with common factor analysis, we did not find any clusters to be differentially enriched consistently across NR patients compared to Rs, and we rather found multiple clusters each mostly present in one NR patient (**Extended data Fig. 1e**) suggesting that NR patients might be driven by different resistance mechanisms (**Fig 1e, Extended data Fig. 2d**).

#### Identifying T cell dynamics associated with therapy outcome

We used a weighted t-test similar to the previous section to compare the change in proportion of each cluster from pre-DLI to post-DLI. We performed a weighted one-sided t-test, summing the total cells in the pre- and post-DLI samples to determine the weights. Specifically, the expression we used for weights was:

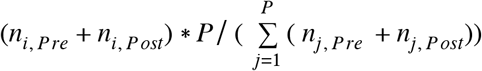

Where *n_i, Pre_* represents total number of cells in the pre-DLI sample of *i*-th patient and *n_i, Post_* represents the total number of cells in the post-DLI sample of *i*-th patient.

Compared to the test in the previous section which was performed on cluster proportions at one time-point (pre-DLI), this test involves computing the *change* in proportion from pre-DLI to post-DLI. Hence, the variance in the variable being tested is higher while the sample size (in this case number of patients) remains the same, meaning we have lower statistical power. In fact, across paired, pre- to post-DLI timepoints shown in Table S1, we found no single cluster to consistently expand or contract over time in Rs or NRs using the above weighted t-test. Thus, to improve our statistical power in detecting consistent changes in clusters over time, we combined clusters that are transcriptionally most similar as described below.

##### Defining meta-clusters

We computed the pairwise distance between each pair of clusters by comparing the distribution of expression of each gene across all cells in one cluster (from Biscuit normalized data) and comparing it to the distribution in another cluster using the Bhattacharyya distance metric ^20^, which is effective in pairwise comparisons of distributions. The advantage of computing cluster distances based on distribution is that we go beyond cluster means and also account for within-cluster variability, e.g. two clusters can have a similar mean expression but different variance. The total distance is then summarized across all genes, resulting in the distance matrix in **Extended Data Fig. 3c**. We then merged clusters that were most similar, resulting in 8 meta-clusters shown with white boxes.

##### Identifying expanding or contracting meta-clusters

By applying the weighted t-test above, we identified two metaclusters consistently expanded and one consistently contracted after DLI therapy (weighted t-test p<0.1), only in Rs, shown in **Fig. 2c**. The two expanding meta-clusters (M1 consisting of clusters {19,28} and M2 consisting of clusters {5,11,23}) are enriched for the Precursor Exhausted T cell gene signature T_PEX_ shown in violinplots in **Fig. 2d and Fig. S5**.

Interestingly, one expanding cluster (19 in M1) is also enriched in the non-relapse control samples (**Extended Data Fig 1e**), suggesting a transformation to normal T cell states after DLI in Rs. It should be noted that no meta-clusters or clusters consistently changed (expanding or contracting) in NRs, mirroring the *Anna Karenina* principle^21^.

**Figure S5.**
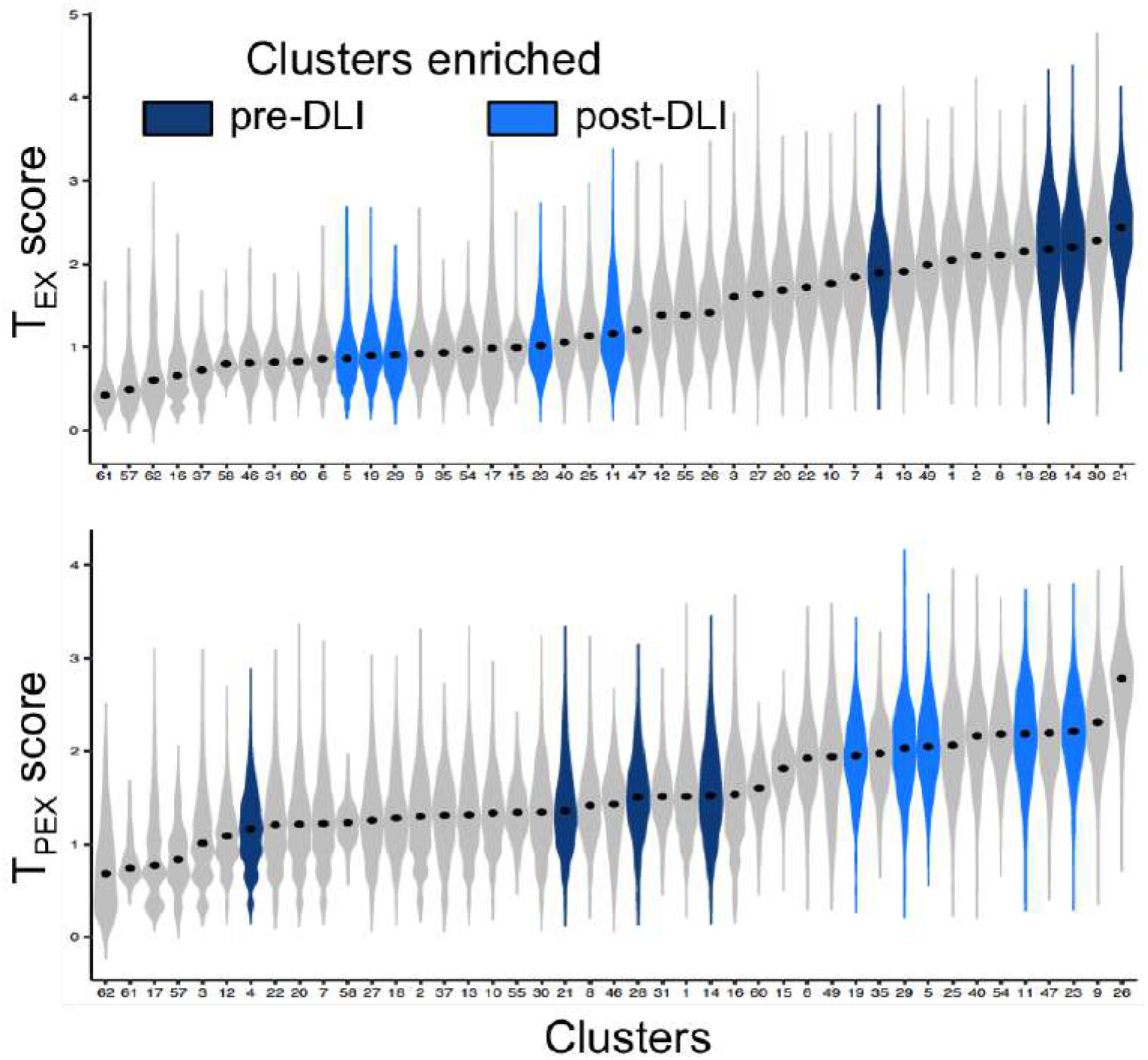
Expansion of Fig. 2d including all cluster IDs. Violin plots showing density of T_EX_ (top) or T_PEX_ (bottom) viral signature scores^23^ across T cells grouped by cluster. Clusters are ordered by median score. Colored violins refer to clusters enriched in pre-DLI Rs (dark blue) or expanding in post-DLI Rs (light blue).

#### Hierarchical Gaussian Process regression model

To study the dynamics of meta-clusters and tumor burden over time, we used a Gaussian Process (GP) model. The advantages of a GP model are (1) it is nonparametric, hence we do not assume a functional form over time and rather learn a distribution over all functions that explain temporal dynamics; (2) we account for dependencies between all pairs of time points which tackles the problem of non-uniform distribution of time-points in our cohort (**Fig. 1a**), for example in patient 5311, we have time-points within 19 days of each other, whereas in patient 5314 we have time-points 2.8 years (1059-29 days) apart from each other post-DLI and including them in the study can elucidate long-term sustainability of T cell states; (3) the probabilistic framework is flexible and we can therefore add priors representing uncertainty in measurements as explained below.

##### Tumor burden dynamics

We fit two GP regression models (*f_b_^R^, f_b_^NR^*), each with an Radial Basis Function (RBF) kernel^22^, to model the temporal changes in tumor burden in each outcome group (R or NR) separately in response to DLI therapy:

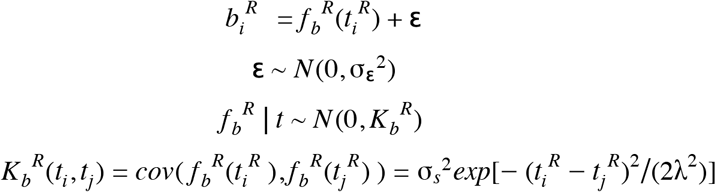

where *b_i_^R^* is tumor burden (see definition in section “Cytogenetic and molecular information on CML tumor burden” in Suppl. Note 1) in sample *i* in Rs and *t_i_^R^* is time relative to DLI therapy in sample *i* of Rs. Similarly for NR samples:

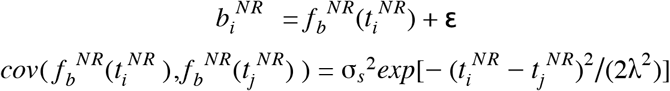

We optimized σ_*s*_ with the gradient-based algorithm Adam to maximize the log likelihood of our observed data. We set σ_ε_^2^ = 10 and λ = 285 (which is median distance between pairs of points). Results were robust to the choice of these parameters as shown in the next section.

Prior to regression, the mean tumor burden in each clinical group was subtracted so that our target variable *b_i_* would have zero mean, consistent with the distribution over *f_b_^RNR^*. This resulted in one model inferred for tumor burden in Rs (*f_b_^R^*) and one model for tumor burden in NRs (*f_b_^NR^*) shown in grey lines (mean) and shaded grey area (+/-1 standard deviation) in **Fig. 2f,g**. The data points for tumor burden are shown in grey crosses.

##### Temporal dynamics of T cell clusters

Similarly, we aimed to use a GP regression model to track the temporal dynamics of proportions of T cell meta-clusters in each outcome group. In other words, we learn two models *f_p_k__^R^, f_p_k__^NR^* on the *proportion* of each meta-cluster *k* over time separately in Rs and in NRs respectively. The proportion of a meta-cluster *k* in a sample *i* is defined as v*_i,k_ = m_i,k_/n_i_* with *m_ik_* being the number of cells in meta-cluster *k* in sample *i* and *n_i_* defined as sample size, i.e. total number of T cells in sample *i*.

Since there were significant differences in the size of samples and meta-clusters, we aimed to account for the uncertainty in detecting a metacluster in each sample (**Extended Data Fig. 3c)**. For example, if metacluster *k* is not observed in two samples *i*_1_ and *i*_2_ such that: v_*i*_1_, *k*_ = v_*i*_2_, *k*_ = 0, and sample *i*_1_ contains *n*_*i*_1__ = 10000 total cells compared to *n*_*i*_2__ = 1000 cells in sample *i*_2_, we have more certainty about the absence of metacluster *k* (representing a T cell state) in sample *i*_1_ than in sample *i*_2_ and the true value for v_*i*_2_, *k*_ could be missing or underestimated due to lack of statistical power.

To build this uncertainty into the probabilistic framework, we use a Gaussian process regression model that accounts for heteroscedastic noise. The measurement precision (β_*i*_) has a conjugate Gamma prior, whose mean is inversely proportional to the number of T cells measured in a given sample. Specifically we set the shape parameter of the prior distribution for β_*i*_ as *r* =1, and use the inverse of the number of cells collected for sample *i* as the rate parameter θ. This places more confidence on samples with larger sizes. For this model we use the RBF kernel *K*, with entries *K_ij_ = k*(*t_i_, t_j_*) and scale parameter σ_*s*_, set to the empirical variance of the response variable.

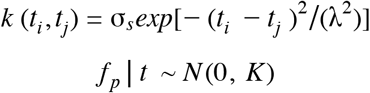

The full generative model is as follows:

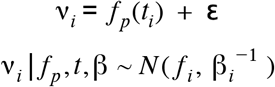

where:

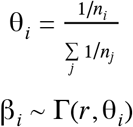

As with standard GP regression, after we fit our model to data *t* and *v*, we use the following joint marginal distribution to estimate the expected *v\** for an input *t\**. Specifically, let *k* = k*(*t_i_, t\**) be a vector representing the kernel function computed between each input time point *t_i_* in our training data, and our out of sample point *t\**, and let *c* = k*(*v*, v\**) be the kernel function computed on the out-of-sample time point. The joint distribution between our training data v and the new point v* is then as follows:

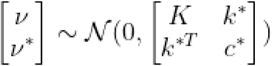

Because this is a multivariate normal distribution, we can use this distribution to compute the conditional distribution over *f_p_^*^* given our training data and *t\**:

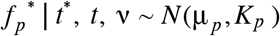

where the predicted mean and covariance are defined as follows:

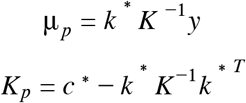

The plate model for this hierarchical GP model is shown in **Extended Data Fig. 3f**. We implemented this model in the probabilistic programming language pyro^23^ (https://pyro.ai/) and inferred the weights and temporal function with Stochastic Variational Inference, which computes an efficient approximation to the posterior by taking stochastic gradient steps to maximize the evidence lower bound (ELBO)^24^. The code for our hierarchical GP model is available at: https://github.com/dpeerlab/dli_gpr.

We first benchmarked this model on data simulated from a sinusoidal process *y* = 5*sin*(*x*) (shown as a grey line below) with two different noise variances representing levels of uncertainty in measurement: *y*_1_ = 5*sin*(*x*_1_) + **ε**_1_ with **ε**_1_ ~ *N*(0,1) (data points shown in blue) and *y*_2_ = 5*sin*(*x*_2_) + **ε**_2_ with **ε**_2_ ~ *N*(0,10) (data points shown in red) in **Fig. S6 (top)**. Please note the *y* notation here is not to be confused with expression in the Biscuit or Symphony models.

**Figure S6.**
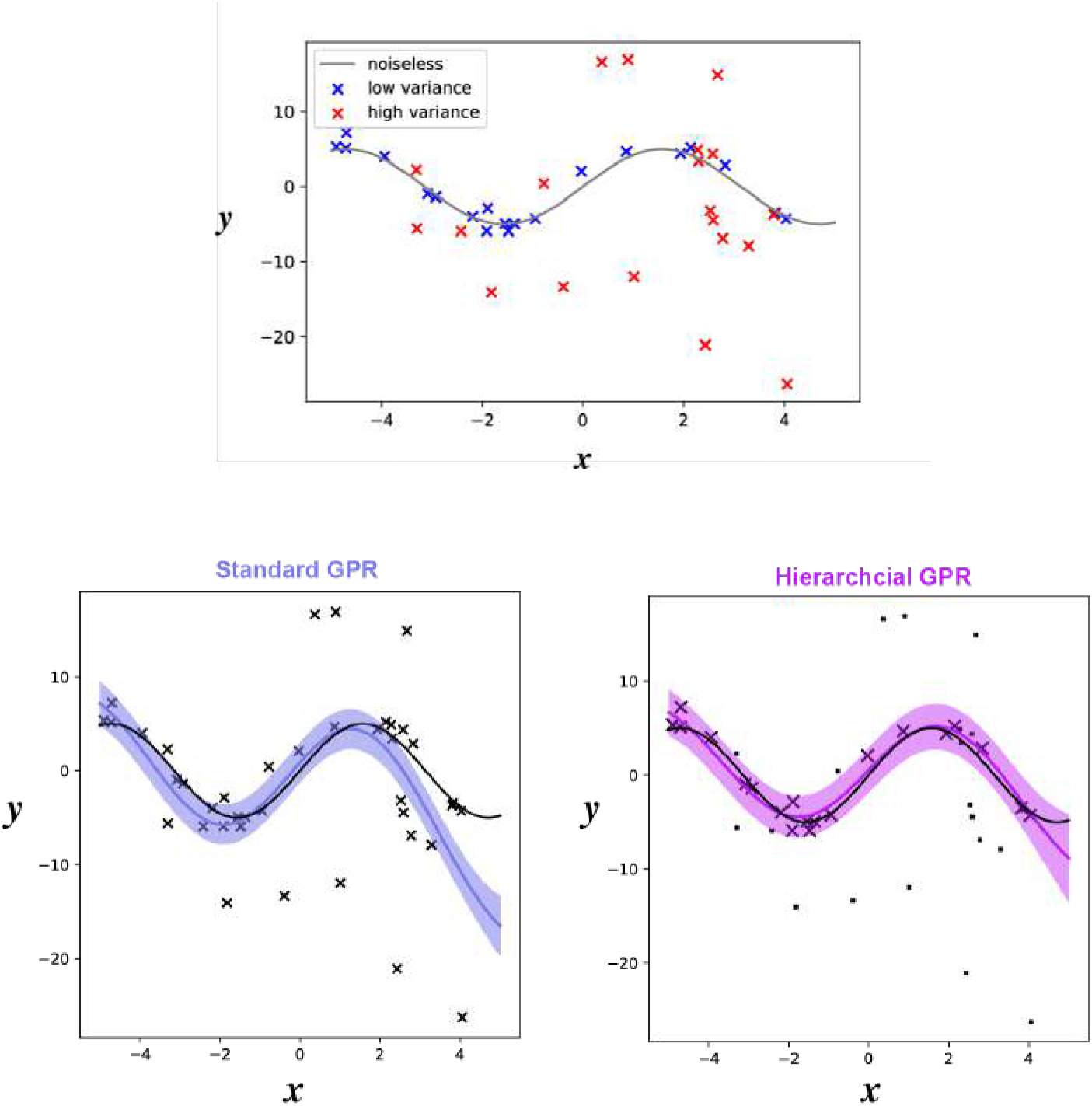
Top: Data simulated from GPR models built from a sinusoidal wave (grey line) with two different levels of noise variance. The data points are shown as red and blue crosses. Bottom: Standard GPR (left) and proposed hierarchical GPR model (right) fit to the simulated data. In the right figure, size of data points is inversely proportional to variance of generative model.

We combined these two datasets and fit the above hierarchical GPR model and compared it to the fit of a standard GPR (without prior) showing that the hierarchical model performs better in reconstructing the underlying sinusoidal function while a standard GPR model can overfit the noisy portion of data as shown in **Fig. S6** (bottom).

For quantitative comparison of the two models, we computed the log likelihood of unobserved noiseless simulated data along with the R2 score of the noiseless data vs. mean of the conditional distribution (Table S2).

**Table S2.** Performance of hierarchical GP on simulated data compared to standard GP regression.

| Model | Negative log likelihood | R2 score |
| --- | --- | --- |
| Hierarchical GPR | 193.24 | 0.801 |
| Standard GPR | 336.68 | 0.412 |

We then applied the hierarchical GPR mode to all meta-clusters in both Rs and NRs (**Fig 2f,g)** and use (T_EX_) metacluster M3 in Rs as an illustration. As reference, we compared the fit of the hierarchical model to a standard (vanilla) GP model (**Extended Data Fig 3g)**. The blue dots show the actual data points with the size of dots proportional to sample size *n_i_*. The blue line and shaded area shows mean and standard deviation of 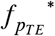.

Interestingly, the inferred hierarchical GP model shows that the T_EX_ meta-cluster tracks the tumor burden dynamics. The strong similarity between the inferred 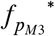 and *f_b_* in Rs is quantified by correlation, i.e. cross correlation at zero lag. M1 and M2 (T_PEX_) meta-clusters did not show a correlation with tumor burden (Table S3 and S4 below).

**Table S3.** Similarity between inferred GP model for metacluster proportion and model for tumor burden in Rs

| Metacluster | Correlation (Pearson R) at lag = 0 for Rs |
| --- | --- |
| M1 | -0.7436 |
| M2 | -0.2633 |
| M3 | 0.9852 |

We found that the dynamics of M3 do not follow tumor burden in NRs (Table S4 below).

**Table S4.** Similarity between inferred GP model for metacluster proportion and model for tumor burden in NRs

| Metacluster | Correlation (Pearson R) at lag = 0 for NR |
| --- | --- |
| M1 | 0.6907 |
| M2 | -0.7549 |
| M3 | -0.7009 |

Additionally, the expansion of T_PEX_ clusters post-DLI is durable in Rs and nonexistent in NRs. Results were robust to choice of of σ_ε_ and λ. As shown in **Fig. S7**, similar fit is achieved on a range of values. This example shows tumor burden and proportion of T_EX_ metacluster M3 in non-responders.

**Figure S7.**
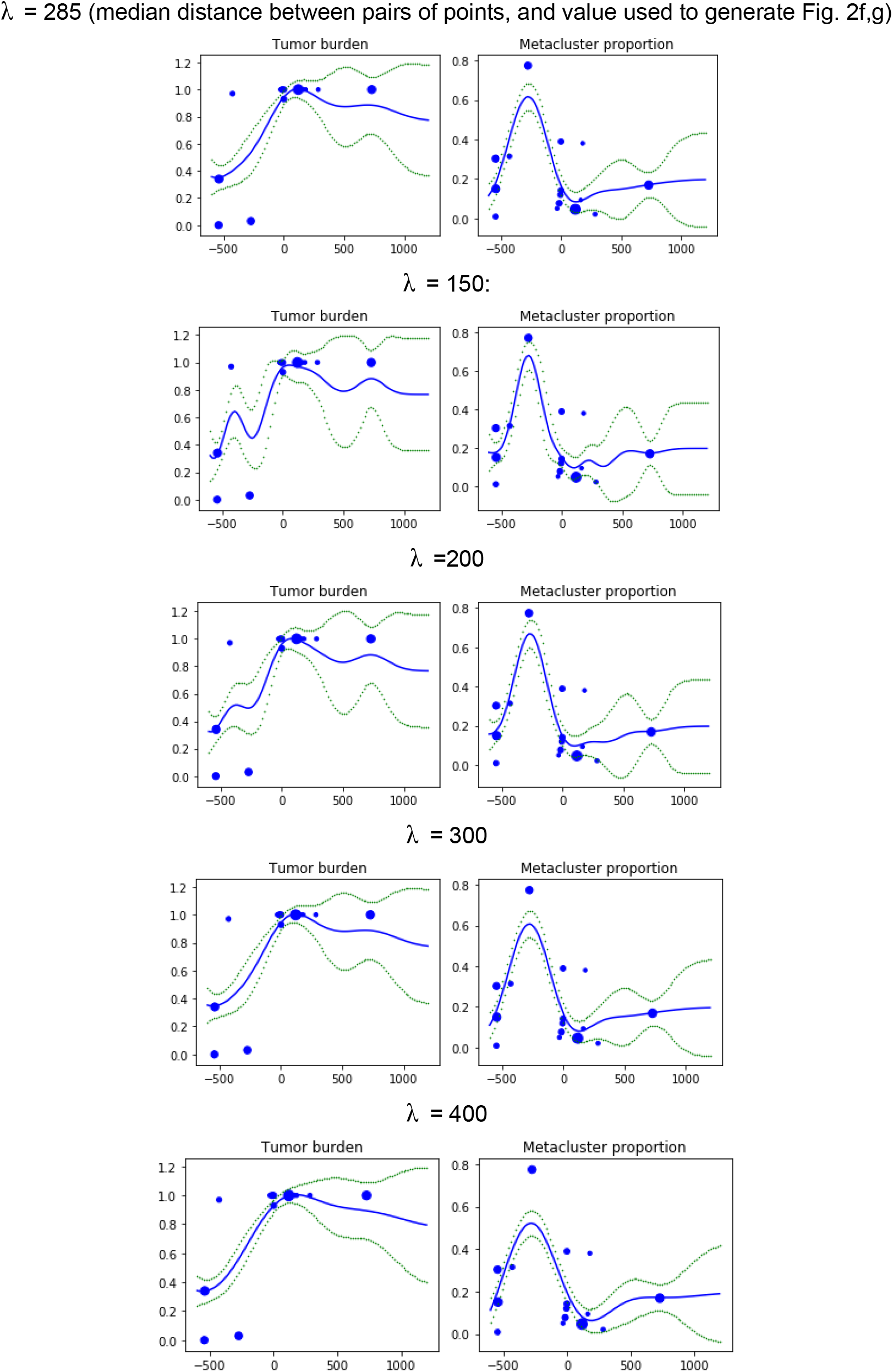
Inferred hierarchical GP models showing robustness to choice of λ; each dot is a sample and size of dots are proportional to total number of cells in the sample; x-axis is time from DLI and y-axis is tumor burden (left) and proportion of cells in metacluster M3 (right) from each sample in NRs.

To quantify the relative timing of T_EX_ and T_PEX_ meta-clusters, we computed the cross-correlation between *f_p_*\* and *f_b_* shown as purple bars in **Fig. 2f** (middle row). The max cross-correlation between M3 and tumor burden in Rs 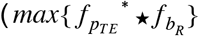 with ★ indicating cross-correlation) is at 75 days which is 1/4 of median time interval between samples (marked with a red line in **Fig. 2f** left middle; t-statistic=8.58, p=0) indicating they are in sync, whereas for the T_PEX_ meta-clusters, 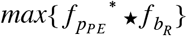 occurs at 703 days (M1: t-statistic=2.05, p=0.02; M2: t-statistic=0.72, p=0.23) indicating a significant lag compared to the T_EX_ M3 meta-cluster and tumor burden.

### Supplementary Note 3: Integration of single-cell RNA-seq and ATAC-seq

#### Preprocessing ATAC-seq data

Bulk ATAC-seq data for each sorted subset of T cells from each bone marrow sample was processed using the automated end-to-end quality control (QC) and processing pipeline (https://github.com/kundajelab/atac_dnase_pipelines) from the ENCODE consortium with configuration `SPECIES=hg38`. Alignment is performed using Bowtie2^25^ and peak calling and normalization is done with MACS2^26^. MACS2 normalization involves comparing ATAC signal to local background noise using a Poisson test^26,27^. The full list of samples and QC metrics for ATAC-seq data are provided in **Suppl. Table 5**.

#### Correlation between accessibility profiles

We first aimed to study the potential impact of DLI in the global epigenetic landscape of T cells. We thus compared ATAC-seq samples from the same time-points as in Table S1, with ATAC-seq ID listed below in Table S5. To compare chromatin accessibility between pairs of samples, we first created a consensus peak set similar to Corces et al 2016^2^ as follows: Peak summits were extended to 150bp windows and a set of maximally non-overlapping peaks was generated across all samples, resulting in 133,968 peaks for CD8+ CD45RO+ and 169,740 peaks for CD8+ CD45RA+ samples. Then Pearson correlation was computed between all pairs of 14 samples in each subset, and then correlations were averaged by pairs of clinical groups (**Fig. 3b**).

**Table S5.** List of ATAC-seq samples from IDs from baseline pre-DLI and the remission timepoint following DLI; n/a denotes low sample quality or excluded based on data preprocessing QC.

| Patient | Outcome | Time | ATAC-seq sample ID for CD8 CD45RA sorted T cells | ATAC-seq sample ID for CD8 CD45RA sorted T cells |
| --- | --- | --- | --- | --- |
| 5309 | Responder | Pre | C44 | C45 |
| 5309 | Responder | Post | n/a | n/a |
| 5310 | Responder | Pre | C31 | C32 |
| 5310 | Responder | Post | C35 | C36 |
| 5311 | Responder | Pre | C63 | C66 |
| 5311 | Responder | Post | C79 | C81 |
| 5312 | Responder | Pre | C105, C106 | n/a |
| 5312 | Responder | Post | n/a | n/a |
| 5314 | Responder | Pre | n/a | n/a |
| 5314 | Responder | Post | n/a | C130 |
| 5317 | Responder | Pre | n/a | C114 |
| 5317 | Responder | Post | C118 | C119 |
| 5318 | Non-Responder | Pre | C133, C134 | n/a |
| 5318 | Non-Responder | Post | n/a | C156 |
| 5322 | Non-Responder | Pre | C38 | C39 |
| 5322 | Non-Responder | Post | C41 | C42 |
| 5324 | Non-Responder | Pre | C48 | C49 |
| 5324 | Non-Responder | Post | C54 | C56 |
| 5325 | Non-Responder | Pre | C91 | C92 |
| 5325 | Non-Responder | Post | C95 | n/a |
| 5326 | Non-Responder | Pre | n/a | n/a |
| 5326 | Non-Responder | Post | n/a | n/a |

#### Symphony model for cell type-specific gene regulatory networks

To study the underlying circuitry of distinct clusters, we developed a novel integrative model named *Symphony*^28^, for inferring gene regulatory networks (GRNs) specific to subsets of cells.

Gene regulatory networks (GRNs) are directed weighted networks between genes depicting the extent to which a regulator gene influences (activation or repression) the expression of each of its downstream target genes. Symphony estimates these networks in each subset by extracting co-expression patterns between TFs and target genes from scRNA-seq and combining them with the presence of TF motifs within regions of chromatin accessibility in the vicinity of targets as derived from ATAC-seq. This is accomplished in Symphony by constructing a generative model that mimics transcriptional regulation illustrated in **Extended Data Fig 4b**.

Since the ATAC-seq data in this study measures accessibility summarized across all cells in a sorted compartment (e.g. CD8+CD45RO+) each consisting of multiple T_TEX_ or T_PEX_ clusters, we also leveraged the deconvolution capability of Symphony: bulk epigenetic data is deconvolved into cluster-specific epigenetic profiles. The deconvolved profiles are then used to explain gene co-expression patterns through GRNs, and thus resolve direct links from indirect links in the network (**Extended Data Fig 4b**).

Symphony^28^ is an extension of the Biscuit^3,5^ model which clusters cells while simultaneously distinguishing biological heterogeneity from technical noise in single-cell gene expression data (also explained in **Suppl. Note 2**). Symphony extends this model by replacing the hyperparameter for gene co-expression in Biscuit with a generative process exclusively driven by epigenetic data (collected from the same sample or a sample with similar composition of cell types). Thus, Symphony models the biological mechanism responsible for the observed gene co-expressions per cell type.

The model also simultaneously deconvolves the bulk epigenetic profiles (which denote accessible DNA) into cell-type (cluster)-specific accessible regions (**Extended Data Fig 4b**) within a unified statistical framework. Within these regions, the binding of transcription factors (TF associated with open regions based on known DNA binding motifs) impacts the expression of nearby genes, such that accessible regions may help explain gene-gene interactions.

Given the observed bulk chromatin accessibility profiles and single-cell RNA-seq count matrix, the model finds a deconvolution of the bulk accessibility data into cluster-specific accessibility profiles that are best able to explain the gene-gene relationships observed in scRNA-seq. We note that Symphony can infer whether a TF impacts a target gene without requiring epigenetic evidence as well, which facilitates inferring the regulatory influence of the many TFs (e.g. *TOX*) for which a binding motif is unknown.

Symphony input, output and model specification are provided below:

##### Input data to Symphony

The observed paired datasets are:

1. Epigenetic data measured with ATAC-seq^29^, denoted as *C*^*w*×*r*^ = [*c*_1_, …, *c_t_* …, *c_r_*] where 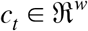 is epigenetic data for one patient (as replicate), containing accessibility (quantified as peak height) in genomic regions *m* = [1, …, *w*] (identified from MACS2^26^).
2. Single-cell RNA-seq data 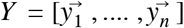 where 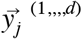 denotes log-transformed normalized single-cell expression data for cell *j* with *d* genes.

##### Symphony output

The main latent variables being estimated (**Extended Data Fig 4b**) are:

1. Epigenetic profile for each cluster *k* represented as 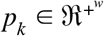 which contains estimated genome accessibility in *w* genomic regions.
2. Gene Regulatory Network (GRN) represented as *R_k_* for each cluster *k*. 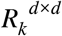 is an asymmetric matrix with nonzero entries 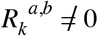 if gene *b* is predicted to be regulated by gene *a*. Positive and negative values for 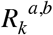 suggest activation and repression respectively.

###### Model details

These latent parameters are estimated simultaneously in an integrative model with three components explained below:

###### Epigenetic model

Bulk epigenetic profiles (*c_t_*) are assumed to be represented as a weighted sum of cluster-specific epigenetic profiles (*p_k_*) such that:

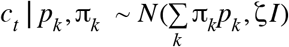

where the weights π_*k*_ represent the proportion of clusters in the sample. This assumption is validated in ^28^ using data on PBMCs with ground truth deconvolved profiles.

We set a Gamma prior for accessibility: *p_k_* ~ *Gamma*(η, Λ) to ensure a positive domain.

###### GRN model

We assume a regulatory link is dependent on genome accessibility as well as motif information within an accessible region. Specifically, a genomic region *m* in *C* is mapped to an interaction between genes *a*, *b* in *Y* with a predefined function *g*(*a*, *b*) = *m*. We also define *M*^*d*×*d*^ based on prior knowledge: *M_a,b_* = 1 if the motif sequence for gene *a* exists in region *m* in the vicinity of gene *b*, suggesting a potential regulatory interaction from gene *a* to gene *b*. Motifs were scanned using FIMO^30^ in this study.

We thus model 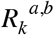 as:

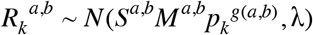

Where *S* is a sign indicator representing activation or repression set according to the sign of empirical covariance:

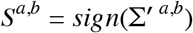

Σ′^*a*,*b*^ is an empirical prior set to the covariance between genes *a*, *b* across all cells in the scRNA-seq data. The variance *λ* allows for 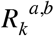 to have non-zero value, even when *M_a,b_* = 0.

###### Expression model

Similar to Biscuit^3,5^, Symphony assumes that log-transformed normalized single-cell expression data follows a multivariate Normal distribution:

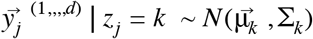

where *z_j_* denotes the assignment of cell *j* to cluster *k* modeled as:

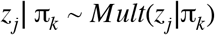

Since the single cell expression data was already normalized and clustered with Biscuit as explained in **Suppl. Note 2**, we did not use the clustering feature of Symphony and instead fixed the assignments (*z_j_*) of cells to clusters as assigned by Biscuit; the proportions π_*k*_ are thus also fixed. The full normalized expression matrix 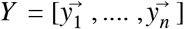 (output from Biscuit) is thus used as the second input to Symphony in this case. However, as a more general tool Symphony is also able to successfully cluster de-novo as demonstrated in simulated data^28^.

The parameters 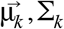 are the mean and covariance, respectively, of the *k*-th cluster. We define the prior for 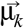 in Symphony as follows:

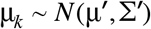

where μ′ is set to the empirical mean expression across all cells and Σ′ was set to *I* (identity) in this study.

Importantly, the covariance in observed gene expression is related to a graph power of the regulatory network, capturing the propagated impact of regulation in the network (indirect regulation) as depicted in **Fig. S8** below. Specifically, co-expressed in each cluster is modeled as:

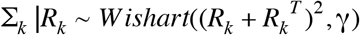

While using a Wishart instead of Inverse Wishart is not conjugate, this is valid as both distributions satisfy the positive semi-definite requirements for priors on the covariance matrix.

**Figure S8.**
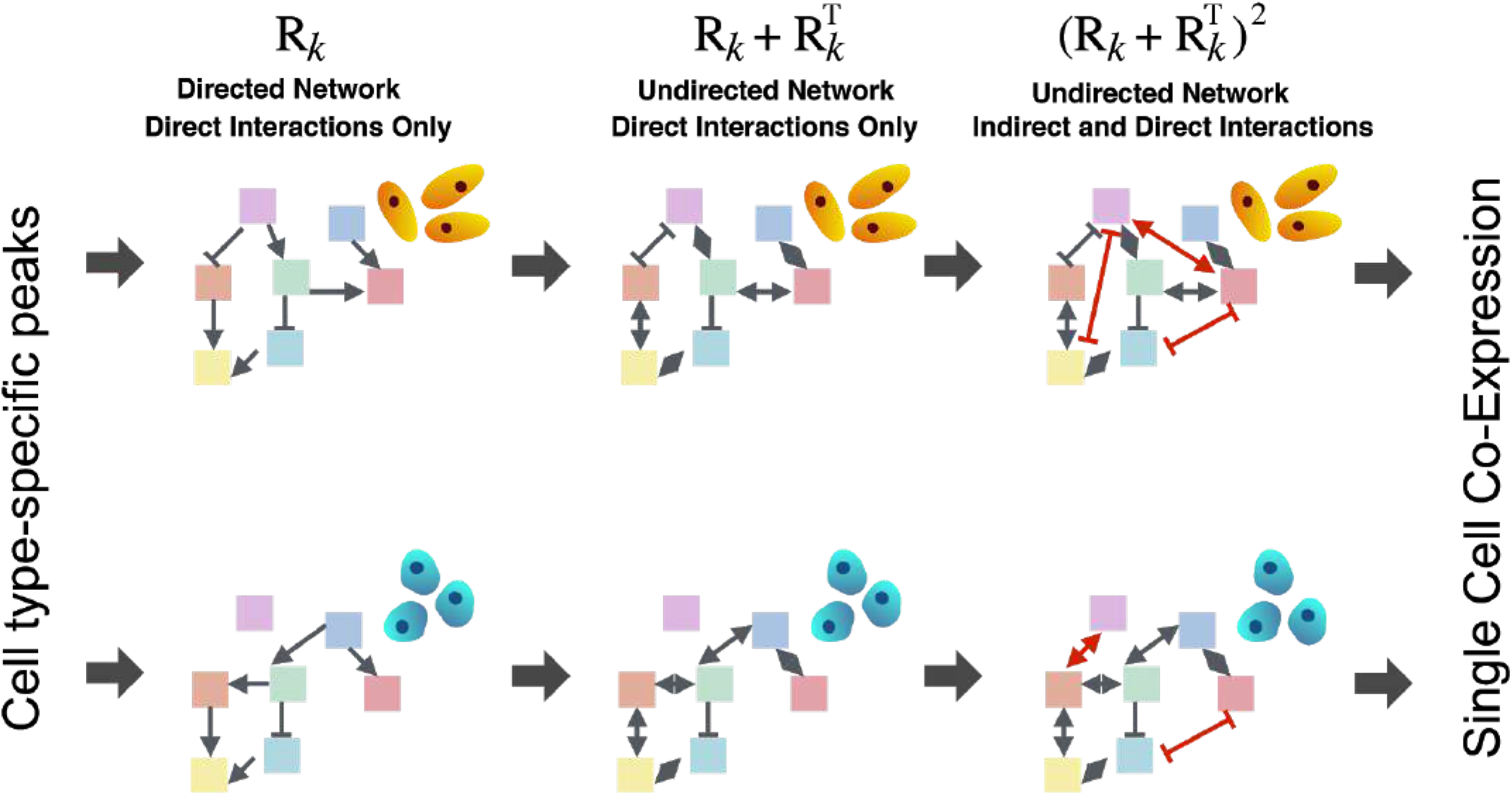
Symphony captures direct and indirect regulation. The impact of regulation is propagated through the network up to path length of two and is reflected in covariance between indirectly connected genes^28^.

The plate model for Symphony used in this study is shown below in **Fig. S9**.

**Figure S9.**
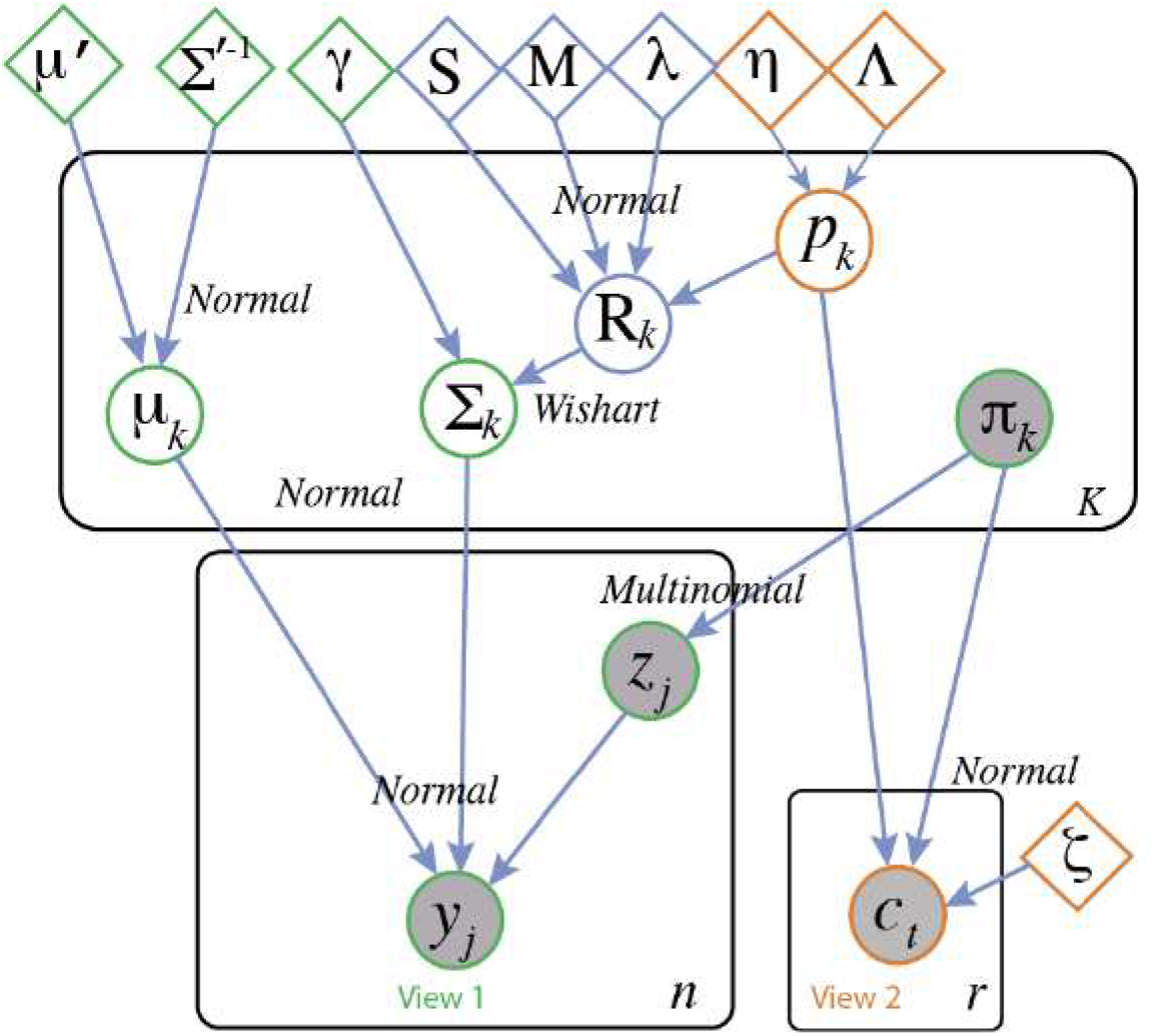
Plate model for Symphony^28^ with fixed cluster assignments used in this study

###### Inference, approximations and scalable implementation

An EM-VI inference procedure was presented for Symphony in ^28^. We also showed the performance of Symphony on well-characterized peripheral blood mononuclear cells (PBMCs), and significant improvement over other deconvolution methods^28^. In this study, given the complexity of the model and size of data, we used a scalable implementation of Symphony in the probabilistic programming language Edward^31^. This implementation in Edward is provided in https://github.com/dpeerlab/Symphony with input data for group 1 accessible for reviewers.

The use of variational algorithms in Edward^31^ allows for fast approximations of the posterior for large gene-by-gene matrices including GRNs and covariances per cluster, and scales well to additional cells and ATAC-seq replicates. Setting constraints on covariance matrices of a multivariate normal distribution are difficult to enforce in the optimization setting of variational inference. Thus, to avoid non-singularity issues during optimization, we define the Wishart distribution in Edward using the Bartlett Decomposition, rather than the built-in Wishart function of tensorflow, which allows us to more easily define variational parameters.

Specifically, we replace the sampling of covariance matrices ∑_*k*_ |*R_k_* ~ *Wishart* with a generative model constructed from univariate chi-squared distributions and normal distributions, which can be shown to produce a valid sample from the Wishart distribution^32^. Given *L_k_* as a cholesky factor of the prior (*R_k_* + *R_k_^T^*)^2^, we sample the cluster-specific covariance as follows:

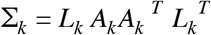

where *A_k_* is a lower triangular matrix whose diagonal elements are composed of χ^2^ random variables with *γ* – *i* +1 degrees of freedom, where *i* indexes the rows of *A_k_*, and the off-diagonal elements in the lower triangle are independent normal distributions. Hence each Σ_*k*_ is a positive semi-definite matrix centered at *L_k_L_k_^T^* or equivalently (*R_k_* + *R_k_^T^*)^2^. In this setting, we define variational distributions corresponding to the dummy variables *h* ~ *chi squared* and *v* ~ *Normal*, as opposed to defining a matrix variate distribution which, during the course of optimization, must fit all the constraints of valid covariance matrices.

Still, in the Edward implementation, we observed that the Barlett product often produced matrices which are not positive semi-definite due to numerical instability, and hence did not generate a valid covariance matrix. As such, we approximated the mean of Σ_*k*_ with the highly-related (unitarily similar) matrix *L_k_^T^L_k_*, which we ensured produced a posterior in covariance which is highly correlated with its mean derived from the posterior GRNs (minimum correlation r=0.745 across all groups in this paper). For additional speed, the cholesky factor was computed from Gram matrix (*R_k_* + *R_k_^T^*)^2^ using the QR decomposition of (*R_k_* + *R_k_^T^*) where *R_k_*′ = *L_k_^T^* given *Q_k_*′*R_k_*′ = *QR*(*R_k_* + *R_k_^T^*).

In addition to the use of the Bartlett Decomposition, the Edward version of Symphony replaces the standard Wishart with a scaled Wishart for added flexibility of the model in the variational inference case. The scaled Wishart necessitates addition of a latent parameter per cluster δ_*k*_, such that *Σ_k_*′ ~ *Wishart* and δ_*k*_ ~ *Normal* and Σ_*k*_ = Δ_*k*_ Σ_*k*_′ Δ_*k*_ where *diag*(Δ_*k*_) = δ_*k*_.

Addition of the normal distribution above to the generative process infuses flexibility to the Wishart, whose variance is usually defined by a single parameter (degrees of freedom^33^). In addition, the resulting matrix will have a diagonal scaled by δ_*k_i_*_ ^2^, hence allowing better fit to the empirical per-gene variances which are not captured directly by the regulatory model driving the prior for covariances. Off-diagonal elements are scaled by δ*_k_i__*δ*_k_j__*, a transformation which decouples the correlation structure embedded in the off-diagonal elements from the scaling of the diagonal. Specifically, correlations between genes in the original matrix Σ_*k*_′ are encoded as Σ′_*k,ij*_/σ′_*i*_σ′_*j*_. After scaling, δ’s in the numerator and denominator cancel, hence allowing the overall structure to be maintained under any arbitrary scaling of per-gene variances to fit the empirical data per cluster.

We note that with the above approximations, the constraint on the sign of *R_k_* is not always enforced to be the same as Σ′. Thus, we have more confidence in the inferred strength of regulation (magnitude of *R_k_*). The estimated regulatory strength is used to identify master regulators in **Fig. 3c** (as explained in section “master regulators” below). We also show the robustness of inferred regulatory strength in the section “robustness analysis” below.

###### Guide for choice of parameters

The variational inference implementation of Symphony requires choice of several hyperparameters. By default, priors on cluster mean expression are set with empirical means across the cells in that cluster as explained above, and shape and rate parameters for the Gamma prior on peak heights are set as 4.5 and 1 respectively for a relatively uninformative prior. Other parameters, particularly those controlling the variance of distributions in the generative model, are user-defined and should be tuned to each dataset.

As Symphony is designed to manage a trade-off between fitting to expression covariance and chromatin accessibility in the posterior distribution over GRNs, the choice of variance parameter on the prior distribution for each *R_k_* denoted by λ, as well as the degrees of freedom in Wishart linking *R_k_* to Σ_*k*_ denoted by γ, can be chosen to prioritize fit to each type of data. To inform the choice of these parameters, we recommend setting these parameters with small values and checking the empirical fit of the posterior to both data types. For example, the parameter settings used in this study (λ = 0.005, γ = *d* + 1 where *d* is the number of genes) ensured strong correlation of posterior GRNs with both the inferred peak heights, which in turn associated strongly with the bulk accessibility data, and further with the posterior covariance which itself associated with the empirical covariance. We also track these correlations over inference to ensure they increase over iterations. Details can be found in https://github.com/dpeerlab/Symphony.

###### ATAC-seq samples used in Symphony

Prior to running Symphony, T_EX_ and T_PEX_ clusters that fell in the same sort compartment of CD4 or CD8, CD45RA or CD45RO were grouped together as listed in Table S6 below. **Fig 3a** and **Extended Data Fig 4a** show ATAC-seq accessibility profiles for these samples (full list of samples and QC metrics are provided in **Suppl. Table 5**). Bigwig files were loaded to IGV^34^ to visualize normalized accessibility signal with differential accessibility identified with DESeq2^35^.

**Table S6.** Groups of ATAC-seq samples used for deconvolution of accessibility profiles in Symphony

| Symphony deconvolution group | Enriched exhaustion state | Enriched exhausted clusters | cell type | ATAC-seq sample ID |
| --- | --- | --- | --- | --- |
| 1 | T_EX | 14,27 | CD8 CD45RO | C149 |
| 1 | T_EX | 14,27 | CD8 CD45RO | C156 |
| 1 | T_EX | 14,27 | CD8 CD45RO | C45 |
| 1 | T_EX | 14,27 | CD8 CD45RO | C66 |
| 1 | T_EX | 14,27 | CD8 CD45RO | C75 |
| 2 | T_EX | M3 (4,7,3,22) | CD8 CD45RA | C70 |
| 2 | T_EX | M3 (4,7,3,22) | CD8 CD45RA | C171 |
| 2 | T_EX | M3 (4,7,3,22) | CD8 CD45RA | C167 |
| 2 | T_EX | M3 (4,7,3,22) | CD8 CD45RA | C144 |
| 2 | T_EX | M3 (4,7,3,22) | CD8 CD45RA | C139 |
| 2 | T_EX | M3 (4,7,3,22) | CD8 CD45RA | C106 |
| 3 | P_EX | M2 (5,11,23) | CD8 CD45RO | C39 |
| 3 | P_EX | M2 (5,11,23) | CD8 CD45RO | C36 |
| 3 | P_EX | M2 (5,11,23) | CD8 CD45RO | C156 |
| 3 | P_EX | M2 (5,11,23) | CD8 CD45RO | C164 |
| 3 | P_EX | M2 (5,11,23) | CD8 CD45RO | C168 |
| 4 | P_EX | M2 (5,11,23) | CD4 CD45RO | C37 |
| 4 | P_EX | M2 (5,11,23) | CD4 CD45RO | C34 |
| 4 | P_EX | M2 (5,11,23) | CD4 CD45RO | C127 |
| 4 | P_EX | M2 (5,11,23) | CD4 CD45RO | C158 |
| 4 | P_EX | M2 (5,11,23) | CD4 CD45RO | C162 |
| 5 | P_EX | M1 (19,28) | CD8 CD45RA | C41 |
| 5 | P_EX | M1 (19,28) | CD8 CD45RA | C79 |
| 5 | P_EX | M1 (19,28) | CD8 CD45RA | C118 |
| 5 | P_EX | M1 (19,28) | CD8 CD45RA | C35 |
| 6 | P_EX | M1 (19,28) | CD4 CD45RA | C76 |
| 6 | P_EX | M1 (19,28) | CD4 CD45RA | C33 |
| 6 | P_EX | M1 (19,28) | CD4 CD45RA | C116 |

In each group 1-6 listed in Table S6, scRNA-seq data and ATAC-seq data from the same samples are used as input to Symphony. Bulk ATAC-seq samples from different patients are assumed as biological replicates, and deconvolved using Symphony to achieve accessibility profiles for each cluster. Combined with scRNA-seq data for the clusters, Symphony infers a GRN for each cluster shown in **Fig. 3d** and **Extended Data Fig. 5**. We limited target genes to the pool of differentially expressed markers (**Suppl. Table 3**) across clusters. We filtered inferred regulatory links (entries of *R_k_*) that had a magnitude less than two (|*R_k_* |<2 selected based on knee-point of distribution, |CV|>0.5).

With this implementation, the runtime for Symphony was 1h 52m on group 4 containing 2593 cells and 1305 pooled DEGs and 5h 54m on group 1 with 7181 cells and 1459 DEGs, on a local machine with 64GB of RAM and 12 CPU cores (2.7 GHz processors). This runtime is at least 40 times faster than MCMC inference used in Biscuit which has a similar model structure.

###### Robustness analysis

To test the robustness of GRN inference, we performed a leave one (patient) out analysis in the T_EX_ CD8 and T_PEX_ CD4 groups. Specifically, we fit Symphony to scRNA-seq and ATAC-seq data for each group and excluded ATAC-seq data from one patient at a time. We then compared the coefficient of variation (CV) of predicted regulatory links across the leave-one-out iterations to the inferred regulation from the entire data. As shown below in **Fig. S10** and **Fig. S11**, CV is lower for stronger regulatory links and the majority of links have CV<1.

**Figure S10.**
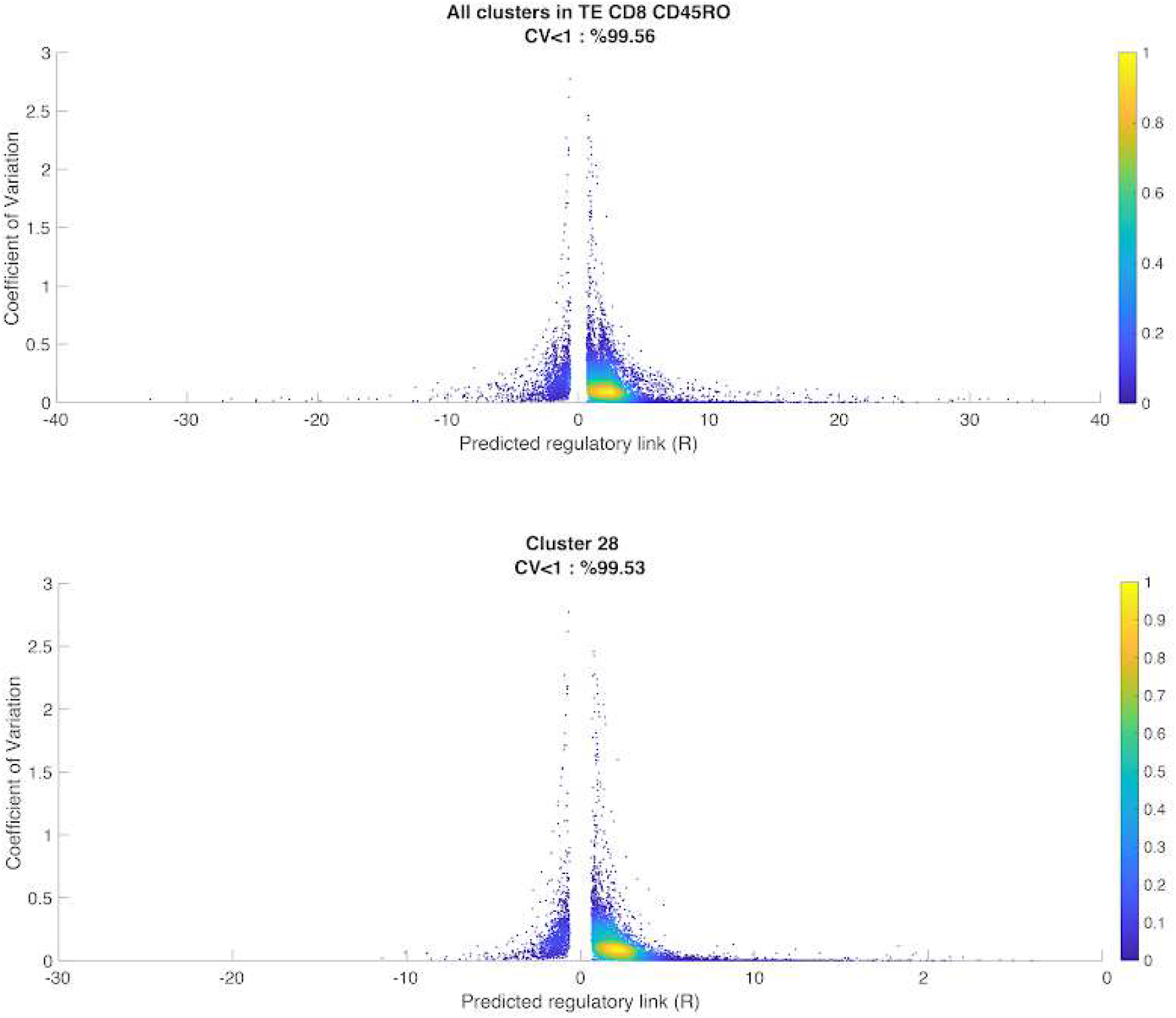
Absolute value of coefficient of variation (|CV| in y-axis) of regulation across leave-one-out analyses vs. inferred regulation from the full dataset (x-axis); each dot is a regulatory link in the network colored by density of data points; top row corresponds to all clusters in in group 1 (T_EX_) and bottom row corresponds to cluster 28 (deconvolved from the group).

**Figure S11.**
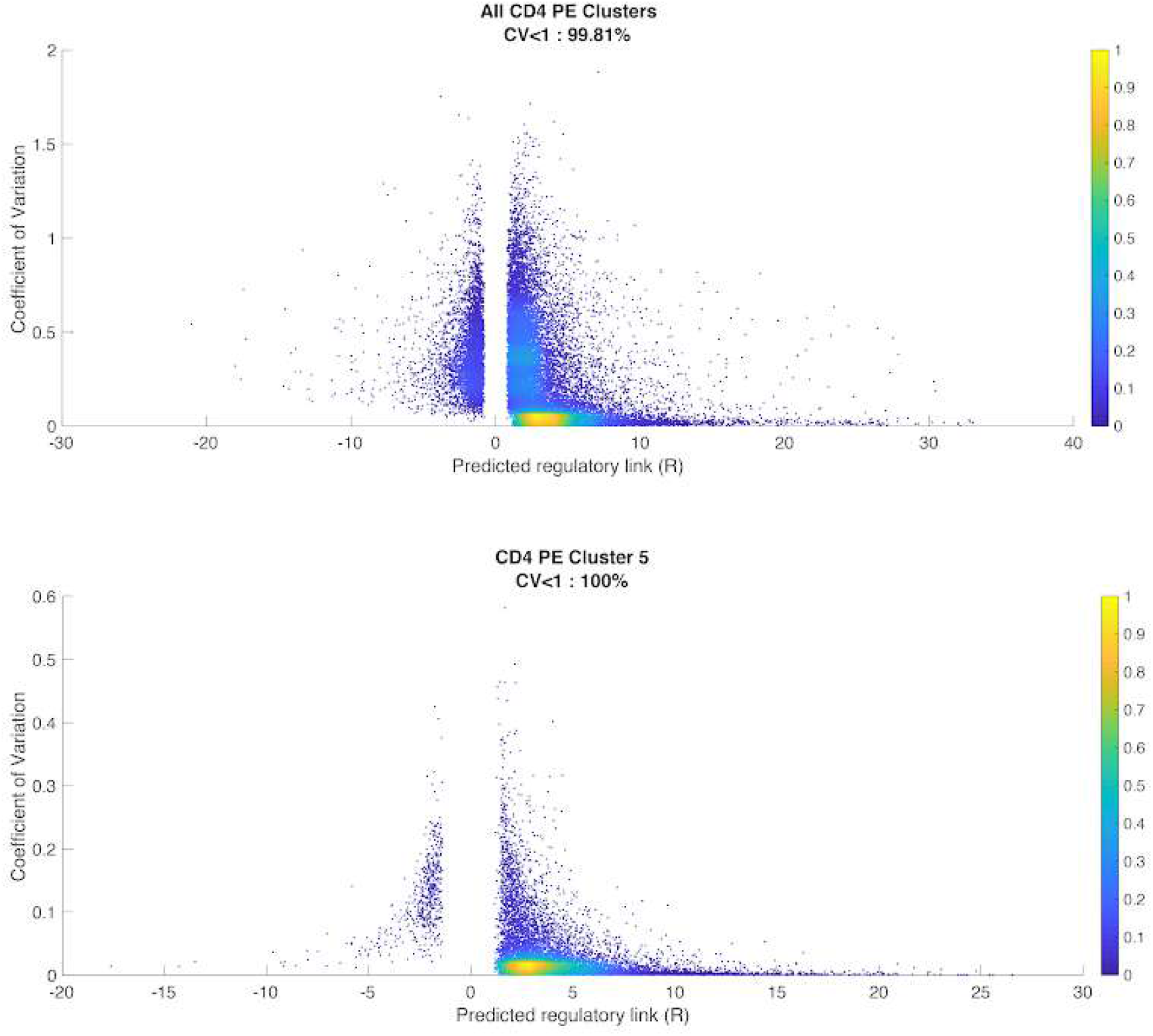
Similar to Figure S10; top row corresponds to inferred links from all clusters in group 4 (T_PEX_) and bottom row corresponds to cluster 5 (deconvolved from the group).

###### Master regulators

We used the output GRNs from Symphony to identify master regulators of each cluster as follows: For cluster *k*, we averaged the inferred impact of each TF *a*, across all targets *b* that are differentially expressed genes (DEGs) in the cluster (listed in **Suppl. Table 3**): 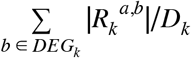 with *D_k_* being the number of DEGs for cluster *k*. The resulting average regulatory strength of each TF in each cluster is shown in **Fig. 3c**. We performed a one-sided t-test between T_EX_ clusters and all other exhausted clusters to find “differential regulators” of T_EX_ clusters shown with dotted line box in **Fig. 3c**, and green nodes in **Fig. 3d** and **Extended data Fig. 5**. Similarly, we identified differential regulators of T_PEX_ M1 and M2 subsets (**Fig. 3c**) shown as pink nodes in **Fig. 3d** and **Extended data Fig. 5**.

###### Regulatory network

To elucidate the target genes impacted most by these master regulators, we filtered the GRNs by centrality or out-degree of regulators (defined as number of target genes predicted to be regulated by the TF) as well as regulatory strength (|*R_k_* |>2). **Fig. 3d** and **Extended data Fig. 5** show these subnetworks containing individual known^36,37^ and novel links. The circuitry for exhausted clusters reveals similarity and differences in network architecture across clusters. We identified mediating regulators such as *BCL6* connecting two other regulators (*TBPL1* and *E2F2*) differentially regulating cluster 27. The network link predictions are supported by co-expression and/or accessibility (**Fig. 3d**). Other predicted repressors such as *TCF7L2* are supported by mutually exclusive (negative) co-expression patterns with DEGs.

#### Supplementary Note 4: Analysis of paired single-cell TCR- and RNA-seq

##### Preprocessing and identification of exhausted clusters

Single cell 5’ RNA-seq reads were processed with the Cell Ranger pipeline available from 10x Genomics. QC metrics for this data is provided in **Suppl. Table S6**. A total of 23K total T cells were identified based on {CD3D, CD3E} expression (similar to **Suppl. Note 2**) and normalized and clustered using Biscuit with the same parameters as in **Suppl. Note 2** (shown in **Fig. S12**).

**Figure S12.**
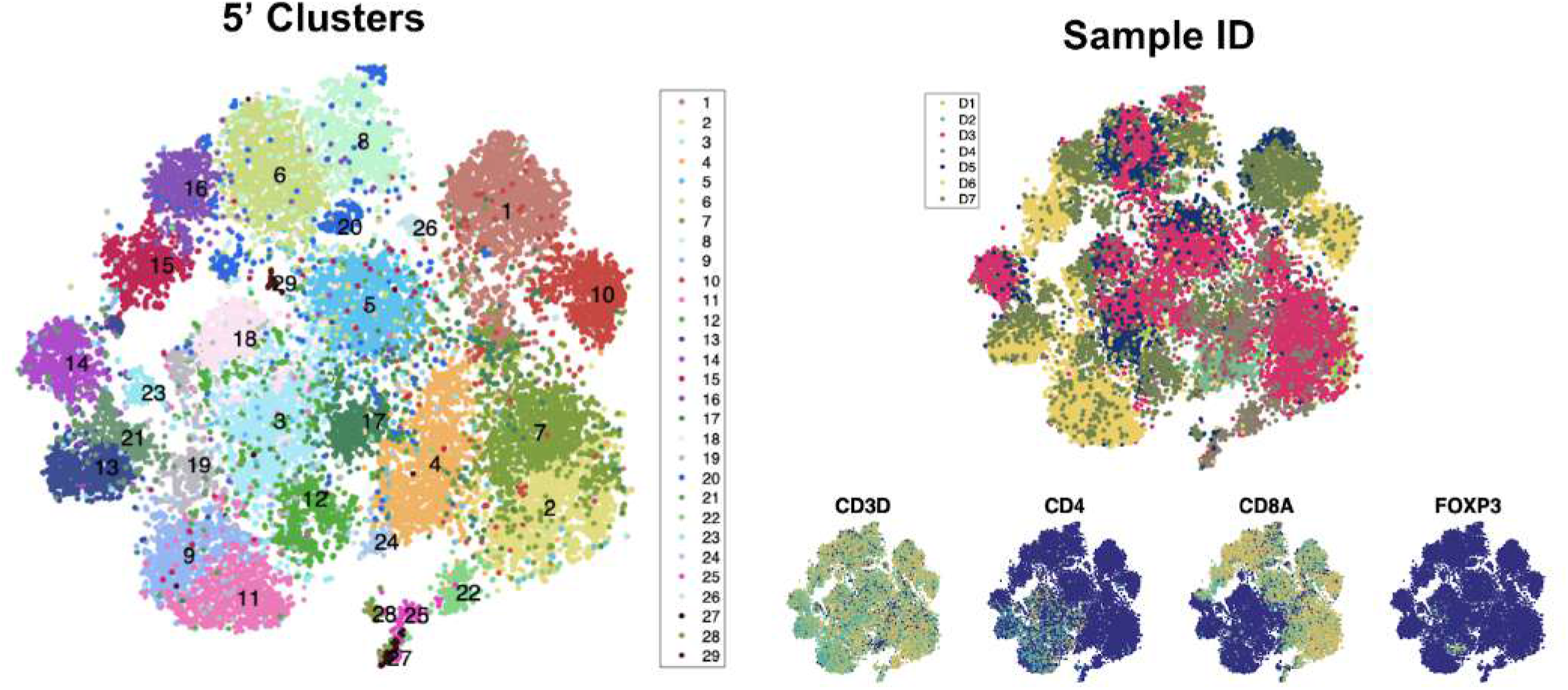
t-SNE projection of normalized 5’ scRNA-seq data for all T cells from two Rs (listed in Suppl. Table 6), each dot represents a cell colored by cluster (left), sample ID (top right), and markers (bottom right).

The 29 newly identified clusters (Fig. S12) were scored for the same T_PEX_ and T_EX_ signatures (listed in **Extended Data Table 3**), and the clusters with the highest scores were identified as T_PEX_ and T_EX_ clusters (**Fig. S13**).

**Figure S13.**
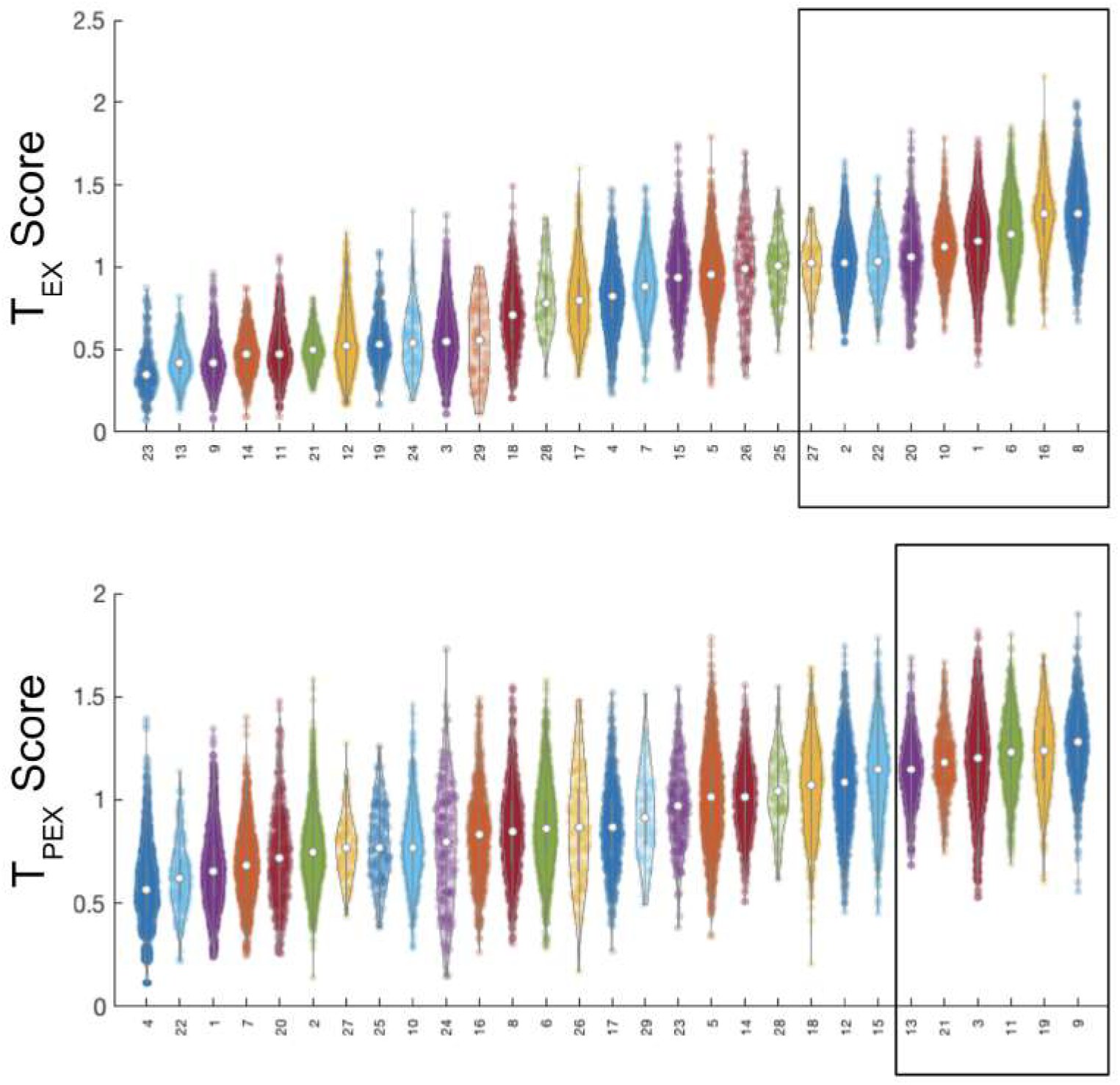
Violin plots showing density of T cell clusters from 5’ scRNA-seq data along T_EX_ (top) or T_PEX_ (bottom) viral gene set scores; x-axis shows cluster IDs ordered by median score; clusters identified as exhausted are marked with boxes.

##### Preprocessing and analysis of TCR clonotypes

Single cell TCR-seq reads were aligned to the GRCh^38^ reference genome and consensus TCR annotation was performed using Cell Ranger V(D)J (10x Genomics, version 2.1.0.). QC metrics are provided in **Suppl. Table 6**.

Clonotypes mapping to *TRB* loci were used to annotate each cell, similar to others^38^. Overlap between clonotypes from T_EX_ cells and T_PEX_ cells (**Fig. 4a**) was measured by counting the number of cells from each group per clonotype and performing a hypergeometric test using the `phyper` function with R. Venn diagrams were drawn using the *eulerr* package.

TCR diversity (**Fig. 4b**) was calculated between all RNA clusters on a per patient basis via Gini coefficient^39^ using the `ineq ()` function within the *ineq* package.

To determine the kinetics of T_EX_ and T_PEX_ clonotypes after DLI (**Fig, 4c,d**), the proportion of pre- and post-treatment cells were calculated for both patients together. Clonotypes were defined as expanding if they significantly enriched pre-DLI (p<0.05 according to Fisher’s exact test), contracting if they were enriched post-DLI (p<0.05 by Fisher’s exact test), and persistent otherwise. Viral-specific clonotypes were identified via VDJdb^40^ and marked (V). Statistical analysis was performed in R version 3.5.3. Plots were generated using the *ggplot* package.

## References

1. Yofe, I., Dahan, R. & Amit, I. Single-cell genomic approaches for developing the next generation of immunotherapies. Nat. Med. 26, 171–177 (2020).

2. Lesterhuis, W. J. et al. Dynamic versus static biomarkers in cancer immune checkpoint blockade: unravelling complexity. Nature Reviews Drug Discovery vol. 16 264–272 (2017).

3. Gubin, M. M. et al. High-Dimensional Analysis Delineates Myeloid and Lymphoid Compartment Remodeling during Successful Immune-Checkpoint Cancer Therapy. Cell 175, 1443 (2018).

4. Jenq, R. R. & van den Brink, M. R. M. Allogeneic haematopoietic stem cell transplantation: individualized stem cell and immune therapy of cancer. Nat. Rev. Cancer 10, 213–221 (2010).

5. Bachireddy, P. & Wu, C. J. Understanding anti-leukemia responses to donor lymphocyte infusion. Oncoimmunology 3, e28187 (2014).

6. Collins, R. H., Jr et al. Donor leukocyte infusions in 140 patients with relapsed malignancy after allogeneic bone marrow transplantation. J. Clin. Oncol. 15, 433–444 (1997).

7. Kolb, H. J. et al. Graft-versus-leukemia effect of donor lymphocyte transfusions in marrow grafted patients. Blood 86, 2041–2050 (1995).

8. Champlin, R. et al. Retention of graft-versus-leukemia using selective depletion of CD8-positive T lymphocytes for prevention of graft-versus-host disease following bone marrow transplantation for chronic myelogenous leukemia. Transplant. Proc. 23, 1695–1696 (1991).

9. Giralt, S. et al. CD8-depleted donor lymphocyte infusion as treatment for relapsed chronic myelogenous leukemia after allogeneic bone marrow transplantation. Blood 86, 4337–4343 (1995).

10. Alyea, E. P. et al. Toxicity and efficacy of defined doses of CD4(+) donor lymphocytes for treatment of relapse after allogeneic bone marrow transplant. Blood 91, 3671–3680 (1998).

11. Soiffer, R. J. et al. Randomized trial of CD8 T-cell depletion in the prevention of graft-versus-host disease associated with donor lymphocyte infusion. Biology of Blood and Marrow Transplantation vol. 8 625–632 (2002).

12. Claret, E. J. et al. Characterization of T cell repertoire in patients with graft-versus-leukemia after donor lymphocyte infusion. J. Clin. Invest. 100, 855–866 (1997).

13. Zhang, W. et al. Graft-versus-leukemia antigen CML66 elicits coordinated B-cell and T-cell immunity after donor lymphocyte infusion. Clin. Cancer Res. 16, 2729–2739 (2010).

14. Bachireddy, P. et al. Reversal of in situ T-cell exhaustion during effective human antileukemia responses to donor lymphocyte infusion. Blood 123, 1412–1421 (2014).

15. Liu, L. et al. Reversal of T Cell Exhaustion by the First Donor Lymphocyte Infusion Is Associated with the Persistently Effective Antileukemic Responses in Patients with Relapsed AML after Allo-HSCT. Biol. Blood Marrow Transplant. 24, 1350–1359 (2018).

16. Porter, D. L., Roth, M. S., McGarigle, C., Ferrara, J. & Antin, J. H. Induction of Graft-versus-Host Disease as Immunotherapy for Relapsed Chronic Myeloid Leukemia. New England Journal of Medicine vol. 330 100–106 (1994).

17. Levine, J. H. et al. Data-Driven Phenotypic Dissection of AML Reveals Progenitor-like Cells that Correlate with Prognosis. Cell 162, 184–197 (2015).

18. Prabhakaran, S., Azizi, E., Carr, A. & Pe’er, D. Dirichlet Process Mixture Model for Correcting Technical Variation in Single-Cell Gene Expression Data. in International Conference on Machine Learning 1070–1079 (2016).

19. Azizi, E. et al. Single-Cell Map of Diverse Immune Phenotypes in the Breast Tumor Microenvironment. Cell 174, 1293–1308.e36 (2018).

20. Zientek, L. R. Exploratory and Confirmatory Factor Analysis: Understanding Concepts and Applications. Struct. Equ. Modeling 15, 729–734 (2008).

21. Li, H. et al. Dysfunctional CD8 T Cells Form a Proliferative, Dynamically Regulated Compartment within Human Melanoma. Cell vol. 176 775–789.e18 (2019).

22. Singer, M. et al. A Distinct Gene Module for Dysfunction Uncoupled from Activation in Tumor-Infiltrating T Cells. Cell 171, 1221–1223 (2017).

23. Kallies, A., Zehn, D. & Utzschneider, D. T Precursor exhausted T cells: key to successful immunotherapy? Nat. Rev. Immunol. 20, 128–136 (2020).

24. Miller, B. C. et al. Subsets of exhausted CD8 T cells differentially mediate tumor control and respond to checkpoint blockade. Nat. Immunol. 20, 326–336 (2019).

25. Wu, T. et al. The TCF1-Bcl6 axis counteracts type I interferon to repress exhaustion and maintain T cell stemness. Sci Immunol 1, (2016).

26. Alfei, F. et al. TOX reinforces the phenotype and longevity of exhausted T cells in chronic viral infection. Nature 571, 265–269 (2019).

27. Scott, A. C. et al. TOX is a critical regulator of tumour-specific T cell differentiation. Nature 571, 270–274 (2019).

28. Khan, O. et al. TOX transcriptionally and epigenetically programs CD8 T cell exhaustion. Nature 571, 211–218 (2019).

29. Leong, Y. A. et al. CXCR5(+) follicular cytotoxic T cells control viral infection in B cell follicles. Nat. Immunol. 17, 1187–1196 (2016).

30. Brummelman, J. et al. High-dimensional single cell analysis identifies stem-like cytotoxic CD8 T cells infiltrating human tumors. J. Exp. Med. 215, 2520–2535 (2018).

31. Im, S. J. et al. Defining CD8+ T cells that provide the proliferative burst after PD-1 therapy. Nature 537, 417–421 (2016).

32. Youngblood, B. et al. Effector CD8 T cells dedifferentiate into long-lived memory cells. Nature 552, 404–409 (2017).

33. Akondy, R. S. et al. Origin and differentiation of human memory CD8 T cells after vaccination. Nature 552, 362–367 (2017).

34. Sen, D. R. et al. The epigenetic landscape of T cell exhaustion. Science 354, 1165–1169 (2016).

35. Pauken, K. E. et al. Epigenetic stability of exhausted T cells limits durability of reinvigoration by PD-1 blockade. Science 354, 1160–1165 (2016).

36. C. Burdziak, E. Azizi, S. Prabhakaran, D. Pe’er. A Nonparametric Multi-view Model for Estimating Cell Type-Specific Gene Regulatory Networks. arXiv (2019).

37. Utzschneider, D. T et al. T Cell Factor 1-Expressing Memory-like CD8(+) T Cells Sustain the Immune Response to Chronic Viral Infections. Immunity 45, 415–427 (2016).

38. Paley, M. A. et al. Progenitor and terminal subsets of CD8+ T cells cooperate to contain chronic viral infection. Science 338, 1220–1225 (2012).

39. Chen, Z. et al. TCF-1-Centered Transcriptional Network Drives an Effector versus Exhausted CD8 T Cell-Fate Decision. Immunity vol. 51 840–855.e5 (2019).

40. Link, C. S. et al. Abundant cytomegalovirus (CMV) reactive clonotypes in the CD8(+) T cell receptor alpha repertoire following allogeneic transplantation. Clin. Exp. Immunol. 184, 389–402 (2016).

41. Ricordel, C., Friboulet, L., Facchinetti, F. & Soria, J.-C. Molecular mechanisms of acquired resistance to third-generation EGFR-TKIs in EGFR T790M-mutant lung cancer. Ann. Oncol. 30, 858 (2019).

42. Goetz, E. M. & Garraway, L. A. Mechanisms of Resistance to Mitogen-Activated Protein Kinase Pathway Inhibition in BRAF-Mutant Melanoma. American Society of Clinical Oncology Educational Book 680–684 (2012) doi:10.14694/edbook_am.2012.32.189.

43. Sade-Feldman, M. et al. Defining T Cell States Associated with Response to Checkpoint Immunotherapy in Melanoma. Cell 176, 404 (2019).

44. Olson, E. M., Lin, N. U., Krop, I. E. & Winer, E. P. The ethical use of mandatory research biopsies. Nat. Rev. Clin. Oncol. 8, 620–625 (2011).

45. TRACERx Renal consortium. TRACERx Renal: tracking renal cancer evolution through therapy. Nat. Rev. Urol. 14, 575–576 (2017).

46. Baitsch, L. et al. Exhaustion of tumor-specific CD8+ T cells in metastases from melanoma patients. J. Clin. Invest. 121, 2350–2360 (2011).

47. Bengsch, B. et al. Epigenomic-Guided Mass Cytometry Profiling Reveals Disease-Specific Features of Exhausted CD8 T Cells. Immunity 48, 1029–1045.e5 (2018).

48. He, R. et al. Follicular CXCR5-expressing CD8 T cells curtail chronic viral infection. Nature vol. 537 412–416 (2016).

49. Siddiqui, I. et al. Intratumoral Tcf1PD-1CD8 T Cells with Stem-like Properties Promote Tumor Control in Response to Vaccination and Checkpoint Blockade Immunotherapy. Immunity 50, 195–211.e10 (2019).

50. Yost, K. E. et al. Clonal replacement of tumor-specific T cells following PD-1 blockade. Nat. Med. 25, 1251–1259 (2019).

51. Aubert, R. D. et al. Antigen-specific CD4 T-cell help rescues exhausted CD8 T cells during chronic viral infection. Proc. Natl. Acad. Sci. U. S. A. 108, 21182–21187 (2011).

52. Zander, R. et al. CD4 T Cell Help Is Required for the Formation of a Cytolytic CD8 T Cell Subset that Protects against Chronic Infection and Cancer. Immunity vol. 51 1028–1042.e4 (2019).

53. Bingham, Eli and Chen, Jonathan P and Jankowiak, Martin and Obermeyer, Fritz and Pradhan, Neeraj and Karaletsos, Theofanis and Singh, Rohit and Szerlip, Paul and Horsfall, Paul and Goodman, Noah D. Pyro: deep universal probabilistic programming. The Journal of Machine Learning Research 20, 973–978 (2019).

54. Tran, D. et al. Edward: A library for probabilistic modeling, inference, and criticism. arXiv [stat.CO] (2016).

55. Coifman, R., Coppi, A., Hirn, M. & Warner, F. Diffusion Geometry Based Nonlinear Methods for Hyperspectral Change Detection. http://dx.doi.org/10.21236/ada524546 (2010) doi:10.21236/ada524546.

## References

1. Alyea, E. P. et al. Toxicity and efficacy of defined doses of CD4(+) donor lymphocytes for treatment of relapse after allogeneic bone marrow transplant. Blood 91, 3671–3680 (1998).

2. Corces, M. R. et al. Lineage-specific and single-cell chromatin accessibility charts human hematopoiesis and leukemia evolution. Nat. Genet. 48, 1193–1203 (2016).

3. Azizi, E. et al. Single-Cell Map of Diverse Immune Phenotypes in the Breast Tumor Microenvironment. Cell 174, 1293–1308.e36 (2018).

4. Levine, J. H. et al. Data-Driven Phenotypic Dissection of AML Reveals Progenitor-like Cells that Correlate with Prognosis. Cell 162, 184–197 (2015).

5. Prabhakaran, S., Azizi, E., Carr, A. & Pe’er, D. Dirichlet Process Mixture Model for Correcting Technical Variation in Single-Cell Gene Expression Data. in International Conference on Machine Learning 1070–1079 (2016).

6. Blackinton, J. G. & Keene, J. D. Functional coordination and HuR-mediated regulation of mRNA stability during T cell activation. Nucleic Acids Res. 44, 426–436 (2016).

7. Singer, M. et al. A Distinct Gene Module for Dysfunction Uncoupled from Activation in Tumor-Infiltrating T Cells. Cell 166, 1500–1511.e9 (2016).

8. Cheadle, C. Stability Regulation of mRNA and the Control of Gene Expression. Ann. N. Y. Acad. Sci. 1058, 196–204 (2005).

9. Marrack, P. et al. Homeostasis of αβ TCR T cells. Nat. Immunol. 1, 107–111 (2000).

10. Maaten, L. van der & Hinton, G. Visualizing Data using t-SNE. J. Mach. Learn. Res. 9, 2579–2605 (2008).

11. Amir, E.-A. D. et al. viSNE enables visualization of high dimensional single-cell data and reveals phenotypic heterogeneity of leukemia. Nat. Biotechnol. 31, 545–552 (2013).

12. Im, S. J. et al. Defining CD8+ T cells that provide the proliferative burst after PD-1 therapy. Nature 537, 417–421 (2016).

13. Gattinoni, L., Speiser, D. E., Lichterfeld, M. & Bonini, C. T memory stem cells in health and disease. Nature Medicine vol. 23 18–27 (2017).

14. Oberdoerffer, S. et al. Regulation of CD45 alternative splicing by heterogeneous ribonucleoprotein, hnRNPLL. Science 321, 686–691 (2008).

15. Li, H. et al. Dysfunctional CD8 T Cells Form a Proliferative, Dynamically Regulated Compartment within Human Melanoma. Cell vol. 176 775–789.e18 (2019).

16. Singer, M. et al. A Distinct Gene Module for Dysfunction Uncoupled from Activation in Tumor-Infiltrating T Cells. Cell 171, 1221–1223 (2017).

17. Tao, T. & Vu, V. On random ±1 matrices: Singularity and determinant. Random Struct. Algorithms 28, 1–23 (2005).

18. Coifman, R., Coppi, A., Hirn, M. & Warner, F. Diffusion Geometry Based Nonlinear Methods for Hyperspectral Change Detection. http://dx.doi.org/10.21236/ada524546 (2010) doi:10.21236/ada524546.

19. Setty, M. et al. Wishbone identifies bifurcating developmental trajectories from single-cell data. Nat. Biotechnol. 34, 637–645 (2016).

20. Bhattacharyya, A. On a Geometrical Representation of Probability Distributions and its use in Statistical Inference. Calcutta Statistical Association Bulletin vol. 40 23–49 (1990).

21. Ahmed, H. I., Herrera, M., Liew, Y. J. & Aranda, M. Long-Term Temperature Stress in the Coral Model Aiptasia Supports the ‘Anna Karenina Principle’ for Bacterial Microbiomes. Frontiers Microbiology vol. 10 (2019).

22. Vert, J.-P. Kernel Methods in Genomics and Computational Biology. Kernel Methods)n Bioengineering, Signal and Image Processing 42–63 doi:10.4018/978-1-59904-042-4.ch002.

23. Bingham, Eli and Chen, Jonathan P and Jankowiak, Martin and Obermeyer, Fritz and Pradhan, Neeraj and Karaletsos, Theofanis and Singh, Rohit and Szerlip, Paul and Horsfall, Paul and Goodman, Noah D. Pyro: deep universal probabilistic programming. The Journal of Machine Learning Research 20, 973–978 (2019).

24. Blei, D. M., Kucukelbir, A. & McAuliffe, J. D. Variational Inference: A Review for Statisticians. Journal of the American Statistical Association vol. 112 859–877 (2017).

25. Langmead, B. & Salzberg, S. L. Fast gapped-read alignment with Bowtie 2. Nat. Methods 9, 357–359 (2012).

26. Zhang, Y. et al. Model-based analysis of ChIP-Seq (MACS). Genome Biol. 9, R137 (2008).

27. Reske, J. J., Wilson, M. R. & Chandler, R. L. ATAC-seq normalization method can significantly affect differential accessibility analysis and interpretation. Epigenetics Chromatin 13, 22 (2020).

28. C. Burdziak, E. Azizi, S. Prabhakaran, D. Pe’er. A Nonparametric Multi-view Model for Estimating Cell Type-Specific Gene Regulatory Networks. arXiv (2019).

29. Buenrostro, J. D., Wu, B., Chang, H. Y. & Greenleaf, W. J. ATAC-seq: A Method for Assaying Chromatin Accessibility Genome-Wide. Curr. Protoc. Mol. Biol. 109, 21.29.1–21.29.9 (2015).

30. Bailey, T. L. et al. MEME SUITE: tools for motif discovery and searching. Nucleic Acids Res. 37, W202–8 (2009).

31. Tran, D. et al. Edward: A library for probabilistic modeling, inference, and criticism. arXiv [stat.CO] (2016).

32. Kshirsagar, A. M. Bartlett Decomposition and Wishart Distribution. The Annals of Mathematical Statistics vol. 30 239–241 (1959).

33. Alvarez, I., Niemi, J. & Simpson, M. BAYESIAN INFERENCE FOR A COVARIANCE MATRIX. Conference on Applied Statistics in Agriculture (2014) doi:10.4148/2475-7772.1004.

34. Robinson, J. T. et al. Integrative genomics viewer. Nat. Biotechnol. 29, 24–26 (2011).

35. Love, M. I., Huber, W & Anders, S. Moderated estimation of fold change and dispersion for RNA-seq data with DESeq2. Genome Biol. 15, 550 (2014).

36. Man, K. et al. Transcription Factor IRF4 Promotes CD8 T Cell Exhaustion and Limits the Development of Memory-like T Cells during Chronic Infection. Immunity 47, 1129–1141.e5 (2017).

37. Wu, T. et al. The TCF1-Bcl6 axis counteracts type I interferon to repress exhaustion and maintain T cell stemness. Sci Immunol 1, (2016).

38. Yost, K. E. et al. Clonal replacement of tumor-specific T cells following PD-1 blockade. Nat. Med. 25, 1251–1259 (2019).

39. Dixon, P. M., Weiner, J., Mitchell-Olds, T. & Woodley, R. Bootstrapping the Gini Coefficient of Inequality. Ecology vol. 68 1548–1551 (1987).

40. Bagaev, D. V. et al. VDJdb in 2019: database extension, new analysis infrastructure and a T-cell receptor motif compendium. Nucleic Acids Research vol. 48 D1057–D1062 (2020).

